# Population history from the Neolithic to present on the Mediterranean island of Sardinia: An ancient DNA perspective

**DOI:** 10.1101/583104

**Authors:** Joseph H. Marcus, Cosimo Posth, Harald Ringbauer, Luca Lai, Robin Skeates, Carlo Sidore, Jessica Beckett, Anja Furtwängler, Anna Olivieri, Charleston Chiang, Hussein Al-Asadi, Kushal Dey, Tyler A. Joseph, Clio Der Sarkissian, Rita Radzevičiūtė, Maria Giuseppina Gradoli, Wolfgang Haak, David Reich, David Schlessinger, Francesco Cucca, Johannes Krause, John Novembre

**Affiliations:** Department of Human Genetics, University of Chicago, Chicago, IL, USA.; Max Planck Institute for the Science of Human History, Jena, Germany.; Institute for Archaeological Sciences, University of Tübingen, Tübingen, Germany.; Department of Anthropology, University of South Florida, Tampa, FL, USA.; Department of Archaeology, Durham University, Durham, United Kingdom.; Istituto di Ricerca Genetica e Biomedica - CNR, Cagliari, Italy.; Dipartimento di Scienze Biomediche, Università di Sassari, Sassari, Italy.; Private contractor, Cagliari, Sardinia.; Dipartimento di Biologia e Biotecnologie “L. Spallanzani”, Università di Pavia, Pavia, Italy.; Center for Genetic Epidemiology, Department of Preventive Medicine, Keck School of Medicine, University of Southern California, Los Angeles, CA, USA.; Department of Statistics, University of Chicago, Chicago, IL, USA.; Committee on Evolutionary Biology, University of Chicago, Chicago, IL, USA.; Department of Epidemiology, Harvard School of Public Health, Boston, Massachusetts 02115, USA.; Department of Computer Science, Columbia University, New York, NY, USA.; Team AGES Laboratory AMIS Faculté de Médecine de Purpan, Toulouse, France.; School of Archaeology and Ancient History, University of Leicester, Leicester, United Kingdom.; Department of Genetics, Harvard Medical School, Boston, Massachusetts 02115, USA.; Broad Institute of Harvard and MIT, Cambridge, MA, USA.; Howard Hughes Medical Institute, Harvard Medical School, Boston, MA, USA.; Max Planck-Harvard Research Center for the Archaeoscience of the Ancient Mediterranean; Laboratory of Genetics, NIA, NIH, Baltimore, MD, USA.; Department of Ecology and Evolution, University of Chicago, Chicago, IL, USA.

## Abstract

Recent ancient DNA studies of western Eurasia have revealed a dynamic history of admixture, with evidence for major migrations during the Neolithic and Bronze Age. The population of the Mediterranean island of Sardinia has been notable in these studies – Neolithic individuals from mainland Europe cluster more closely with Sardinian individuals than with all other present-day Europeans. The current model to explain this result is that Sardinia received an initial influx of Neolithic ancestry and then remained relatively isolated from expansions in the later Neolithic and Bronze Age that took place in continental Europe. To test this model, we generated genome-wide capture data (approximately 1.2 million variants) for 43 ancient Sardinian individuals spanning the Neolithic through the Bronze Age, including individuals from Sardinia’s Nuragic culture, which is known for the construction of numerous large stone towers throughout the island. We analyze these new samples in the context of previously generated genome-wide ancient DNA data from 972 ancient individuals across western Eurasia and whole-genome sequence data from approximately 1,500 modern individuals from Sardinia. The ancient Sardinian individuals show a strong affinity to western Mediterranean Neolithic populations and we infer a high degree of genetic continuity on the island from the Neolithic (around fifth millennium BCE) through the Nuragic period (second millennium BCE). In particular, during the Bronze Age in Sardinia, we do not find significant levels of the “Steppe” ancestry that was spreading in many other parts of Europe at that time. We also characterize subsequent genetic influx between the Nuragic period and the present. We detect novel, modest signals of admixture between 1,000 BCE and present-day, from ancestry sources in the eastern and northern Mediterranean. Within Sardinia, we confirm that populations from the more geographically isolated mountainous provinces have experienced elevated levels of genetic drift and that northern and southwestern regions of the island received more gene flow from outside Sardinia. Overall, our genetic analysis sheds new light on the origin of Neolithic settlement on Sardinia, reinforces models of genetic continuity on the island, and provides enhanced power to detect post-Bronze-Age gene flow. Together, these findings offer a refined demographic model for future medical genetic studies in Sardinia.

## Introduction

The whole-genome sequencing of Ötzi, a Neolithic individual who was preserved in ice for over 5,000 years near the Italo-Austrian border, revealed a surprisingly high level of shared ancestry with present-day Sardinian individuals (Ermini *et al.*, 2008; Keller *et al.*, 2012; Sikora *et al.*, 2014). Subsequent work on genome-wide variation in ancient Europeans expanded upon this observation, finding that many “early European farmer” individuals have their highest genetic affinity with present-day Sardinian individuals, even when from geographically distant locales (e.g. from Hungary, Germany, Spain, Sweden) (e.g. Skoglund *et al.*, 2012, 2014).

Accumulating ancient DNA (aDNA) results have provided a potential framework for understanding how early European farmers, such as Ö tzi, show such genetic affinity to modern Sardinians. In this framework, Europe was first inhabited by Paleolithic hunter-gatherer groups. Then, starting about 7,000 BCE, farming peoples arrived from the Middle East as part of a Neolithic transition (Ammerman and Cavalli-Sforza, 2014; Lazaridis *et al.*, 2014), spreading through Anatolia and the Balkans (Hofmanová *et al.*, 2016; Mathieson *et al.*, 2018) while progressively admixing with local hunter-gatherers (Lipson *et al.*, 2017). Subsequently, major movements from the Eurasian Steppe, beginning about 3,000 BCE, resulted in further admixture throughout Europe (Allentoft *et al.*, 2015; Haak *et al.*, 2015; Olalde *et al.*, 2018, 2019). These events are typically modeled in terms of three ancestry components, hunter-gatherers (and more specifically western hunter gatherers, “WHG”), early European farmers (“EEF”), and Steppe pastoralists (“Steppe”). Within this broad framework, the island of Sardinia is thought to have received a high level of EEF ancestry early on in its history and subsequently remained relatively isolated from the admixture occurring on mainland Europe (Keller *et al.*, 2012; Sikora *et al.*, 2014). However, this specific model for Sardinian population history has not been tested with genome-wide aDNA samples from the island.

The oldest known human remains on Sardinia have been dated to be ∼ 20, 000 years old (Melis, 2002), implying that humans first reached the island during the Paleolithic Age. Archaeological evidence suggests that the island was not densely populated in the Mesolithic, with only irregular and episodic settlements, mostly concentrated near the coast (Lugliè, 2018). The archaeological record shows that a population expansion coincided with a Neolithic transition in the sixth millennium BCE (Francalacci *et al.*, 2013). At the same time, the early Neolithic “Cardial Impressed Ware” culture was spreading across the western Mediterranean (Barnett, 2000), with radio-carbon dates indicating a rapid maritime expansion about 5,500 BCE (Zilhão, 2001; Martins *et al.*, 2015). Obsidian originating from Sardinia is found throughout many western Mediterranean archaeological sites associated with the middle Neolithic (Tykot, 1996), indicating that the island was integrated into a maritime trade network. In the middle Bronze Age, about 1,600 BCE, the “Nuragic” culture emerged that derives its name from thousands of distinctive stone towers, Nuraghi, constructed across Sardinia’s landscape and in many instances still well preserved. More recently, the archaeological and historical record shows the influence of several major Mediterranean groups, such as Phoenicians, Carthaginians, the Roman and Byzantine empires, and later with North Africa, Tuscany, Genoa, Catalonia, Spain, Southern France, and Piedmont (Ortu, 2011; Mastino, 2005).

The population genetics of Sardinia has long been studied (e.g. see Calò *et al.*, 2008) in part because of its importance as a population for medical genetics (Lettre and Hirschhorn, 2015). Pioneering studies, using classical genetic loci such as G6PD, HBB, and HLA and later maps of linkage disequilibrium, revealed that Sardinia is a genetic isolate with heterogeneous population sub-structure (e.g. Siniscalco *et al.*, 1966; Contu *et al.*, 1992; Barbujani and Sokal, 1990; Eaves *et al.*, 2000; Zavattari *et al.*, 2000; Cavalli-Sforza, 2005; Sidore *et al.*, 2015). Recently, Chiang *et al.* (2018) analyzed the whole genome sequences of 3,514 individuals from Sardinia to investigate the population genomic history of the island in finer resolution. In line with previous studies, they found substructure in which the mountainous Ogliastra region of central/eastern Sardinia carries a signature of relative isolation, presumably due to restricted gene flow across the rugged terrain. They also used a small sample of continental European aDNA to show suggestive evidence for differential contributions of ancestry from WHG, EEF, and Steppe to Sardinian genetic variation. This initial observation and the increased resolution of temporal, geographic, and cultural sampling in aDNA prompted us to investigate aDNA from Sardinia to gain further understanding.

Four previous studies have analyzed aDNA to provide an initial view of the genetics of pre-historic Sardinia, in each case, using mitochondrial DNA. Ghirotto *et al.* (2009) contrasted patterns of continuity between Ogliastra (the mountainous and historically isolated central region) and Gallura (a region in northern Sardinia with cultural and linguistic connections to Corsica), finding evidence for more genetic turnover in Gallura. Modi *et al.* (2017) provided the first complete mitogenomes of two Mesolithic individuals and found support for a model in which Mesolithic ancestry on the island was replaced by incoming populations in the Neolithic. Olivieri *et al.* (2017), in a companion project to the work described here, analyzed 21 ancient mitogenomes from Sardina as well as 3,491 mitogenomes from contemporary Sardinians and estimated the coalescent times of Sardinian-specific mtDNA haplogroups finding support for most of them originating in the Neolithic or later, but with a few coalescing earlier. Finally, Matisoo-Smith *et al.* (2018) analyzed mitogenomes in a Phoenician colony on Sardinia and found evidence of continuity and exchange between the colony and broader Sardinia. Despite the initial insights these studies reveal, none of them analyze genome-wide autosomal data, which has proven to be of great use for studies of population history (Pickrell and Reich, 2014).

To provide a more detailed perspective on Sardinian population history, we generated genome-wide data from the skeletal remains of 43 Sardinian individuals radiocarbon dated to between 4,100-1,000 BCE. We analyzed their genetic variation in the context of reference panels of ancient and contemporary individuals. Our goal was to investigate three aspects of Sardinian population history: First, the ancestry of Neolithic Sardinian individuals (ca. 5,700-3,400 BCE) – who were the early peoples expanding onto the island at this time? Second, the genetic structure through the Sardinian Chalcolithic (i.e. Copper Age, ca. 3,400-2,300 BCE) to the Bronze Age (ca. 2,300-1,000 BCE) – were there genetic turnover events through the different cultural transitions observed in the archaeological record? And third, the post-Bronze Age contacts with major Mediterranean civilizations and more recent Italian populations – have they resulted in detectable gene flow? Our results revealed insights about each of these three periods of Sardinian history.

## Results

### Ancient DNA from Sardinia

We organized a collection of skeletal remains from: 1) previously excavated samples throughout Sardinia, in part drawing from samples initially used for isotopic analysis in Lai *et al.* (2013, Supp. Info. 1), and 2) the Seulo caves of central Sardinia (Skeates *et al.*, 2013, Supp. Info. 2). We generated and sequenced DNA libraries enriched for reads overlapping the complete mitochondrial genome as well as a targeted set of 1.2 million single nucleotide polymorphisms (SNPs) (Fu *et al.*, 2015; Haak *et al.*, 2015; Mathieson *et al.*, 2015). After applying several standard ancient DNA quality control filters, we arrived at a final set of 43 individuals with an average coverage of 1.31 × (ranging from 0.04 × to 5.39 × per individual) and a median number of 715,737 sites covered at least once per individual. We obtained age estimates for each individual by either direct radiocarbon dating (*n* = 29), using previously reported radiocarbon dates (*n* = 10), or using a combination of archaeological context and radiocarbon dates from the same burial site (*n* = 4, Fig. 1). The estimated ages in our sample range from 4,100 years BCE to 1,000 years BCE (Fig. 1, Supp. Mat. 1A). To facilitate analyses of temporal structure within ancient Sardinia, we pragmatically grouped the data into three broad periods: Neolithic and Early Copper Age (‘Sar-NECA’, 4,100-3,000 BCE, *n* = 4), Early Middle Bronze Age (‘Sar-EMBA’, 2,500-1,500 BCE, *n* = 24) and Nuragic (‘Sar-Nur’, 1,500-1,000 BCE, *n* = 15). Figure 1 provides an overview of the sample.

**Figure 1:**
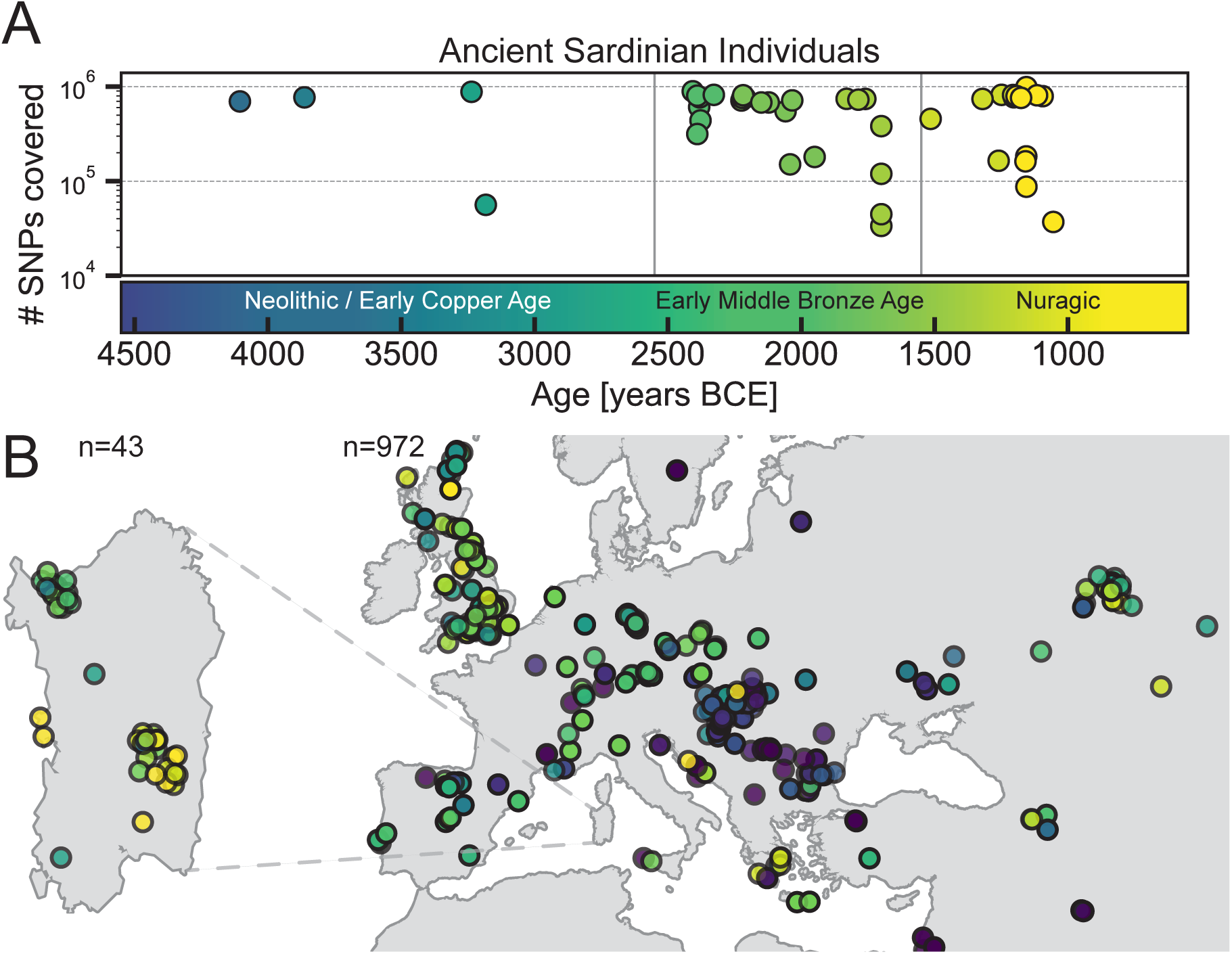
Average depth, sampling locations and ages of ancient individuals. A: The number of SNPs covered at least once and age (mean of 2*σ* radio-carbon age estimates) for the 43 ancient Sardinian individuals. B: The sampling locations of ancient Sardinian individuals and a reference dataset of 972 ancient individuals collected across western Eurasia, spanning a broad temporal period.

### Uniparentally inherited markers

We were able to infer mitochondrial haplogroups for all 43 ancient Sardinian individuals (Supp. Mat. 1E), including a subset (*n*= 10) previously reported by Olivieri *et al.* (2017). We confirm the observation that ancient Sardinian mtDNA haplotypes belong almost exclusively to macro-haplogroups HV (*n* = 16), JT (*n* = 17) and U (*n* = 9), a composition broadly similar to other European Neolithic populations.

Our genome-wide data allowed us to assign Y haplogroups for 25 ancient Sardinian individuals. More than half of them consist of R1b-V88 (*n* = 10) or I2-M223 (*n* = 7) (Sup. Fig. 3, Supp. Mat. 1B). In our reference data set, these two Y-haplogroups appear first in Balkan hunter-gatherer and Balkan Neolithic individuals, and also in more recent western Neolithic populations. Francalacci *et al.* (2013) identified three major Sardinia-specific founder clades based on present-day variation within the haplogroups I2-M26, G2-L91 and R1b-V88, and here we found each of those broader haplogroups in at least one ancient Sardinian individual. Two major present-day Sardinian haplogroups, R1b-M269 and E-M215, are absent (Sup. Fig. 3). Compared to other Neolithic and present-day European populations, the number of identified R1b-V88 carriers is relatively high (Supp. Info 4, Supp. Fig. 4). However, we observed clustering of haplogroups by sample location, consistent with substructure (Supp. Mat. 1B); therefore some caution should be exercised with interpreting our results as estimates for island-wide Y haplogroup frequencies (see Supp. Mat. 1C).

### Results from genome-wide aDNA

We then assessed the relationship of the ancient Sardinian individuals to other ancient and present-day west Eurasian populations using autosomal DNA data. For this purpose, we used: 1) a subset of the Human Origins array dataset from contemporary human individuals (*n* = 1, 963, Lazaridis *et al.*, 2014), 2) an extensive panel of genomic capture data from previously published ancient individuals (*n* = 972, Mathieson *et al.*, 2015; Lazaridis *et al.*, 2016, 2017; Mathieson *et al.*, 2018, 2017; Lipson *et al.*, 2017; Olalde *et al.*, 2018), and 3) a large sample of contemporary Sardinian individuals from our previous studies (*n* = 1, 577, Sidore *et al.*, 2015; Chiang *et al.*, 2018). For some analyses, we grouped these individuals into those from the more isolated Sardinian province of Ogliastra (‘Sar-Ogl’, *n* = 419) and the remainder (‘Sar-non Ogl’, *n* = 1,158). For other analyses, we subset Sardinia into 8 geographic regions (see inset in panel C of Figure 2 for listing and abbreviations, also see Supp. Mat. 1G). Unless otherwise specified (see Materials and Methods), we refer to particular samples of individuals using the group labels used in the datasets they were derived from. As with other human genetic variation studies, consideration of the population annotations is important to consider in the interpretation of results.

**Figure 2:**
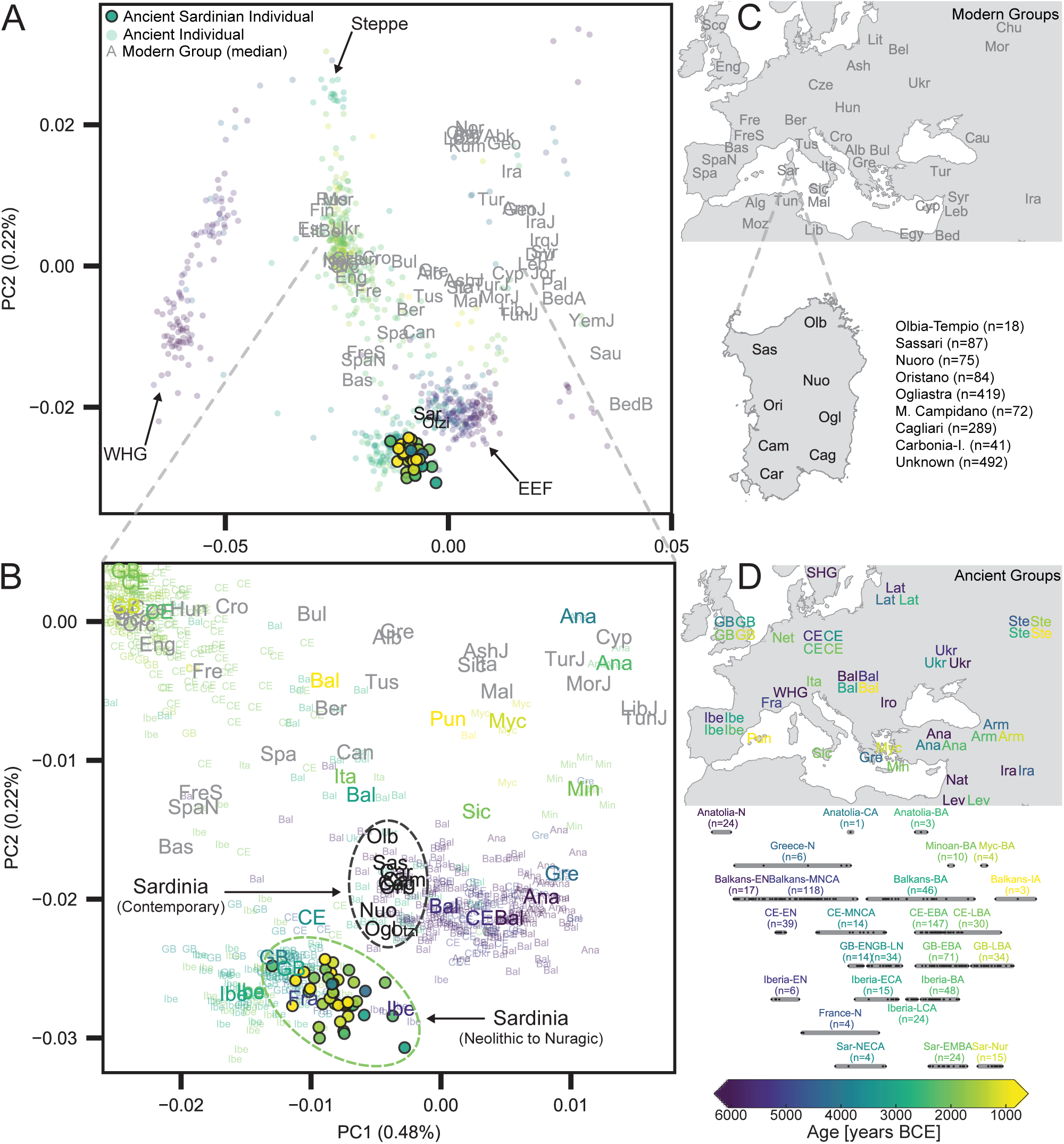
Principal Components Analysis based on the Human Origins dataset. A: Projection of ancient individuals’ genotypes onto principal component axes defined by modern Western Eurasians (gray labels). B: Zoom into the region most relevant for Sardinian individuals. Each projected ancient individual is displayed as a transparent colored point in panel A or a three-letter abbreviation in panel B, with the color determined by the age of each sample (see panel D for legend). In panel B, median PC1 and PC2 values for each population are represented by larger three-letter abbreviations, with black or gray font for moderns and color-coded font based on age for ancient populations. Ancient Sardinian individuals are plotted as circles with edges, and color-coded by age. The full set of labels and abbreviations are described in Sup. Mat. 1F and 1G. C: Geographic legend of present-day individuals from the Human Origins and our Sardinian reference dataset. D: Timeline of selected ancient groups. Note: The same geographic abbreviation can appear multiple times with different colors to represent groups with different median ages.

### Similarity to western mainland Neolithic populations

Importantly, we found low levels of differentiation between Neolithic Sardinian individuals and several Neolithic western mainland European populations, in particular, Cardial Ware-associated groups from Spain (Iberia-EN) and southern France (France-N). When projecting ancient individuals onto the top two principal components (PCs) defined by modern variation, the Neolithic (and also later) ancient Sardinian individuals sit between early Neolithic Iberian and later Copper Age Iberian populations, roughly on an axis that differentiates WHG and EEF populations and embedded in a cluster that additionally includes Neolithic British individuals (Fig. 2). This result is also evident in terms of absolute genetic differentiation, with low pairwise *F*_*ST*_ ≈ 0.005 ± 0.002, Fig. 3) between Neolithic Sardinian individuals and Neolithic western mainland European populations. Pairwise outgroup-*f*_3_ analysis shows a very similar pattern, with the highest values of *f*_3_ (i.e. most shared drift) being with Neolithic and Copper Age Iberia (Fig. 3), gradually dropping off for temporally and geographically distant populations.

**Figure 3:**
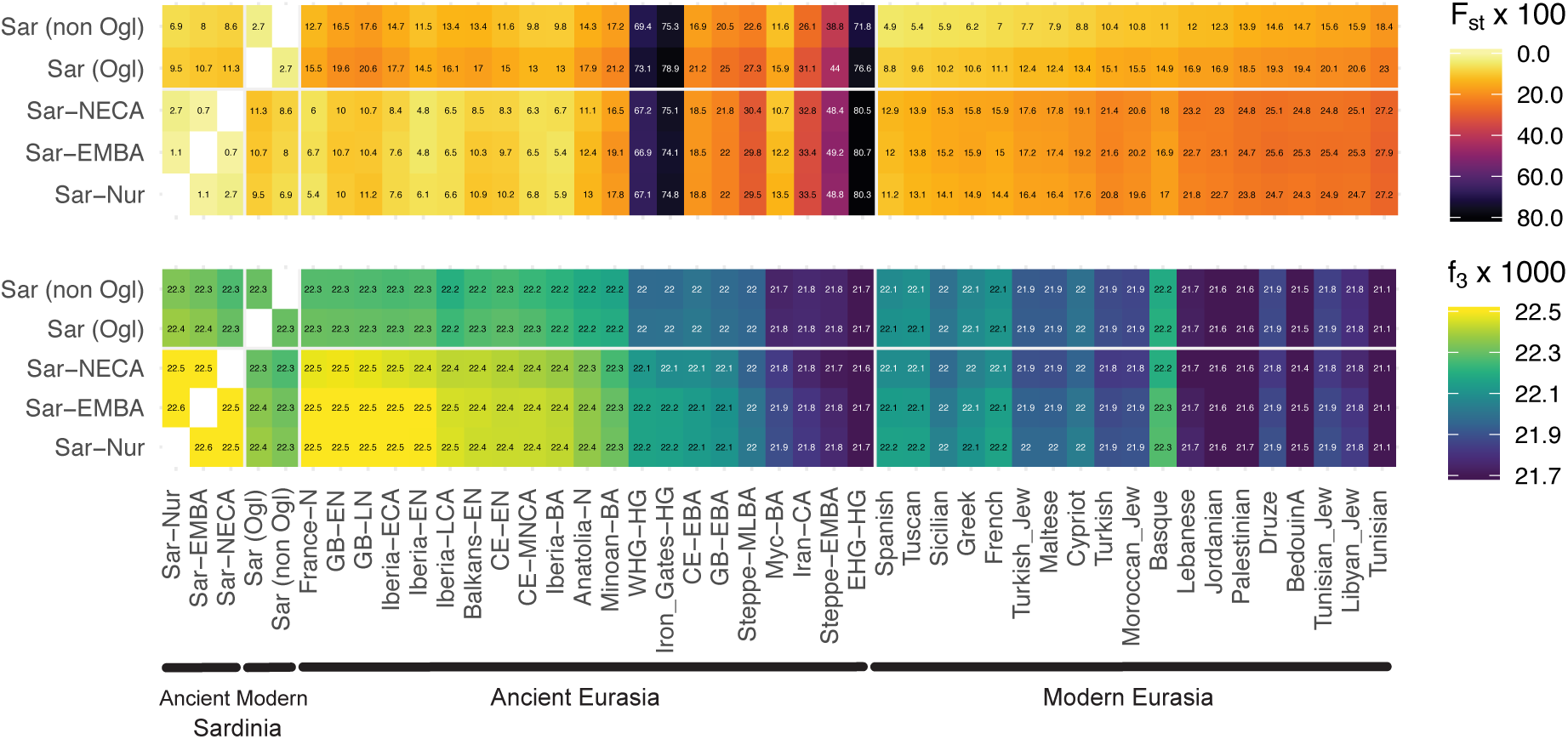
Genetic similarity matrices. We calculated *F*_*ST*_ (upper panel) and outgroup-*f*_3_ (lower panel) of ancient Sardinian and modern Sardinian individuals (grouped into within and outside the Ogliastra region) with each other (left), various ancient (middle), and modern populations (right) of interest.

In explicit admixture models (using qpAdm, see Methods) the southern French Neolithic individuals (France-N) are the most consistent with being a single source for Neolithic Sardinia (*p* ≈ 0.074 to reject the model of one population being the direct source of the other); followed by other populations associated with the western Mediterranean Neolithic Cardial Ware expansion (Supp. Tab. 3). As we discuss below, caution is necessary for interpreting the result as there is a lack of aDNA from other relevant populations of the same period (such as neighboring islands and mainland Italian Cardial Ware cultures).

### Constancy of Western Hunter Gatherer ancestry

Similar to western European Neolithic and central European Late Neolithic populations, ancient Sardinian individuals are shifted towards WHG individuals in the top two PCs relative to early Neolithic Anatolians (Fig. 2). Admixture analysis using qpAdm infers that ancient Sardinian individuals harbour HG ancestry (≈ 17%) that is higher than early Neolithic mainland populations (including Iberia, ≈ 8%), but lower than Copper Age Iberians (≈ 25%) and about the same as Southern French Middle-Neolithic individuals (≈ 21%) (Tab. 1, Supp. Fig. 9). A null model of a two-way admixture between WHG and Neolithic Anatolian populations is inferred to be consistent with our data (*p* ≈ 0.22, Tab. 1). This *p*-value, describing the power to reject the null model of two-way admixture, is similar to the value observed for other western European populations of the early Neolithic (Supp. Tab. 2).

**Table 1:** Results from fitting models of admixture with qpAdm. A) Two-way models of admixture for ancient Sardinia using Western Hunter-Gatherer (WHG) and Neolithic Anatolia (Anatolia-N) individuals as proxy sources. B) Three-way models of admixture for ancient Sardinia using Western Hunter-Gatherer (WHG), Neolithic Anatolia (Anatolia-N), and Early Middle Bronze Age Steppe (Steppe-EMBA, abbreviated Steppe in table), individuals as proxy sources. C) Single-source models to assess continuity of each Sardinian period with the previous one (see main text for guide to abbreviations). E) Results of three-way models showing multiple eastern Mediterranean populations that can produce viable models (Results shown with individuals from Lombardy [Bergamo in the HOA dataset] as one of several possible proxies for north Mediterranean ancestry, see part F). F) Results of three-way models showing multiple north Mediterranean populations that can produce viable models (Results shown with Jewish individuals from Turkey [‘Turkish-Jew’ in the HOA dataset] used as one of several possible proxies for east Mediterranean ancestry, see part E). As a visual aid, p-values greater than 0.01 are bolded. Full results are reported in Supp. Info. 6.

|  | Target | Proxy Source Populations |  |  | p-value | Admixture Fractions |  |  | Standard Error |  |  |
| --- | --- | --- | --- | --- | --- | --- | --- | --- | --- | --- | --- |
|  |  | a | b | c |  | a | b | c | a | b | c |
| A | Sar-NECA | WHG | Anatolia-N | - | **0.677** | 0.173 | 0.827 | - | 0.014 | 0.014 | - |
|  | Sar-EMBA | WHG | Anatolia-N | - | **0.062** | 0.163 | 0.837 | - | 0.007 | 0.007 | - |
|  | Sar-Nur | WHG | Anatolia-N | - | **0.182** | 0.166 | 0.834 | - | 0.009 | 0.009 | - |
| B | Sar-NECA | WHG | Anatolia-N | Steppe | **0.588** | 0.172 | 0.824 | 0.003 | 0.016 | 0.023 | 0.026 |
|  | Sar-EMBA | WHG | Anatolia-N | Steppe | **0.041** | 0.163 | 0.837 | 0.000 | 0.009 | 0.012 | 0.014 |
|  | Sar-Nur | WHG | Anatolia-N | Steppe | **0.123** | 0.167 | 0.833 | 0.000 | 0.010 | 0.014 | 0.016 |
|  | Iberia-EN | WHG | Anatolia-N | Steppe | **0.276** | 0.081 | 0.919 | 0.000 | 0.013 | 0.017 | 0.019 |
| | Iberia-BA | WHG | Anatolia-N | Steppe | $6.0 \cdot 10^{-3}$ | 0.239 | 0.689 | 0.072 | 0.010 | 0.014 | 0.016 |
|  | CE-EN | WHG | Anatolia-N | Steppe | **0.663** | 0.046 | 0.954 | 0.000 | 0.007 | 0.010 | 0.012 |
|  | CE-LBA | WHG | Anatolia-N | Steppe | **0.079** | 0.128 | 0.403 | 0.469 | 0.008 | 0.011 | 0.013 |
| C | Sar-EMBA | Sar-NECA | - | - | **0.548** | - | - | - | - | - | - |
|  | Sar-Nur | Sar-EMBA | - | - | **0.385** | - | - | - | - | - | - |
| | Cagliari | Sar-Nur | - | - | $3.2 \cdot 10^{-37}$ | - | - | - | - | - | - |
| D | Cagliari | Sar-Nur | Sicilian | - | **0.031** | 0.565 | 0.435 | - | 0.021 | 0.021 | - |
|  | Cagliari | Sar-Nur | Maltese | - | **0.013** | 0.590 | 0.410 | - | 0.022 | 0.022 | - |
| | Cagliari | Sar-Nur | Tuscan | - | $2.8 \cdot 10^{-6}$ | 0.540 | 0.460 | - | 0.026 | 0.026 | - |
| | Cagliari | Sar-Nur | Lebanese | - | $1.2 \cdot 10^{-6}$ | 0.724 | 0.276 | - | 0.016 | 0.016 | - |
| | Cagliari | Sar-Nur | Spanish | - | $8.0 \cdot 10^{-26}$ | 0.680 | 0.320 | - | 0.027 | 0.027 | - |
| E | Cagliari | Sar-Nur | N Mediterranean | Turkish-Jew | **0.186** | 0.529 | 0.212 | 0.259 | 0.023 | 0.043 | 0.035 |
|  | Cagliari | Sar-Nur | N Mediterranean | Libyan-Jew | **0.086** | 0.532 | 0.272 | 0.196 | 0.024 | 0.039 | 0.028 |
|  | Cagliari | Sar-Nur | N Mediterranean | Maltese | **0.07** | 0.562 | 0.086 | 0.351 | 0.027 | 0.074 | 0.060 |
|  | Cagliari | Sar-Nur | N Mediterranean | Tunisian-Jew | **0.064** | 0.515 | 0.285 | 0.200 | 0.023 | 0.037 | 0.029 |
|  | Cagliari | Sar-Nur | N Mediterranean | Sicilian | **0.06** | 0.546 | 0.065 | 0.389 | 0.024 | 0.068 | 0.060 |
|  | Cagliari | Sar-Nur | N Mediterranean | Moroccan-Jew | **0.05** | 0.536 | 0.247 | 0.217 | 0.024 | 0.044 | 0.033 |
|  | Cagliari | Sar-Nur | N Mediterranean | Lebanese | **0.049** | 0.571 | 0.269 | 0.160 | 0.026 | 0.040 | 0.023 |
|  | Cagliari | Sar-Nur | N Mediterranean | Druze | **0.04** | 0.562 | 0.260 | 0.178 | 0.024 | 0.040 | 0.025 |
|  | Cagliari | Sar-Nur | N Mediterranean | Cypriot | **0.024** | 0.538 | 0.270 | 0.192 | 0.023 | 0.037 | 0.027 |
|  | Cagliari | Sar-Nur | N Mediterranean | Jordanian | **0.018** | 0.563 | 0.291 | 0.146 | 0.025 | 0.037 | 0.022 |
|  | Cagliari | Sar-Nur | N Mediterranean | Palestinian | **0.014** | 0.563 | 0.301 | 0.135 | 0.026 | 0.037 | 0.020 |
| | Cagliari | Sar-Nur | N Mediterranean | Turkish | $8.6 \cdot 10^{-3}$ | 0.668 | 0.089 | 0.243 | 0.037 | 0.074 | 0.042 |
| | Cagliari | Sar-Nur | N Mediterranean | Tunisian | $1.0 \cdot 10^{-4}$ | 0.547 | 0.375 | 0.079 | 0.028 | 0.034 | 0.014 |
| F | Cagliari | Sar-Nur | E Mediterranean | Lombardy | **0.186** | 0.529 | 0.259 | 0.212 | 0.023 | 0.035 | 0.043 |
|  | Cagliari | Sar-Nur | E Mediterranean | Greek | **0.153** | 0.560 | 0.165 | 0.274 | 0.020 | 0.054 | 0.057 |
|  | Cagliari | Sar-Nur | E Mediterranean | Tuscan | **0.115** | 0.544 | 0.215 | 0.241 | 0.022 | 0.048 | 0.056 |
|  | Cagliari | Sar-Nur | E Mediterranean | French | **0.094** | 0.573 | 0.321 | 0.105 | 0.021 | 0.028 | 0.024 |
|  | Cagliari | Sar-Nur | E Mediterranean | Basque | **0.066** | 0.548 | 0.337 | 0.115 | 0.023 | 0.026 | 0.026 |
|  | Cagliari | Sar-Nur | E Mediterranean | Spanish | **0.052** | 0.555 | 0.309 | 0.137 | 0.022 | 0.030 | 0.032 |

### Continuity from the Sardinian Neolithic through the Nuragic

We found several lines of evidence supporting genetic continuity from the Sardinian Neolithic into the Bronze Age and Nuragic times. Importantly, we observed low genetic differentiation between ancient Sardinian individuals from various time periods. We estimated *F*_*ST*_ to be 0.0027 ± 0.0014 between Neolithic and late Bronze Age (mostly Nuragic) Sardinian individuals (Fig. 3). Furthermore, we did not observe temporal substructure within the ancient Sardinian individuals in the top two PCs – they form a coherent cluster (Fig. 2). In stark contrast, ancient individuals from many mainland geographic regions, such as central Europe, show larger movements over the first two PCs from the Late Neolithic to the Bronze Age, and also have higher pairwise differentiation (*F*_*ST*_=0.0194 ±0.0003).

In the presence of significant influx, differential genetic affinity of a test population *x* would cause *f*_4_ statistics of the form *f* (Sard Period 1 - Sard Period 2; Pop *x* - Ancestral Allele) to be non-zero (where “Ancestral Allele” is an inferred ancestral allelic state from a multi-species alignment). However, we observe that no such statistic differs significantly from zero for all test populations *x* (Supp. Mat. 2D). A qpAdm analysis, which is based on simultaneously testing *f*-statistics with a number of outgroups and adjusts for correlations, cannot reject a model of Neolithic Sardinian individuals being a direct predecessor of Nuragic Sardinian individuals either (*p* = 0.54, Supp. Tab. 3). Our qpAdm analysis further shows that the WHG ancestry proportion, in a model of admixture with Neolithic Anatolia, remains stable at ∼17% throughout three ancient time-periods (Tab. 1A). When using a three-way admixture model, we do not detect significant Steppe ancestry in any ancient Sardinian individual, as is inferred, for example, in later Bronze Age Iberians (Tab. 1B, Supp. Fig. 9).

### From the Nuragic to present-day Sardinia

Our results demonstrate that ancient Sardinian individuals are genetically closest to contemporary Sardinian individuals among all the ancient individuals analyzed (Fig. 3), and relative to other European populations, there is lower differentiation between present-day and ancient individuals (Supp. Fig. 7). However, we also find multiple lines of evidence for appreciable gene flow into Sardinia after the Nuragic period.

Firstly, present-day Sardinian individuals are shifted from the ancients towards more eastern Mediterranean populations on the western Eurasian PCA (Fig. 2). We observe a corresponding signal in our *f*_4_ analysis, in that we see significantly higher affinity of many present-day and some ancient populations to modern Sardinian versus Nuragic Sardinian individuals (*f*_4_ of the form *f* (Mod Sard - Ancient Sard; Pop *x* - Ancestral Allele), see Fig. 4 and Supp. Mat. 2D). Similarly, *f*_3_ statistics that directly test for admixture of present-day Sardinians, with Nuragic Sardinian individuals as one source, yield highly significant negative values, indicating admixture (Fig. 4). Using qpAdm we find that models of continuity from Nuragic Sardinia to present-day Sardinian populations (e.g. Cagliari) without influx are rejected (*p* < 10^−40^, Tab. 1C). Moreover, genetic differentiation between the Nuragic and present is higher than across ancient periods (between Nuragic and present-day non-Ogliastra individuals pairwise *F*_*ST*_ = 0.00695 ± 0.00041; compared to *F*_*ST*_ = 0.0027 ± 0.0014 between Late Neolithic and Nuragic individuals.)

**Figure 4:**
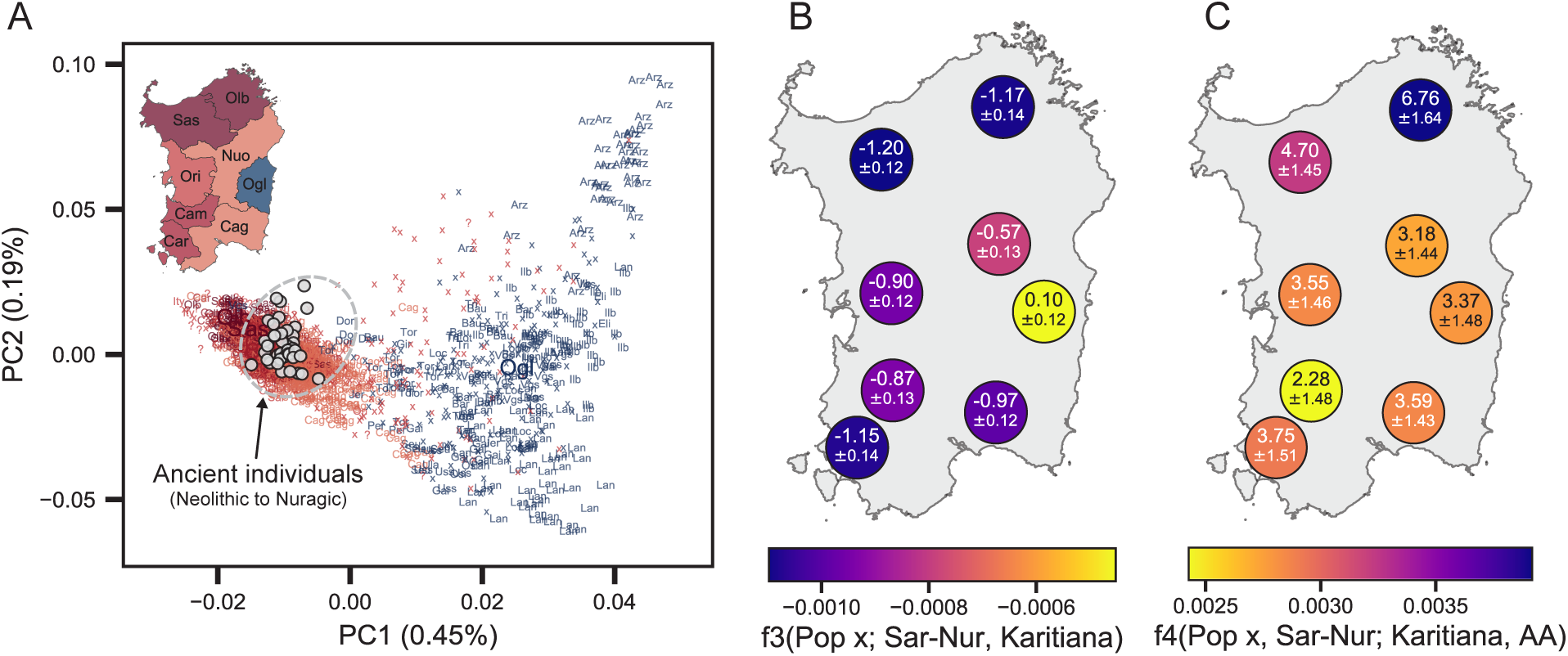
Present-day genetic structure in Sardinia reanalyzed with aDNA. A: Scatter plot of the first two principal components trained on 1577 present-day individuals with grand-parental ancestry from Sardinia. Each individual is labeled with a location if at least 3 of the 4 grandparents were born in the same geographical location (“small” three letter abbreviations); otherwise with “x” or if grand-parental ancestry is missing with “?”. We calculated median PC values for each Sardinian province (large abbreviations). We also projected each ancient Sardinian individual on to the top two PCs (gray points). B/C: We plot *f*-statistics that test for admixture of modern Sardinian individuals (grouped into provinces) when using Nuragic Sardinian individuals as one source population. Uncertainty ranges depict one standard error (calculated from block bootstrap). Karitiana are used in the *f*-statistic calculation as a proxy for ANE/Steppe ancestry (Patterson *et al.*, 2012).

Second, we find many populations that can produce significant *f*_4_ and *f*_3_ statistics consistent with admixture (Supp. Mat. 2C and D). Many of these populations carry high levels of Ancestral North Eurasian (ANE) ancestry, and likely serve as a proxy for ancient Eurasian ancestry that entered Europe after the Neolithic with Steppe expansions, as similarly observed for many present-day mainland Europeans (Patterson *et al.*, 2012).

ADMIXTURE analysis gives further insight into this signal of gene flow. While contemporary Sardinian individuals show the highest affinity towards EEF-associated populations among all of the modern populations, they also display membership with other clusters (Fig. 5). In contrast to ancient Sardinian individuals, present-day Sardinian individuals carry a modest “Steppe-like” ancestry component (but generally less than continental present-day European populations), and an appreciable broadly “eastern Mediterranean” ancestry component (also inferred at a high fraction in other present-day Mediterranean populations, such as Sicily and Greece).

**Figure 5:**
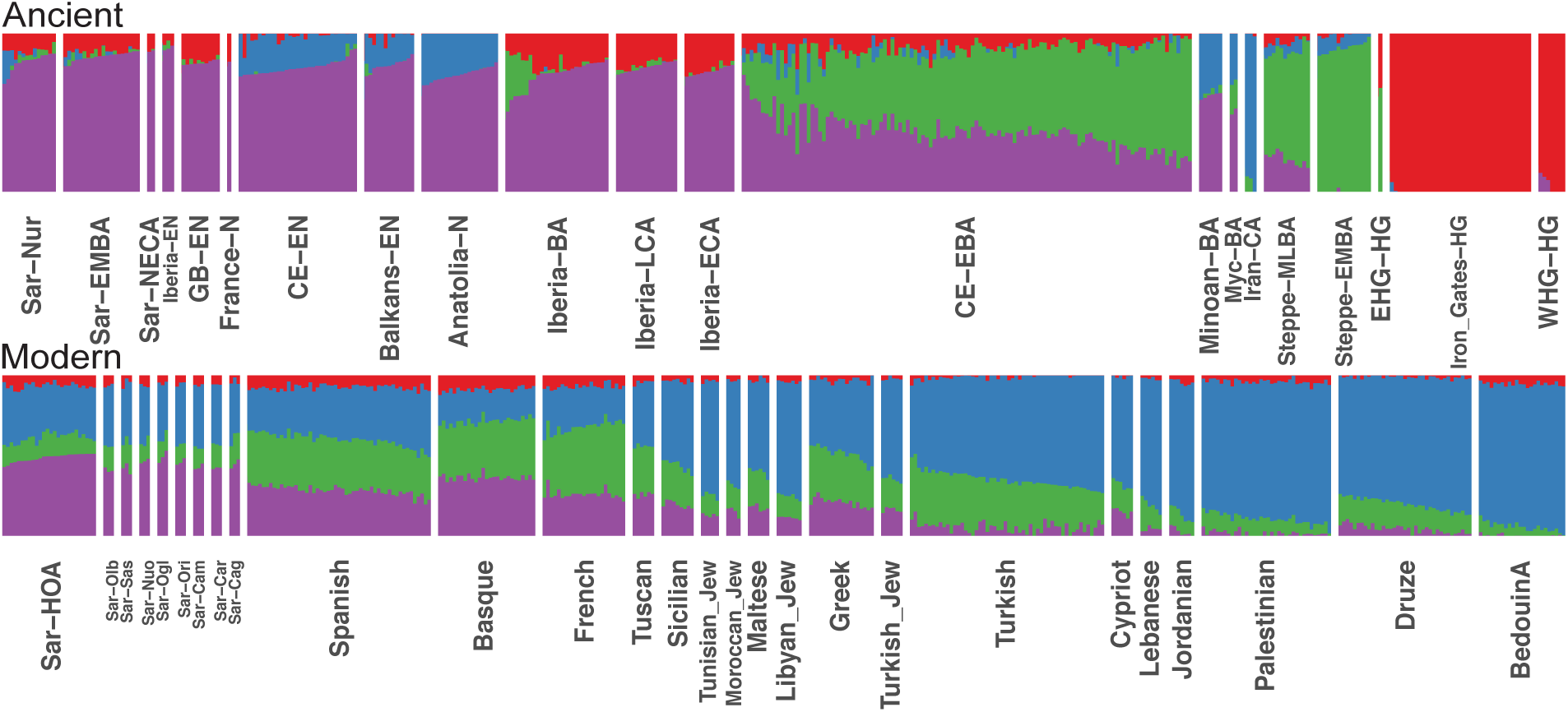
Admixture coefficients estimated by ADMIXTURE (*K* = 4). Each stacked bar represents one individual and color fractions depict the fraction of the given individual’s ancestry coming from a given “cluster”. For *K* = 4 (depicted here), ancient Sardinian individuals share similar admixture proportions as other western European Neolithic individuals. Present-day Sardinian individuals additionally have elevated Steppe-like ancestry (but less than other European populations), and an additional ancestry component prevalent in Near Eastern / Levant populations. ADMIXTURE results for all *K*=2, …, 11 are depicted in the supplement (Supp. Fig. 14).

To further characterize signatures of admixture, we used qpAdm to test the fit of a model of present-day Sardinian populations as a simple two-way admixture between Nuragic Sardinian individuals and potential other source populations (Table 1D, Supp. Tab. 4). A model of admixture with modern Sicilians (*p* = 0.031) had the best support, followed by Maltese (*p* = 0.0128), Turkish (*p* = 0.0086) and Greeks (*p* = 0.00071). For the model of a mixture of Sicilians and ancient Sardinian individuals, we infer an admixture proportion of 43.5 ± 2.1 percent Sicilian admixture (Tab. 1, Sup. Tab. 4, Supp. Fig. 10).

We also considered three-way models of admixture with qpAdm to further refine the geographic origins of this recent admixture signal (Supp. Info 6). Indeed, we find models with admixture between Nuragic Sardinia, one northern Mediterranean source and one eastern Mediterranean source fit well (*p* > 0.01 for several combinations, Table 1E,F). For a representative sample from Sardinia (Cagliari), across various proxies (excluding Sicily and Malta) the admixture fractions range 10-30% for the “northern Mediterranean” component, 13-33% for the “eastern Mediterranean”, with the remaining 52-57% coming from Nuragic Sardinia. For models with Sicilian or Maltese as proxy sources, the estimates of ancestry for the N. Mediterranean component shrink to small values (8.6% and 6.5% for Sicilian and Maltese, respectively), essentially bringing the fitted parameters towards the two-way mixture models. Maltese and Sicilian individuals appear to reflect a mixture of N. Mediterranean and E. Mediterranean ancestries, and as such they can serve as single-source proxies in two-way admixture models with Nuragic Sardinia (Table 1D).

Caution is warranted when interpreting inferred admixture fractions with each of these simple models; however, the signal across multiple analyses indicates that complex post-Nuragic gene flow, partly from sources originating in the eastern Mediterranean and partly from the northern Mediterranean, has likely played a role in the population genetic history of Sardinia.

### Fine-scale structure in contemporary Sardinia

Ancient DNA can shed new light on present-day genetic variation. We, therefore, re-assessed spatial substructure previously observed in a dense geographic modern sampling (1,577 whole genome sequences) from Sardinia (Chiang *et al.*, 2018).

In a PCA of the modern Sardinian variation, individuals from Ogliastra fall furthest away from the ancient Sardinian individuals (Chiang *et al.*, 2018) (Fig. 4). In stark contrast, in the PCA of modern Western Eurasian variation, the pattern reverses: Ogliastra is placed closest of all provinces to the ancient Sardinian individuals (Fig. 2). Direct tests for admixture using *f*_3_ statistics with Nuragic Sardinian individuals as one source yielded highly significant results for all present-day provinces except Ogliastra (Fig. 4). The non-significant value of Ogliastra can have two causes: An actual lack of admixture or high levels of drift that mask admixture *f*_3_. However, the *f*_4_ statistics and admixture proportions of qpAdm are robust to recent drift of the admixed population, and in both analyses Ogliastra shows an admixture signal that is only slightly weaker than most other provinces (Fig. 4, Supp. Fig. 12). Together, these results suggest high levels of drift specific to Ogliastra (likely also driving the first two PCs of present-day Sardinian variation), but simultaneously also less admixture than other Sardinian provinces.

In the previous section, we reported finding that many non-Sardinian modern populations have a higher affinity to present-day Sardinian individuals than to Nuragic Sardinian individuals (using a *f*_4_ statistic of the form *f*_4_(Mod. Sard Pop *y* - Sar-Nur; Pop *x* - Ancestral Allele) where *x* are test non-Sardinian modern populations, Fig. 4, Sup. Mat. 2D). Interestingly, the northern province Olbia (north-east) and to some degree also Sassari (north-west) have the highest affinity to most tested populations (Fig. 4). A three-way admixture model fit with qpAdm finds a similar signal. In a model with Tuscan as a proxy for northern Mediterranean immigration and Lebanese as a proxy for a second additional, more eastern Mediterranean source, the inferred admixture fractions vary across Sardinia, with the highest eastern Mediterranean ancestry in the southwest (Carbonia, Campidano) and the highest northern Mediterranean ancestry in the northeast of the island (Olbia, Sassari, Supp. Fig. 12). In addition, we observed a marked shift of individuals from Olbia and Sassari towards continental populations in the PCA (Fig. 4).

## Discussion

Our analysis of genome-wide data from 43 ancient Sardinian individuals generated new direct evidence regarding the population history of Sardinia and the Mediterranean. Importantly, we detected a strong genetic affinity of Neolithic Sardinian individuals to other early Neolithic western Mediterranean populations. This signal is especially interesting in light of archaeological evidence for a rapid maritime spread of Neolithic Cardial Impressed Ware culture through the western Mediterranean occurring around 5,500 BCE (Zilhão, 2001; Martins *et al.*, 2015). While we lack aDNA data from Neolithic mainland Italy, a putative center of this spread to Sardinia, we find that Neolithic Sardinian individuals are closely related to other hypothesized populations of this initial wave, in particular, Neolithic Spanish and southern French populations. This new evidence points towards Neolithic Sardinian individuals descending principally from mainland Neolithic populations. This hypothesis is also consistent with a signal of population turnover associated with the Neolithic transition observed in Sardinian ancient mtDNA (Modi *et al.*, 2017), and with a gap in the Sardinian archaeological record before its Neolithic transition (Lugliè, 2018).

Neolithic Sardinian individuals fit well as a two-way admixture between EEF and WHG sources, similar to other EEF populations including Linear Pottery and Cardial cultures. Recent evidence based on aDNA indicates that traces of WHG ancestry were already part of the initial wave of the Mediterranean Neolithic transition and that similar to other EEF populations, subsequent local admixture increased WHG ancestry substantially over time in Iberia (Lipson *et al.*, 2017). In stark contrast, in Sardinia, we observed remarkable constancy of WHG ancestry close to 20% throughout our sampling periods, well into the second millennium BCE. This reflects the continuity of Sardinia through this time period and is consistent with a model of a low density or even absence of local Mesolithic hunter-gatherers at the time of arrival of EEF individuals (Lugliè, 2018). However, we can not rule out a model with an initial pulse of local admixture. Genome-wide data from a Mesolithic or a very early Neolithic individual from Sardinia could help settle this question.

Additional insight into the origins of Neolithic populations of Sardinia comes from Y chromosome variation in the ancient samples. We detected Y haplogroups R1b-V88 and I2-M223 in the majority of the ancient Sardinian males. In our reference dataset, both haplogroups appear earliest in Mesolithic hunter-gatherers and then Neolithic groups of the Balkans (Mathieson *et al.*, 2018) and also EEF Iberians, but not in Neolithic Anatolians or more western WHG individuals. Further sampling is necessary, but the current data are consistent with hypotheses of the expansion of Cardial Impressed Ware related cultures through the Mediterranean via the Balkans. Future studies, including ancient DNA from early Neolithic sites in mainland Italy, will help to further resolve details of these putative migrations.

From the Neolithic onwards, Sardinia appears to have been relatively isolated until at least the late second millennium BC, unlike many other parts of Europe which had experienced substantial gene flow from central Eurasian Steppe ancestry starting about 3,000 years BCE (Haak *et al.*, 2015; Allentoft *et al.*, 2015). While we cannot exclude influx from genetically similar populations such as early Iberian Bell Beakers, the absence of Steppe ancestry suggests genetic isolation from many Bronze Age mainland populations - including later Iberian Bell Beakers, who would already have carried substantial Steppe ancestry (Olalde *et al.*, 2018). As further support, the Y haplogroup R1b-M269, the most frequent present-day western European haplogroup and the haplogroup associated with expansions that brought Steppe ancestry into Britain (Olalde *et al.*, 2018) and Iberia (Olalde *et al.*, 2019) about 2,500-2,000 BCE, remains absent in our sample of ancient Sardinian individuals through the end of our sampling period (1,200-1,000 BCE).

The genetic continuity inferred throughout our ancient sampling period does not continue fully into the present. Previously, admixture tests based on *f*-statistics did not provide significant evidence for gene flow (Chiang *et al.*, 2018), likely because no suitable proxy for the Nuragic Sardinian ancestry component was available. Here, including direct aDNA data increased the power of admixture tests, which resulted in uncovering multiple lines of evidence of moderate gene flow into Sardinia. This post-Nuragic admixture likely brought additional Y chromosome haplotype diversity to Sardinia (Sup. Fig. 3), such as R1b-M269 and also E-M215 (now prevalent in northern Africa).

We find evidence for at least two phases of post-Nuragic gene flow. First, there is a general shift towards central and eastern Mediterranean sources, demonstrated by the direction of the overall change in the PCA and ADMIXTURE, and the results of modeling population relationships using qpAdm. Second, we detected variation in the signals in the PCA and qpAdm analysis suggesting that the northern provinces of Olbia, and to a lesser degree Sassari, have received more northern Mediterranean immigration after the Bronze Age than the other provinces; meanwhile the southwestern provinces of Campidano and Carbonia show more eastern Mediterranean ancestry. Together, these signals suggest temporally and geographically complex post-Nuragic gene flow into Sardinia. Ultimately, aDNA data from these historical periods will be needed to clarify and refine the interpretation.

A preliminary hypothesis would be that an influx from eastern Mediterranean sources is overlayed by more recent influx from the Italian mainland. Historically, both of these seem plausible. Sardinia hosted major Phoenician colonies in the first millennium BCE, principally along the south and west coasts of the island, and previous studies based on uni-parentally inherited markers have found evidence for Phoenician contact and gene flow (Zalloua *et al.*, 2008; Matisoo-Smith *et al.*, 2018). Sardinia was also an important Roman province and then was later under occupation by the Vandals and the Byzantine Empire. There are also more recent sources of immigration in the last few hundred years from Italy, Spain, and Corsica. Shepherds from Corsica immigrated to occupy large pastures left largely empty since the late Middle Ages, bringing an Italian-Corsican dialect (Gallurese) now prevalent in the northeastern part of Sardinia (Lannou, 1941). The differing historical impacts of these external contacts in different regions of Sardinia is supported in the patterns we observe, with more northern Mediterranean ancestry inferred in the north (where Gallurese is prevalent), eastern Mediterranean ancestry inferred in the south and west of Sardinia (where more Punic colonies existed), and more isolation in central regions of Ogliastra and Nuoro.

The evidence for gene flow after the second millennium BCE seems to contradict previous models emphasizing Sardinian isolation, but we confirm that contemporary Sardinian individuals have retained an exceptionally high degree of EEF ancestry (Haak *et al.*, 2015). Compared to other European populations, Sardinia experienced relative genetic isolation through the Bronze age, and our models also fit the majority of modern Sardinian ancestry being retained from the Nuragic period. The subsequent post-Nuragic admixture appears to derive from Mediterranean sources that have relatively little Steppe ancestry (Sarno *et al.*, 2017; Lazaridis *et al.*, 2017). Therefore, contemporary Sardinians still cluster with several mainland European Copper Age individuals such as Ö tzi (Sikora *et al.*, 2014), even as they are shifted from ancient Sardinian individuals of a similar time period (Fig. 2).

The history of gene flow into Sardinia is also relevant to understanding its relationship to the Basque populations of Iberia. Previous studies have suggested both present-day and ancient Basque individuals share a genetic connection with modern Sardinian individuals (Günther *et al.*, 2015; Chiang *et al.*, 2018). We detected a similar signal, with modern Basque having, of all modern samples, the largest pairwise outgroup-*f*_3_ with Sardinians in each of our time periods (Fig. 3). A plausible explanation arises from the observation that both Basque and Sardinians have remained relatively isolated since the Neolithic transition (e.g. see Olalde *et al.*, 2019, for novel aDNA evidence on the Basque). While both Basque and Sardinians have received some immigration, apparently from different sources, both populations also retained an exceptionally high fraction of EEF ancestry (e.g., Fig. 5). This shared ancestry component likely contributes to the high pairwise outgroup-*f*_3_ (Fig. 3) between Basque and Sardinians, and explains how both populations share a genetic affinity despite their geographic separation.

Overall, we find that genome-wide ancient DNA provides unique insights into the population history of Sardinia. We do not detect any significant admixture from the the Neolithic period of Sardinia through the Nuragic. From the Nuragic to the present we observe a significant shift in ancestry tied to northern and eastern Mediterranean sources. Genetic analyses that include Sardinian individuals spanning the post-Nuragic period to the present, as well as individuals from plausible sources of this gene flow, such as Punic, Roman, and other Mediterranean groups, will help to more precisely date these events and to relate them to the historical and archaeological record. Ultimately, having a more refined model of post-Nuragic demographic history will provide a better framework to understand the evolutionary history of genetic disease-variants prevalent in Sardinia and throughout the Mediterranean, such as beta-thalassemia and G6PD deficiency.

## Materials and Methods

### Archaeological sampling

The archaeological samples used in this project derive from two major collection avenues. The first was a sampling effort led by co-author Luca Lai, leveraging a broad base of samples from different existing collections in Sardinia, a subset of which were previously used in isotopic analyses to understand dietary composition and change in prehistoric Sardinia (Lai *et al.*, 2013). The second was from the Seulo Caves project (Skeates *et al.*, 2013), an on-going project on a series of caves that span the Middle Neolithic to late Bronze Age near the town of Seulo. The project focuses on the diverse forms and uses of caves in the prehistoric culture of Sardinia. All samples were handled in collaboration with local scientists and with the approval of the local Sardinian authorities for the handling of archaeological samples (Ministero per i Beni e le Attivita Culturali, Direzione Generale per i beni Archeologici, request dated 11 August 2009; Soprintendenza per le Beni Archeologici per le province di Sassari e Nuoro, prot. 12278 dated 05 Dec. 2014; Soprintendenza ai Beni Archeologici per le Province di Cagliari e Oristano, prot. 62, dated 08 Jan 2015; Soprintendenza Archeologia, Belle arti e Paesaggio per le provincie di Sassari, Olbia-Tempio e Nuoro, prot. 4247 dated 14 March 2017: Soprintendenza per i Beni Archeologici per le Province di Sassari e Nuoro, prot. 12930 dated 30 Dec. 2014: Soprintendenza Archeologia, belle arti e paesaggio per le province di Sassari e Nuoro, prot. 7378 dated 9 May, 2017: Soprintendenza Archeologia, belle arti e paesaggio per le province di Sassari e Nuoro, prot. 16258 dated 26 Nov. 2017). For more, detailed description of the sites please see Supplemental Information Sections 1 and 2.

### Initial sample screening and sequencing

The ancient DNA (aDNA) workflow was implemented in dedicated facilities at the Palaeogenetic Laboratory of the University of Tübingen and at the Department of Archaeogenetics of the Max Planck Institute for the Science of Human History in Jena. The only exception was for four samples from the Seulo Cave Project which had DNA isolated at the Australian Centre for Ancient DNA and capture and sequencing carried out in the Reich lab at Harvard University. Different skeletal elements were sampled using a dentist drill to generate bone and tooth powder respectively. DNA was extracted following an established aDNA protocol (Dabney *et al.*, 2013) and then converted into double-stranded libraries retaining (Meyer and Kircher, 2010) or partially reducing (Rohland *et al.*, 2015) the typical aDNA substitution pattern resulting from deaminated cytosines that accumulate towards the molecule’s termini. After indexing PCR (Meyer and Kircher, 2010) and differential amplification cycles, the DNA was shotgun sequenced on Illumina platforms. Samples showing sufficient aDNA preservation where captured for mtDNA and ≈1.24 million SNPs across the human genome chosen to intersect with the Affymetrix Human Origins array and Illumina 610-Quad array (Fu *et al.*, 2015). The resulting enriched libraries were also sequenced on Illumina machines in single-end or paired-end mode. Sequenced data were pre-processed using the `EAGER` pipeline (Peltzer *et al.*, 2016). Specifically, DNA adapters were trimmed using `AdapterRemoval` v2 (Schubert *et al.*, 2016) and paired-end sequenced libraries were merged. Sequence alignment to the mtDNA (RSRS) and nuclear (hg19) reference genomes was performed with `BWA` (Li and Durbin, 2009) (parameters –n 0.01, seeding disabled), duplicates were removed with `DeDup` (Peltzer *et al.*, 2016) and a mapping quality filter was applied (MQ ⩾ 30). For genetic sexing, we compared relative X and Y-chromosome coverage to the autosomal coverage with a custom script. For males, nuclear contamination levels were estimated based on heterozygosity on the X-chromosome with the software `ANGSD` (Korneliussen *et al.*, 2014). Data originating from mtDNA capture was processed with `schmutzi` (Renaud *et al.*, 2015), which jointly estimates mtDNA contamination and reconstructs mtDNA consensus sequences that were assigned to the corresponding mtDNA haplogroups using `Haplofind` (Vianello *et al.*, 2013) (Supp. Mat. 1D). We applied several standard ancient DNA quality control metrics: We retained endogenous DNA content in shotgun sequencing >0.2%, evidence of an average damage pattern present at the molecule termini, mtDNA contamination <4% (average 1.6%) and nuclear contamination <6% (average 1.1%).

We next generated genotype calls that were used for downstream population genetic analyses. To account for sequencing errors we first removed any reads that overlapped a SNP on the capture array with a base quality score less than 20. We also removed the last 3-bp on both sides of every read to reduce the effect of DNA damage on the resulting genotype calls (Al-Asadi *et al.*, 2018). With these filtered aligned reads in hand, we used custom python scripts (https://github.com/mathii/gdc3) to generate pseudo-haploid genotypes by sampling a random read for each SNP on the capture array and setting the genotype to be homozygous for the allele present on the randomly sampled read.

### Merging newly generated data with published data

#### Ancient DNA datasets from Western Eurasia

To provide context for the study of ancestry of the ancient individuals from Sardinia, we downloaded and processed several ancient datasets from continental Europe and the Middle-east (Mathieson *et al.*, 2015; Lazaridis *et al.*, 2016, 2017; Mathieson *et al.*, 2018, 2017; Lipson *et al.*, 2017; Olalde *et al.*, 2018). To minimize technology-specific batch effects in genotype calls and thus downstream population genetic inference, we focused on previously published ancient samples that had undergone the capture protocol on the same set of SNPs targeted in our study. We processed these samples through the same pipeline and filters described above, resulting in a dataset of 972 ancient samples. Throughout our analysis, we used a subset of *n* = 1,013,439 variants that was created by removing SNPs missing in more than 80% of all ancients individuals (Sardinian and reference dataset) with at least 60% of all captured SNPs covered.

This ancient dataset spans a wide geographic distribution and temporal range. Ancient individuals are associated with a variety of different cultures, which provides rich context for interpreting downstream results. Our reference ancient dataset is comprised of many individuals sampled from a particular geographic locale, such as Germany or Hungary, in a transect of multiple cultural changes through time (Fig. 2). For the PCA (Fig. 2), we additionally included a single low-coverage ancient individual (label “Pun”) dated to 361-178 BCE from a Punic necropolis on the west Mediterranean island of Ibiza (Zalloua *et al.*, 2018).

We merged individuals into groups (Supp. Mat. 1F,G). For ancient samples, these groups were chosen manually, trying to strike a balance between reducing overlap in the PCA and keeping culturally distinct populations separate. We used geographic location to first broadly group samples into geographic areas (such as Iberia, Central Europe and Balkans), and then further annotated each of these groups by different time periods.

#### Contemporary DNA datasets from Western Eurasia

We downloaded and processed the Human Origins dataset to characterize a subset of Eurasian human genetic diversity at 594,924 autosomal SNPs (Lazaridis *et al.*, 2014). To be consistent with previous studies (Lazaridis *et al.*, 2014; Mathieson *et al.*, 2015), we focused on a subset of 777 individuals from Western Eurasia.

#### Contemporary DNA dataset from Sardinia

We merged in a whole-genome sequence dataset which was described and previously analyzed by Chiang *et al.* (2018). It consists of 1,577 unrelated individuals, grouped into multiple geographic regions within Sardinia (Fig. 2C).

### Principal Components Analysis

We performed Principal Components Analysis (PCA) on two large-scale datasets of modern genotypes from Western Eurasian (777 individuals from the Human Origins dataset) and Sardinia (1,577 individuals from the SardiNIA project). For both datasets, we normalized the genotype matrix by mean-centering and scaling the genotypes at each SNP using the inverse of the square-root of heterozygosity (Patterson *et al.*, 2006). We additionally filtered out rare variants with minor allele frequency (*p*_min_ < 0.05).

To assess population structure in the ancient individuals, we projected them onto our pre-computed principal axes using only the non-missing SNPs via a least-squares approach, and correcting for the shrinkage effect observed in high-dimensional PC score prediction (Reich *et al.*, 2008; Lee *et al.*, 2010). More details on how we corrected the biased PC scores are discussed in Supp. Info. 8.

We also projected a number of out-sample sub-populations from Sardinia onto our PCs. Reassuringly, these out-of-sample Sardinian individuals project very close to Humans Origins Sardinian individuals (Fig. 2). Moreover, the test-set Sardinia individuals with grand-parental ancestry from Southern Italy cluster with reference individuals with ancestry from Sicily.

### ADMIXTURE Analysis

We applied `ADMIXTURE` to an un-normalized genotype matrix of ancient and modern samples (Alexander *et al.*, 2009). `ADMIXTURE` is a maximum-likelihood based method for fitting the Pritchard, Stephens and Donnelly model Pritchard *et al.* (2000) using sequential quadratic programming. We first LD pruned the data matrix based off the modern Western Eurasian genotypes, using plink1.9 with parameters [`--indep-pairwise 200 25 0.4`]. We then ran 5 replicates of `ADMIXTURE` for values of *K* 2, …, 11. We display results for the replicate that reached the highest log-likelihood after the algorithm converged (Supp. Fig. 8).

### Estimation of *f*-statistics

We measured similarity between groups of individuals through computing an outgroup-*f*_3_ statistic. The outgroup-*f*_3_ statistic can be interpreted as a measure of the internal branch length of a three-taxa population phylogeny and thus does not depend strongly on genetic drift or systematic error in the focal pair of populations that are being compared (Patterson *et al.*, 2012).

Here we used the ancestral allelic states as an outgroup, inferred from a multi-species alignment from Ensembl Compara release 59, as annotated in the 1000 Genomes Phase3 sites vcf (ftp://ftp.1000genomes.ebi.ac.uk/vol1/ftp/release/20130502/ALL.wgs.phase3_shapeit2_mvncall_integrated_v5b.20130502.sites.vcf.gz) (1000 Genomes Project Consortium *et al.*, 2015). We fixed the ancestral allele counts to *n* = 10^6^ to avoid finite sample size correction when calculating outgroup *f*_3_.

The *f*_3_- and *f*_4_-statistics that test for admixture were computed with `scikit-allel` using `average_patterson_f3` and `average_patterson_d`. We estimated standard errors with a block jack-knife over 1000 markers (`blen=1000`). When analyzing ancient individuals that were represented as pseudo-haploid genotypes, we analyzed only one allele to avoid an artificial appearance of genetic drift - that could for instance mask a negative *f*_3_ signal of admixture.

### Estimation of *F*_*ST*_-coefficients

To measure pairwise genetic differentiation between two populations, we estimated average pair-wise *F*_ST_ and standard error via block-jackknife over 1000 markers, using `average_patterson_fst` from the package `scikit-allel`. When analyzing ancient individuals that were represented as pseudo-haploid genotypes, we analyzed only one allele to avoid artificial genetic drift. For this analysis, we removed first degree relatives within each population. Another estimator, `average_hudson_fst` gave highly correlated results (*r*^2^ = 0.89), differing mostly for populations with very low sample size (*n* ⩽ 5) and did not change any qualitative conclusions.

### Estimation of admixture proportions with qpAdm

We estimated admixture fractions of a selected target population as well as model consistency for one-, two- or three-source models. We used the framework of Haak *et al.* (2015) as implemented in `qpAdm`, which relates a set of “left” populations (the population of interest and candidate ancestral sources) to a set of “right” populations (diverse out-groups). To test the robustness of our results to the choice of right populations, we ran one analysis with a previously used set of modern populations as outgroup (Haak *et al.*, 2015), and another analysis with a set of ancient Europeans that have been previously used to disentangle divergent strains of ancestry present in Europe (Lazaridis *et al.*, 2017). The full qpAdm results are discussed in Supp. Info. 7.

### Inference of Y haplogroups

To determine the haplotype branch of the Y chromosome of male ancient individuals, we analyzed informative SNPs on the Y-haplotype tree. For reference, we used markers from https://isogg.org/tree/ (Version: 13.238, 2018). We merged this data-set with our set of calls and identified markers available in both to create groups of equivalent markers for sub-haplogroups. Our targeted sequencing approach yielded calls for up to 32,681 such Y-linked markers per individual. As the conventions for naming of haplogroups are subject to change, we annotate them in terms of carrying the derived state at a defining SNP. We analyzed the number of derived and ancestral calls for each informative marker for all ancient Sardinian individuals, and additionally reanalyzed male ancient West Eurasians in our reference data set.

## Data Availability

The raw reads and aligned sequences of the data generated from this study will available through the European Nucleotide Archive (ENA) before publication under accession number [TBD prior to publication]. Processed genotype calls will be posted to the Novembre Lab website and on Data Dryad. The contemporary Sardinia data used to support this study have allele frequency summary data deposited to EGA under accession number EGAS00001002212. The disaggregated individual-level sequence data for 1,577 used in this study is a subset of 2,105 samples (adult volunteers of the SardiNIA cohort longitudinal study) from Sidore et al (2015) and are available from dbGAP under project identifier phs000313 (v4.p2). The remaining individual-level sequence data are from a case-control study of autoimmunity from across Sardinia, and per the obtained consent and local IRB, these data are only available for collaboration by request from the project leader (Francesco Cucca, Consiglio Nazionale delle Ricerche, Italy)

## Supporting information

Supplemental Info Sections 1-2

Supplemental Info Sections 3-8

## Acknowledgements

We thank Maanasa Rhagavan for in-depth feedback on drafts, Anna Di Rienzo and Goncalo Abecasis for helpful discussions, and Magdalena Zoledziewska for useful comments and early assistance. We would like to thank Antje Wissgott, Cäcilia Freund and other members of the wet laboratory and computationally teams at MPI-SHH in Jena. We thank Nadin Rohland, Éadaoin Harney, Shop Mallick, and Alan Cooper for contributing to generating the data for the four samples processed at the Australian Centre for Ancient DNA and in D.R.’s ancient DNA laboratory. We also thank Dan Rice, Chi-chun Liu, and members of the Novembre lab for helpful discussion and feedback. This study was supported in part by the National Science Foundation fellowship DGE-1746045 and the National Institute of General Medical Sciences under training grant award number T32GM007197 to J.H.M. and by funding from the MacArthur Foundation to J.N.. The contribution of D.R. to this study was supported by NSF HOMINID grant BCS-1032255 and by the Howard Hughes Medical Institute. We also thank Alan Cooper for a role in the collection of those 4 samples.

## Author contributions

We annotate author contributions using the CRediT Taxonomy labels (https://casrai.org/credit/). Where multiple individuals serve in the same role, the degree of contribution is specified as ‘lead’, ‘equal’, or ‘supporting’.

- Conceptualization (Design of study) – lead: FC, JN, JK, LL; supporting: CS, CP, DS, JHM, GA
- Investigation (Collection of skeletal samples) – lead: LL, RS; supporting: JB, MGG, CDS, CP (minor contribution from CS, JN)
- Investigation (Ancient DNA isolation and sequencing) – lead: CP, AF; supporting: CDS, JK, DR *, RR
- Data Curation (Data quality control and initial analysis) – lead: JHM, CP; supporting: HR, CS, CC, KD, HA, AO
- Formal Analysis (General population genetics) – lead: JHM, HR; supporting: TAJ
- Writing (original draft preparation) – lead: JHM, HR, JN; supporting: CP, RS, LL, FC
- Writing (review and editing) – input from all authors*
- Supervision – equal: FC, JK, JN
- Funding acquisition – lead: JK, FC, JN; supporting: RS

*: D.R. contributed data for four samples and reviewed the description of the data generation for these samples. As he is also senior author on a separate manuscript that reports data on a non-overlapping set of ancient Sardinians and his group and ours wished to keep the two studies intellectually independent, he did not review any other parts of this manuscript.

## References

1000 Genomes Project Consortium, et al., 2015 A global reference for human genetic variation. Nature 526: 68.

Al-Asadi, H., K. Dey, J. Novembre, and M. Stephens, 2018 Inference and visualization of dna damage patterns using a grade of membership model. bioRxiv : 327684.

Alexander, D. H., J. Novembre, and K. Lange, 2009 Fast model-based estimation of ancestry in unrelated individuals. Genome Research 19: 1655–1664.

Allentoft, M. E., M. Sikora, K.-G. Sjögren, S. Rasmussen, M. Rasmussen, et al., 2015 Population genomics of Bronze Age Eurasia. Nature 522: 167–172.

Ammerman, A. J., and L. L. Cavalli-Sforza, 2014 The Neolithic transition and the genetics of populations in Europe, volume 836. Princeton University Press.

Barbujani, G., and R. R. Sokal, 1990 Zones of sharp genetic change in europe are also linguistic boundaries. Proceedings of the National Academy of Sciences 87: 1816–1819.

Barnett, W. K., 2000 Cardial pottery and the agricultural transition in mediterranean europe. Europe’s first farmers : 93–116.

Calò, C., A. Melis, G. Vona, and I. Piras, 2008 Review synthetic article: Sardinian population (italy): A genetic review. International Journal of Modern Anthropology 1: 39–64.

Cavalli-Sforza, L. L., 2005 The human genome diversity project: past, present and future. Nature Reviews Genetics 6: 333.

Chiang, C. W., J. H. Marcus, C. Sidore, A. Biddanda, H. Al-Asadi, et al., 2018 Genomic history of the Sardinian population. Nature Genetics : 1.

Contu, L., M. Arras, C. Carcassi, G. L. Nasa, and M. Mulargia, 1992 Hla structure of the sardinian population: a haplotype study of 551 families. Tissue Antigens 40: 165–174.

Dabney, J., M. Knapp, I. Glocke, M.-T. Gansauge, A. Weihmann, et al., 2013 Complete mitochondrial genome sequence of a middle pleistocene cave bear reconstructed from ultrashort dna fragments. Proceedings of the National Academy of Sciences : 201314445.

Eaves, I. A., T. R. Merriman, R. A. Barber, S. Nutland, E. Tuomilehto-Wolf, et al., 2000 The genetically isolated populations of finland and sardinia may not be a panacea for linkage disequilibrium mapping of common disease genes. Nature Genetics 25: 320.

Ermini, L., C. Olivieri, E. Rizzi, G. Corti, R. Bonnal, et al., 2008 Complete mitochondrial genome sequence of the Tyrolean Iceman. Current Biology 18: 1687–1693.

Francalacci, P., L. Morelli, A. Angius, R. Berutti, F. Reinier, et al., 2013 Low-pass DNA sequencing of 1200 Sardinians reconstructs European Y-chromosome phylogeny. Science 341: 565–569.

Fu, Q., M. Hajdinjak, O. T. Moldovan, S. Constantin, S. Mallick, et al., 2015 An early modern human from romania with a recent neanderthal ancestor. Nature 524: 216.

Ghirotto, S., S. Mona, A. Benazzo, F. Paparazzo, D. Caramelli, et al., 2009 Inferring genealogical processes from patterns of bronze-age and modern dna variation in sardinia. Molecular biology and evolution 27: 875–886.

Günther, T., C. Valdiosera, H. Malmström, I. Ureña, R. Rodriguez-Varela, et al., 2015 Ancient genomes link early farmers from atapuerca in spain to modern-day basques. Proceedings of the National Academy of Sciences 112: 11917–11922.

Haak, W., I. Lazaridis, N. Patterson, N. Rohland, S. Mallick, et al., 2015 Massive migration from the steppe was a source for Indo-European languages in Europe. Nature 522: 207–211.

Hofmanová, Z., S. Kreutzer, G. Hellenthal, C. Sell, Y. Diekmann, et al., 2016 Early farmers from across Europe directly descended from Neolithic Aegeans. Proceedings of the National Academy of Sciences 113: 6886–6891.

Keller, A., A. Graefen, M. Ball, M. Matzas, V. Boisguerin, et al., 2012 New insights into the tyrolean iceman’s origin and phenotype as inferred by whole-genome sequencing. Nature communications 3: 698.

Korneliussen, T. S., A. Albrechtsen, and R. Nielsen, 2014 Angsd: analysis of next generation sequencing data. BMC bioinformatics 15: 356.

Lai, L., R. H. Tykot, E. Usai, J. F. Beckett, R. Floris, et al., 2013 Diet in the sardinian bronze age: models, collagen isotopic data, issues and perspectives. Préhistoires Méditerranéennes.

Lannou, M. L., 1941 Pâtres et Paysans de la Sardaigne. Tours 8: 364.

Lazaridis, I., A. Mittnik, N. Patterson, S. Mallick, N. Rohland, et al., 2017 Genetic origins of the Minoans and Mycenaeans. Nature 548: 214–218.

Lazaridis, I., D. Nadel, G. Rollefson, D. C. Merrett, N. Rohland, et al., 2016 Genomic insights into the origin of farming in the ancient Near East. Nature 536: 419–424.

Lazaridis, I., N. Patterson, A. Mittnik, G. Renaud, S. Mallick, et al., 2014 Ancient human genomes suggest three ancestral populations for present-day Europeans. Nature 513: 409–413.

Lee, S., F. Zou, and F. A. Wright, 2010 Convergence and prediction of principal component scores in high-dimensional settings. Annals of Statistics 38: 3605.

Lettre, G., and J. N. Hirschhorn, 2015 Small island, big genetic discoveries. Nature Genetics 47: 1224–1225.

Li, H., and R. Durbin, 2009 Fast and accurate short read alignment with burrows–wheeler transform. bioinformatics 25: 1754–1760.

Lipson, M., A. Szécsényi-Nagy, S. Mallick, A. Pósa, B. Stégmár, et al., 2017 Parallel palaeogenomic transects reveal complex genetic history of early European farmers. Nature.

Lugliè, C., 2018 Your path led trough the sea … the emergence of Neolithic in Sardinia and Corsica. Quaternary International 470: 285–300.

Martins, H., F. X. Oms, L. Pereira, A. W. Pike, K. Rowsell, et al., 2015 Radiocarbon dating the beginning of the Neolithic in Iberia: new results, new problems. Journal of Mediterranean Archaeology 28: 105–131.

Mastino, A., 2005 Storia della Sardegna antica, volume 2. Il Maestrale.

Mathieson, I., S. Alpaslan-Roodenberg, C. Posth, A. Szécsényi-Nagy, N. Rohland, et al., 2018 The genomic history of southeastern Europe. Nature 555: 197.

Mathieson, I., I. Lazaridis, N. Rohland, S. Mallick, N. Patterson, et al., 2015 Genomewide patterns of selection in 230 ancient Eurasians. Nature 528: 499–503.

Mathieson, I., S. A. Roodenberg, C. Posth, A. Szécsényi-Nagy, N. Rohland, et al., 2017 The genomic history of Southeastern Europe. bioRxiv : 135616.

Matisoo-Smith, E., A. Gosling, D. Platt, O. Kardailsky, S. Prost, et al., 2018 Ancient mitogenomes of Phoenicians from Sardinia and Lebanon: A story of settlement, integration, and female mobility. PloS one 13: e0190169.

Melis, P., 2002 Un Approdo della costa di Castelsardo, fra etá nuragica e romana. Atti del XIV Congresso L’Africa Romana – Sassari 7-10 dicembre 2000 : 1331–1343.

Meyer, M., and M. Kircher, 2010 Illumina sequencing library preparation for highly multiplexed target capture and sequencing. Cold Spring Harbor Protocols 2010: pdb–prot5448.

Modi, A., F. Tassi, R. R. Susca, S. Vai, E. Rizzi, et al., 2017 Complete mitochondrial sequences from Mesolithic Sardinia. Scientific reports 7: 42869.

Olalde, I., S. Brace, M. E. Allentoft, I. Armit, K. Kristiansen, et al., 2018 The Beaker phenomenon and the genomic transformation of Northwest Europe. Nature 555: 190.

Olalde, I., S. Mallick, N. Patterson, N. Rohland, V. Villalba-Mouco, et al., 2019 The genomic history of the Iberian Peninsula over the past 8000 years. Science 363: 1230–1234.

Olivieri, A., C. Sidore, A. Achilli, A. Angius, C. Posth, et al., 2017 Mitogenome diversity in Sardinians: a genetic window onto an island’s past. Molecular Biology and Evolution 34: 1230–1239.

Ortu, L., 2011 Storia della Sardegna dal Medioevo all’etá contemporanea. Cuec.

Patterson, N., P. Moorjani, Y. Luo, S. Mallick, N. Rohland, et al., 2012 Ancient admixture in human history. Genetics 192: 1065–1093.

Patterson, N., A. L. Price, and D. Reich, 2006 Population structure and eigenanalysis. PLoS Genetics 2: e190.

Peltzer, A., G. Jäger, A. Herbig, A. Seitz, C. Kniep, et al., 2016 Eager: efficient ancient genome reconstruction. Genome Biology 17: 60.

Pickrell, J. K., and D. Reich, 2014 Toward a new history and geography of human genes informed by ancient dna. Trends in Genetics 30: 377–389.

Pritchard, J. K., M. Stephens, and P. Donnelly, 2000 Inference of population structure using multilocus genotype data. Genetics 155: 945–959.

Reich, D., A. L. Price, and N. Patterson, 2008 Principal component analysis of genetic data. Nature Genetics 40: 491.

Renaud, G., V. Slon, A. T. Duggan, and J. Kelso, 2015 Schmutzi: estimation of contamination and endogenous mitochondrial consensus calling for ancient dna. Genome Biology 16: 224.

Rohland, N., E. Harney, S. Mallick, S. Nordenfelt, and D. Reich, 2015 Partial uracil– dna–glycosylase treatment for screening of ancient dna. Philosophical Transactions of the Royal Society B: Biological Sciences 370: 20130624.

Sarno, S., A. Boattini, L. Pagani, M. Sazzini, S. De Fanti, et al., 2017 Ancient and recent admixture layers in Sicily and Southern Italy trace multiple migration routes along the Mediterranean. Scientific reports 7: 1984.

Schubert, M., S. Lindgreen, and L. Orlando, 2016 Adapterremoval v2: rapid adapter trimming, identification, and read merging. BMC research notes 9: 88.

Sidore, C., F. Busonero, A. Maschio, E. Porcu, S. Naitza, et al., 2015 Genome sequencing elucidates Sardinian genetic architecture and augments association analyses for lipid and blood inflammatory markers. Nature Genetics 47: 1272–1281.

Sikora, M., M. L. Carpenter, A. Moreno-Estrada, B. M. Henn, P. A. Underhill, et al., 2014 Population genomic analysis of ancient and modern genomes yields new insights into the genetic ancestry of the Tyrolean Iceman and the genetic structure of Europe. PLoS Genetics 10: e1004353.

Siniscalco, M., L. Bernini, G. Filippi, B. Latte, P. M. Khan, et al., 1966 Population genetics of haemoglobin variants, thalassaemia and glucose-6-phosphate dehydrogenase deficiency, with particular reference to the malaria hypothesis. Bulletin of the World Health Organization 34: 379.

Skeates, R., M. G. Gradoli, and J. Beckett, 2013 The cultural life of caves in Seulo, central Sardinia. Journal of Mediterranean Archaeology 26.

Skoglund, P., H. Malmström, A. Omrak, M. Raghavan, C. Valdiosera, et al., 2014 Genomic diversity and admixture differs for Stone-Age Scandinavian foragers and farmers. Science 344: 747–750.

Skoglund, P., H. Malmström, M. Raghavan, J. Storå, P. Hall, et al., 2012 Origins and genetic legacy of Neolithic farmers and hunter-gatherers in Europe. Science 336: 466–469.

Tykot, R. H., 1996 Obsidian procurement and distribution in the central and western Mediterranean. Journal of Mediterranean Archaeology 9: 39–82.

Vianello, D., F. Sevini, G. Castellani, L. Lomartire, M. Capri, et al., 2013 Haplofind: A new method for high-throughput mt dna haplogroup assignment. Human mutation 34: 1189–1194.

Zalloua, P., C. J. Collins, A. Gosling, S. A. Biagini, B. Costa, et al., 2018 Ancient DNA of Phoenician remains indicates discontinuity in the settlement history of Ibiza. Scientific reports 8: 17567.

Zalloua, P. A., D. E. Platt, M. El Sibai, J. Khalife, N. Makhoul, et al., 2008 Identifying genetic traces of historical expansions: Phoenician footprints in the Mediterranean. The American Journal of Human Genetics 83: 633–642.

Zavattari, P., E. Deidda, M. Whalen, R. Lampis, A. Mulargia, et al., 2000 Major factors influencing linkage disequilibrium by analysis of different chromosome regions in distinct populations: demography, chromosome recombination frequency and selection. Human molecular genetics 9: 2947–2957.

Zilhão, J., 2001 Radiocarbon evidence for maritime pioneer colonization at the origins of farming in west Mediterranean Europe. Proceedings of the national Academy of Sciences 98: 14180–14185.

