## Supplemental Info Sections 1-2 for "Population history from the Neolithic to present on the Mediterranean island of Sardinia: An ancient DNA perspective"

### 1 SITE DESCRIPTIONS FROM THE SAMPLING EFFORT OF LUCA LAI

**LUCA LAI**

A set of skeletal elements, previously excavated from across numerous sites, was organized to form the first major portion of samples in the study and with a focus on sampling from the Neolithic to the end of the Bronze Age.

**SITE: Is Arutas**

The site is a small partially modified natural cave near the seashore and a few miles from the brackish water Cabras Lagoon, in W Sardinia (Cabras municipality, Oristano). Thorough archaeological information regarding the site has never been published; in fact, the context had been looted and the mistaken chronology that was attributed to the skeletal remains for several decades had been inferred from the cultural markers associated with them, interpreted as Late Neolithic-Early Copper Age but as well unpublished (Germanà 1980).

The select remains of 25 individuals were recovered by Prof. Atzeni, in a burial arrangements described as both primary and secondary; further information on the remains, which were analyzed mostly to trace origin of the population, can be found in detailed publications (Germanà 1980, 1982). The general picture that the osteologist drew on the population was that of a group with diverse morphological features, fairly healthy, with a balanced nutrition. Among the associated faunal remains – as well unpublished and apparently lost – several specimens of *Prolagus sardus* are mentioned, and a whale vertebra (Germanà 1995: 55-64).

The remains were then sampled for stable isotopic analyses in 2003 (Lai 2014); in this occasion, besides oxygen isotopes supporting a scenario of high variability, AMS dating disproved the attribution to the Neolithic, yielding a range compatible with Nuragic Late Bronze Age (AA-64824, 3054 ± 55 BP = 3382-3079 cal BP 2σ). The two samples used for aDNA extraction yielded virtually identical Late-to-Final Bronze Age dates (MAMS-26896, 2941 ± 27 BP = 3180-3000 cal BP 2σ and MAMS-26894, 2952 ± 25 BP = 3210-3010 cal BP 2σ), largely overlapping with the previous one.

**SITE: Ingurtosu Mannu**

The human remains that yielded the sample used for aDNA analysis were recovered through excavation by the Soprintendenza per le Provincie di Cagliari e Oristano in 1996, inside the structure of a chambered tomb of the canonical Nuragic type, in the municipality of Donori (S Sardinia), about 20 miles North of Cagliari. Whereas no report of the excavation context has ever been published, the osteological analysis enabled the identification of at least 37 individuals of all ages (Martella et al. 2014) and of specific pathological specimens (Canci et al 2002). Preservation of long bones was so good that stature estimation and the study of stress markers was possible for many individuals, drawing a picture of a group dedicated to intense physical stress, particularly affecting the lower limbs. Tissue preservation turned out to be also very good: most individuals had collagen yields higher than 10%, with peaks over 20% of the original weight.

One AMS determination on bone provided the first absolute indication of chronology, placing the collection in the Nuragic Final Bronze Age (1205-910 cal BC: Martella et al. 2014: 69, no raw date reported). A further date comes from sample MA110, used for aDNA extraction (MAMS-26893, 2941

± 24 BP = 3169-3004 cal BP, 2σ), which resulted slightly earlier but largely overlapping, confirming the Late-Final Bronze Age chronological placement.

**SITE: Cannas di Sotto, t.12**

The site is a vast necropolis of rock-carved tombs, mostly unexcavated, located on a low limestone plateau partially incorporated into the urban area of the city of Carbonia (SW Sardinia). Survey and partial excavation of tomb 12 was carried out in 1983. Only the corridor and one room of the tomb (room A) was brought to light then, which yielded only a preliminary report with site plan and select materials (Santoni&Usai 1995). The remains were not studied, and only a few select cranial fragments from six individuals were sampled for stable isotopic analyses in 2003, which also yielded the first absolute dating (Lai 2009: 318), among which was one that later has been analyzed for aDNA for this project.

New excavation was carried out after 2012, which uncovered archaeological deposits in the inner room, named B, with large amounts of human remains that underwent a preliminary analysis (Salis et al. 2015). Such skeletal remains, largely disarticulated, were recovered in no apparent order in the two rooms, mixed with infiltrated soil, plain pottery, lithic tools, and a few female figurines. Whereas use during the Early Copper Age (ca. 5350-4750 cal BP) was already ascertained in the previous excavation based on pottery style (Santoni &Usai 1995; Melis 2000: 152), a few cultural markers from the last investigation pointed to a longer use of the rock-carved burial, beginning in a Middle-to-Late Neolithic transitional phase (ca. 6150-5750 cal BP).

One AMS date on a sample from the 1983 excavation had already confirmed the chronological placement (AA-64825, 4476 ± 43 BP = 5298-4973 cal BP 2σ: Lai 2009: 318), and the new one from the same batch, contextually obtained as part of the present project, is virtually overlapping (MAMS-26903, 4551 ± 26 BP = 5318-5057 cal BP 2σ).

**SITE: Filigosa, tomb 1**

The site is a necropolis of four rock-carved tombs near Macomer (Nuoro), central-Northwestern Sardinia, a type lasting from the final Middle Neolithic through the Early Copper Age, in some cases reused for depositions until Nuragic times (tomb 4: FoschiNieddu, 1995).More specifically, tomb1, composed of seven rooms with addition of an entrance corridor, did not yield any diagnostic indicators earlier than the Early Copper Age, and was not apparently reused after the Early Copper Age, constituting one of the rare examples of this kind that has survived intact through the 20th century AD. However, it was excavated in 1965 only after being looted, which left behind only ceramic items, the object of an in-depth monograph (FoschiNieddu 1986), and a wealth of bone specimens, but little in terms of stratigraphy. The skeletal remains were not analyzed in any detail, but only used for stature determination by F. Germanà, despite their remarkable physical preservation, probably due to a waterlogged, muddy environment.

Mandibles were sampled for isotopic analyses in 2011, and in such occasion the first AMS date was obtained with funding from the Sardinian Autonomous Region (CRP1_661), supporting the attribution suggested by the associated material culture identifiers (OxA-25337, 4401 ± 32 BP = 5213-4865 cal BP 2σ), and finally a new radiocarbon date is presented in this study, which still confirms the phase of first attribution (MAMS-38276, 4472 ± 25 BP = 5286-4979 cal BP 2σ. All remains are currently undergoing examination according to present-day standards by Dr. C. Rodriguez, doctoral candidate at Universitat Autonoma de Barcelona.

**SITE: S'Iscia 'e sasPiras**

The site is located in Northwestern Sardinia, Usini municipality (Sassari). It consists of a necropolis of three rock-carved tombs. One, perhaps to be identified with tomb 2, was excavated by E. Castaldi in

1966 after looting was reported; it is a typically Nuragic tomb that fulfils the function of chamber tombs in the same time period, although potentially obtained by retouching a previous Neolithic tomb. The remains of over at least 14 individuals were collected and later studied by F. Germanà (1973), who aimed at classifying crania into types to infer ethnicity, and to investigate lifestyle and health. Stress markers on long bones were interpreted as reflections of a very active lifestyle, possibly linked to herding. These remains were later sampled for stable isotopic analyses, and in such occasion one AMS date was obtained with funding from the Sardinian Autonomous Region (CRP1_661), which supported the attribution suggested by the associated material culture identifiers, narrowing it to the Late-Final Bronze Age (OxA-25338, 2918 ± 28 BP = 3159-2971 cal BP 2σ). However, the date yielded by the sample in this study pertains to remains of a previous use of the burial, occurred between the Final Copper and the Early Bronze Age (MAMS-38277, 3794 ± 25 BP = 4244-4091 cal BP 2σ). Considering the commingled conditions of the remains, this date extends the potential chronological range of all undated specimens to well over a millennium, witnessing to the permanence of some specimens from previous burials.

**SITE: S'Orcu 'e Tueri**

The site is a natural cave, located in the Perdasdefogu municipality (Nuoro), a mountainous area of Eastern central Sardinia. Already known by locals, it was discovered to science in 1963, when looting was reported and prompted a salvage recovery of the best-preserved human remains from the floor of the cave, which are curated at the University of Cagliari, Dept. of Life and Environmental Sciences (Maxia 1964). However, looting continued, and more skeletal remains were recovered by the local speleological association Gruppo Grotte Ogliastra, and finally in 2014 all the recoverable remains on the surface were removed in a controlled manner by volunteers coordinated by the Soprintendenza Archeologica Sassari-Nuoro. From these remains, curated at the local museum in Perdasdefogu, samples were removed for the present aDNA investigation.

The cave was used for deposition of the dead mostly in Nuragic times, as no different cultural markers were found, and the early attribution based on physical proximity to a Nuragic tower and settlement is supported by radiocarbon dating. The remains recovered earlier were studied by C. Maxia (1964), whereas those recovered recently were studied by P. Martella as part of her PhD research. One radiocarbon date obtained in the 1990s (Cosseddu et al. 1994) supported the attribution to the Nuragic period (2880 ± 60 BP = 3178-2855 cal BP 2σ). Finally, several dates presented in this study on one hand extended backwards the range of use, still fully within the Nuragic phase (eight dates covering a cal BP 2σ range 3335-2949), whereas one outlier records burial in the cave in times when Carthage controlled Sardinian coasts, but could still pertain to indigenous groups (MAMS-38281, 2255 ± 22 BP = 2343-2161 cal BP 2σ).

**SITE: Serra Crabiles, t.3**

The site is located in Northwestern Sardinia, Sennori municipality (Sassari). It consists of a necropolis of at least four tombs of the *domu de janas* type, rock-carved rooms dating to between the final Middle Neolithic and the Early Copper Age. One of them, tomb 1, yielded human remains which were attributed to the Late Copper Age (Monte Claro culture) based on ceramic sherds association – despite the lack of any reliable stratigraphy; in fact, the finding of a Bell Beaker decorated sherd is also mentioned (Germanà 1980). Tombs 2, 3 and 4 were investigated through excavation led by the Soprintendenza Archeologica Sassari-Nuoro in 1981 (Foschi Nieddu 1984), and in 1993-94 several additional rooms connected with tomb 4 were discovered and excavated (Rovina 1994).

The human remains sampled for aDNA extraction come from tomb 3, which yielded like tomb 1 potsherds attributed to the Monte Claro phase, but also Bell Bell Beaker cultural markers. Radiocarbon dates obtained from the human remains analyzed in this study leave some uncertainty, with chronology covering the interface between the two phases (five dates with a cumulative cal BP 2σ range between 4422 and 4151), but more stringent overlap appears with the Bell Beaker dates from Padru Jossu (Lai 2009: 318), and some unpublished ones from Bingia 'e Monti.

**SITE: Su Crucifissu Mannu, t.16, t.22**

The site is located in Northwestern Sardinia, Porto Torres municipality, bordering with Sassari. It is one of the many reused burial areas surrounding a unique ceremonial site dating to the Late Neolithic-Early Copper Age, consisting in rock-carved tombs composed of several interconnected rooms (Demartis 1998). The necropolis was excavated in different campaigns between 1958 and the early 1970s, with the best-documented tomb being t.16, which yielded at least 13 individuals recognized upon discovery. Most of them were attributed to the Early Bronze Age 1 based on associated material remains and a clear stratigraphy, with the possibility for some individuals from rooms D and especially E of pertaining to the previous Monte Claro or Bell Beaker phase (Late-Final Copper Age) (Ferrarese Ceruti 1976: 191). Most other tombs, including t.22 which the remaining skeletal materials are from, were similarly assigned to the EBA1 based on association with cultural markers. The skeletal remains have been partially studied by F. Germanà (1995: 129 and references therein), but only a small fraction of the data was published, and a complete analysis according to modern standards is still missing. Animal bone remains were also recovered but only a short preliminary report exists on a small assemblage from t. 16 (Ferrarese Ceruti 1976).

The nine radiocarbon dates carried out for the present study largely confirmed the overall attribution of materials from t.16, whereas opened the way to new interpretations for t.22: of the seven for t.16, five were fully compatible with an EBA attribution (cumulative cal BP 2σ range between 4243 and 3894), whereas two (MAMS-38299, 3909 ± 19 BP = cal 4420-4260 BP 2σ, and MAMS-38300, 3880 ± 22 BP = 4411-4245 cal BP 2σ) appear early enough to be potentially assigned to the Bell Beaker phase, which culturally can be considered ancestral to Sardinian EBA.

As concerns the two specimens from t.22, one yielded a date that still belongs to the Bronze Age continuum, but falling mostly within the initial MBA (MAMS-38302, 3421 ± 20 BP = 3808-3612 cal BP 2σ), the other yielded an unexpectedly early date that places it fully within the Late Neolithic (MAMS-38301, 5042 ± 21 BP = 5894-5732 cal BP 2σ), showing that the commingled bone assemblage is the result of multiple phases of burial.

**REFERENCES**

Alessandro Canci, Elisabetta Marini, Giuseppina Mulliri, Elena Usai, Lucia Vacca, Giovanni Floris, Silvana Borgognini Tarli, A Case of Madelung's Deformity in a Skeleton from Nuragic Sardinia. *International Journal of Osteoarchaeology* 12:173–177, 2002.

Giovanni Gesuino Cosseddu, Giovanni Floris, Emanuele Sanna, Verso una revisione dell'inquadramento cronologico e morfometrico delle serie scheletriche paleo-protosarde. I: Craniometria, primi dati. *Rivista di Antropologia* 72:153-162, 1994.

Giovanni Maria Demartis, *Tomba V di Montalè. Necropoli di Su Crucifissu Mannu*. Sassari: Betagamma, 1998.

Maria Luisa Ferrarese Ceruti, La tomba XVI di Su CrocifissuMannu e la cultura di Bonnanaro, *Bullettino di Paletnologia Italiana* 81 (1972-74):113-218, 1976.

Alba Foschi Nieddu, I risultati degli scavi 1981 nella necropoli prenuragica di Serra Crabiles, Sennori (Sassari). In W.H. Waldren, R. Chapman, J. Lewthwaite, and R.C. Kennard (eds.), *The Deya Conference* *of Prehistory. Early Settlement in the Western Mediterranean Islands and the Peripheral Areas*, British Archaeological Reports 229:533-552, 1984.

Franco Germanà, II gruppo umano nuragico di S'Iscia 'e sas Piras (Usini-Sassari) (antropologia e paleopatologia). *Studi Sardi* 23:53-124, 1973.

Franco Germanà, Forme umane preistoriche di Serra Crabiles (Sennori-Sassari) nel contesto antropico paleosardo. In *Atti della XXII Riunione Scientifica dell'Istituto Italiano di Preistoria e Protostoria*, Sardegna centro-settentrionale, 21-27 ottobre 1978:305-330, Istituto Italiano di Preistoria e Protostoria, 1980.

Franco Germanà, I paleosardi di Is Aruttas (Cabras-Oristano). Nota I. *Archivio per l'Antropologia e l'Etnologia* 109-110:343-391, 1980.

Franco Germanà, I paleosardi di Is Aruttas (Cabras-Oristano). Nota II. *Archivio per l'Antropologia e l'Etnologia* 120:233-280, 1982.

Franco Germanà, *L'uomo in Sardegna dal paleolitico all'età nuragica*. Sassari: Carlo Delfino, 1995.

Luca Lai, Il clima nella Sardegna preistorica e protostorica: problemi e nuove prospettive. In *Atti della XLIV Riunione Scientifica dell'Istituto Italiano di Preistoria e Protostoria*, Cagliari, Barumini, Sassari, 23-28 November, 2009, vol. I – Relazioni generali: 313-24, 2009.

Luca Lai, Robert H. Tykot, Jessica F. Beckett, Ornella Fonzo, Elena Usai, Ethan Goddard, David Hollander, Diet in the Sardinian Bronze Age: models, isotopic data, issues and perspectives. *Préhistoires Méditerranéennes* 4:1-19, online. URL :http://pm.revues.org/795, 2014.

Patrizia Martella, Rosalba Floris, Elena Usai, Primi dati osteologici su resti scheletrici provenienti da due tombe della Sardegna meridionale: Ingurtosu Mannu (Donori) e Sa Serra Masì (Siliqua). *Annali dell'Università di Ferrara, Museologia Scientifica e Naturalistica* 10(2):68-73, 2014.

Carlo Maxia, Osservazioni sul materiale scheletrico di una grotta funeraria nuragica a Perdasdefogu. . In *Atti della VIII e IX Riunione Scientifica dell'Istituto Italiano di Preistoria e Protostoria*, Trieste, 19-20 October 1963 – Calabria, 6-8 April 1964:157-163, Istituto Italiano di Preistoria e Protostoria, 1964.

Maria Grazia Melis, *L'età del rame in Sardegna: origine ed evoluzione degli aspetti autoctoni*. Villanova Monteleone: Soter, 2000.

Daniela Rovina, Necropoli preistorica: Sennori – Sassari, Loc. Serra Crabiles. *Bollettino di Archeologia* 43/45:105-106, 1994.

Gianfranca Salis, Felicita Farci, Marco Sarigu, Valeria Pusceddu, Necropoli di Cannas di Sotto, Carbonia. Lo scavo della tomba 12. Notizia preliminare. *Quaderni Soprintendenza Cagliari* 26, online, URL: http://quaderniarcheocaor.beniculturali.it/index.php/quaderni/index, 2015.

Vincenzo Santoni and Luisanna Usai. Domus de janas in località Cannas di Sotto (Carbonia). In V. Santoni (ed.), *Carbonia e il Sulcis: archeologia e territorio*: 53-82. Oristano: S'alvure, 1995.

### 2 SITE AND INDIVIDUAL DESCRIPTIONS FROM THE SEULO CAVES PROJECT

#### ROBIN SKEATES

For an introduction to the Seulo Caves see reference: Skeates et al 2013.

#### SITE: Riparo sotto roccia Su Asedazzu

Cannisoni, Seulo, then Cagliari prov., now South Sardinia prov.

Lat: 39o51’22.10” N  Long: 9 o14’56.68” E

Excavation: 2014 (dir. Robin Skeates)

A small cave and rockshelter, used in successive phases of the Bronze Age as a human burial place and historically as a herder’s shelter. (All body parts are represented, suggesting the primary burial of whole bodies and the later accumulation and dispersal of defleshed bones.)

| aDNA sample ID (small find #) | Archaeological context | Human bone type | C14 lab. code | C14 determination | Calibrated date-range BC (CALIB 7.10) | Period/ culture (after Tykot 1994) | Reference |
|---|---|---|---|---|---|---|---|
| MA87 (134) | Surface find | Juvenile maxilla | SUERC-38110 | 2865±35 BP | 1110–981 cal BC (68.3 %) 1187–923 cal BC (95.4 %) | Final Bronze Age (Nuragic III) | Skeates et al. 2013: 105; Oliveri et al. 2017: Tab. S7 |
| SUA003 (307) | Context 1, Grid Square 9 | Petrous portion of skull | MAMS-28656 | 2953±27 BP | 1211- 1125 cal BC (68.3 %) 1257-1056 cal BC (95.4 %) | Late Bronze Age (Nuragic II) | This publication |
| SUA001 (301) | Context 1, Grid Square 5 | Petrous portion of skull | MAMS-28654 | 3060±28 BP | 1388-1279 cal BC (68.3 %) 1409-1233 cal BC (95.4 %) | Middle Bronze Age (Nuragic I) | This publication |
| MA78 (102) | Context 3, Grid Square 12 | 1st molar extracted from a child's mandible | MAMS-26901 | 3658±26 BP | 2124–1977 cal BC (68.3%) 2134–1949 cal BC (95.4%) | Early Bronze Age (Bonnanaro A) | Oliveri et al. 2017: Tab. S7 |
| SUA002 (303) | Context 2, Grid Square 3 | Petrous portion of skull | MAMS-28655 | 3732±30 BP | 2198-2048 cal BC (68.3%) 2205-2033 cal BC (95.4%) | Early Bronze Age (Bonnanaro A) | This publication |
| MA88 (136) | Context 5, Grid Square 9 | 2nd molar, extracted from the left portion of a female | MAMS-26902 | 3794±34 BP | 2286–2150 cal BC (68.3%) | Early Bronze Age (Bonnanaro A) | Oliveri et al. 2017: Tab. S7 |

| | | | | | |
|---|---|---|---|---|---|
| | | adult mandible | | | 2344–2062 cal BC (95.4%) |

**SITE: Riparo sotto roccia Su Cannisoni 1**

Cannisoni, Seulo, then Cagliari prov., now South Sardinia prov.

Lat: 39 o 51’ 38.45” N  Long: 9 o 15’ 9.08” E

Excavation: 2009 (dir. Robin Skeates)

A large rock-shelter with a Middle Bronze Age secondary burial deposit covered by a cairn, placed below a small natural spring. (One of the human vertebrae has been matched to another from the nearby burial cave of Sa Grutta ‘e is Bittuleris, from where it was probably obtained.)

| | | | | | | | |
|---|---|---|---|---|---|---|---|
| MA82/SC1004 (117) | Context 2, Grid Sq 5 | Skull fragment | OxA-22194 | 3220±28 BP | 1508-1449 cal BC (68.3 %)<br><br>1601-1426 cal BC (95.4 %) | Middle Bronze Age (Nuragic I) | Skeates et al. 2013: 105; Oliveri et al. 2017: Tab. S7 |

**SITE: Riparo sotto roccia Su Cannisoni 2**

Cannisoni, Seulo, then Cagliari prov., now South Sardinia prov.

Lat: 39 o 51’ 37.2492” N  Long: 9 o 15’ 12.276” E

Survey: 2009 (dir. Robin Skeates)

A rock-shelter used in the Early Bronze Age as a human burial place.

| | | | | | | | |
|---|---|---|---|---|---|---|---|
| MA81/SC1003 (116) | Surface find | Proximal left ulna | SUERC-38111 | 3555±35 BP | 1952-1783 cal BC (68.3 %)<br><br>2015-1771 cal BC (95.4 %) | Early Bronze Age (Nuragic I) | Skeates et al. 2013: 105; Oliveri et al. 2017: Tab. S7 |

**SITE: Su Stampu Erdi**

Tonnulù, Seulo, then Cagliari prov., now South Sardinia prov.

Lat: 39 o 52’ N  Long: 9 o 12’ E

Survey: 2009 (dir. Robin Skeates)

A cave complex with two entrances, corridors and speleothems, used in the Early and Middle phases of the Bronze Age as a human burial place.

| | | | | | | | |
|---|---|---|---|---|---|---|---|
| (-)No id # published | Surface find | Human bone | Beta-37705 | 3190±80 BP | 1605-1323 cal BC (68.3 %)<br><br>1643-1262 cal BC (95.4 %) | Middle Bronze Age (Nuragic I) | Sanna et al. 1999: 244 |
| MA85/ SE1011 (132) | Surface find | Tibia | SUERC-38111 | 3579±27 BP | 1956-1890 cal BC (68.3 %)<br><br>2023-1880 cal BC (95.4 %) | Early Bronze Age (Bonnanaro A) | Skeates et al. 2013: 104; Oliveri et al. 2017: Tab. S7 |

**SITE: Sa Forada de Gastea / Grotta Gastea**

Monte Gastea, Seulo, then Cagliari prov., now South Sardinia prov.

Lat: 39 º 51' N  Long: 9 º 12' E

Survey: 2009 (dir. Robin Skeates)

A small cave used in the Early Bronze Age as a human burial place.

| | | | | | | | |
|---|---|---|---|---|---|---|---|
| MA86/ MG1012 (133) | Surface find | Adult right fibula | OxA-22650 | 3647±29 BP | 2114-1959 cal BC (68.3 %)<br><br>2134-1937 cal BC (95.4 %) | Early Bronze Age (Bonnanaro A) | Skeates et al. 2013: 103; Oliveri et al. 2017: Tab. S7 |

**SITE:Su Grutta 'e is Bittuleris / Sa Omu 'e is Ossus**

Cannisoni, Seulo, then Cagliari prov., now South Sardinia prov.

Lat: 39 º 51' 38.46" N  Long: 0 º 15' 9.36" E

Excavation: 2009 (dir. Robin Skeates)

A small cave used in the Middle Bronze Age as a human burial place. (Osteological study of the human remains indicates successive primary inhumations of adults and children, males and females, and later disturbance and fragmentation of their bones.)

| | | | | | | | |
|---|---|---|---|---|---|---|---|
| (175) | Context 3, Grid Sq 4, Spit 2 | Adult longbone | OxA-22193 | 3398±26 BP | 1740-1661 cal BC 1749-1629 (68.3 %)<br><br>cal BC (95.4 %) | Middle Bronze Age (Bonnanaro B) | Skeates et al. 2013: 105 |
| ISB001 (1009) | Surface find | Petrous portion of skull | MAMS-28658 | 3460±29 BP | 1873-1698 cal BC (68.3 %)<br><br>1880-1692 cal BC (95.4 %) | Middle Bronze Age (Bonnanaro B) | This publication |
| S1249 (711) | Grid Square 4, Context 1 | tooth | NA | NA | NA | Middle Bronze Age (Bonnanaro B) | This publication (*aDNA by Haak/Reich) |
| S1250 (723) SuB7 | Grid Square 4, Context 1 | tooth | NA | NA | NA | Middle Bronze Age (Bonnanaro B) | This publication (*aDNA by Haak/Reich) |
| S1252 (737) | Grid Square 4, Context 1 | tooth | NA | NA | NA | Middle Bronze Age (Bonnanaro B) | This publication (*aDNA by Haak/Reich) |

| | | | | | | | |
|---|---|---|---|---|---|---|---|
| S1253 (738) | Grid Square 4, Context 1 | tooth | NA | NA | NA | Middle Bronze Age (Bonnanaro B) | This publication (*aDNA by Haak/Reich) |

**SITE: Grutta I de Longu Fresu**

Foresta di Addoli, Seulo, then Cagliari prov., now South Sardinia prov.

Lat: 39 º 51' 6.25" N  Long: 9 º 16' 13.32" E

Excavation: 2009 (dir. Robin Skeates)

A small cave comprising a tunnel with lateral niches and former springs (plus a hole leading down to lower parallel tunnel), used in the Middle Neolithic as a cult cave, with human remains deposited at its innermost end. (Adult and child remains, probably originally deposited as primary burials then later disturbed.) A Middle Bronze Age mortuary phase is also indicated by a more recent radiocarbon determination on human bone.

| | | | | | | | |
|---|---|---|---|---|---|---|---|
| LF1001 (-) | Surface find | Female left temporal with petrous | MAMS-28657 | 3509±28 BP | 1885-1774 cal BC (68.3 %)<br><br>1910-1749 cal BC (95.4 %) | Middle Bronze Age (Bonnanaro B) | Posth |
| MA79/ LF1002 (-) | Surface find in hole at back of cave | Juvenile female tibia | OxA-22195 | 5258±34 BP | 4224-3991 cal BC 4229-3981 (68.3 %)<br><br>cal BC (95.4 %) | Middle Neolithic (Bonu Ighinu) | Skeates et al. 2013: 104; Oliveri et al. 2017: Tab. S7 |
| (-) No id # published | Surface find | Adult skull | OxA-X-2236-44 | 5315±36 BP | 4231-4059 cal BC (68.3 %)<br><br>4257-4042 cal BC (95.4 %) | Middle Neolithic (Bonu Ighinu) | Gradoli and Meaden 2011: 221 |
| (1) No id # published | Context 2, Grid Sq 3, Spit 1 | Adult skull | OxA-22196 | 5354±34 BP | 4315-4074 cal BC 4324-4053 (68.3 %)<br><br>cal BC (95.4 %) | Middle Neolithic (Bonu Ighinu) | Skeates et al. 2013: 104 |

### REFERENCES

Gradoli, M.G. and Meaden, T. 2011. Underworld and Neolithic rituality: the rock art of the Su Longu Fresu cave in central Sardinia. In: E. Anati, ed. *Art and Communication in Pre-Literate Societies*. Capo di Ponte: Centro Camuno di Studi Preistorici, pp. 220-224.

Olivieri, A., Sidore, C., Achilli, A., Angius, A., Posth, C., Furtwängler, A., Brandini, S., Rosario Capodiferro, M., Gandini, F., Zoledziewska, M., Pitzalis, M., Maschio, A., Busonero, F., Lai, L., Skeates, R., Gradoli, M.G., Beckett, J., Marongiu, M., Mazzarello, V., Marongiu, P., Rubino, S., Rito, T., Macaulay, V., Semino, O., Pala, M., Abecasis, G.R., Schlessinger, D., Conde-Sousa, E., Soares, P., Richards, M.B., Cucca, F. and Torroni, A., 2017. Mitogenome diversity in Sardinians: a genetic window onto an island's past'. *Molecular Biology and Evolution*, 34: 1230–1239.

Sanna, E., Liguori, A., Fagioli, M.B. and Floris, G., 1999. Verso una revisione dell'inquadrimento cronologico e morfometrico delle serie scheletriche paleo-protosarde. II: cranometria, ulteriori aggiornamenti. *Archivio per l'Antropologia e l'Etnologia*, 79: 239-250.

Skeates, R., Gradoli, M.G. and Beckett, J., 2013. The cultural life of caves in Seulo, central Sardinia. *Journal of Mediterranean Archaeology*, 26: 97-126.

Tykot, R.H., 1994. Radiocarbon dating and absolute chronology in Sardinia and Corsica. In: R. Skeates and R. Whitehouse, eds. *Radiocarbon Dating and Italian Prehistory*. London: The British School at Rome and Accordia Research Centre, pp. 115-145.
