## Supplemental Info Sections 3-8 for "Population history from the Neolithic to present on the Mediterranean island of Sardinia: An ancient DNA perspective"

March 21, 2019

NOTE: Supplemental Information Sections 1 and 2 (containing the archaeological collection descriptions) are in a separate document.

#### 3 Validating quality and contamination of aDNA

**Joseph H. Marcus, Kushal Dey, Hussein Al-asadi, Harald Ringbauer, Cosimo Posth**

##### Postmortem Damage Filtering

Individuals with high levels of mtDNA or X chromosome based contamination estimates were removed in our main analysis. However, population genetic analyses could still be affected by more subtle modern contamination. To assess this possibility, we filter out reads that do not show a signature of post-mortem damage (pmd), as reads that carry a damage signature are less likely to be introduced by modern contamination (Skoglund et al., 2014). We used `pmdtools` (`https://github.com/pontussk/PMDtools`) to compute a likelihood-based damage score for each read and subsequently removed reads which showed little evidence of being damaged. We then generated pseudo-haploid genotype calls on these "pmd-filtered" individuals and projected them onto the PCs computed in the modern West-Eurasian individuals from the Human Origins dataset, as described in the Materials and Methods. We corrected for the regression towards the mean effect in high-dimensional PCA using a simple jackknife estimator (see Supp. Info. 8).

We found that all the samples we analyzed in the main results and that have enough covered SNPs after pmd filtering show little difference between the pmd-filtered and corresponding unfiltered PC scores (Fig. 1). This observation supports our sample filtering criteria, and as such our population genetic analyses are unlikely to be strongly affected by contamination. A few individuals had too few covered SNPs after pmd filtering to make accurate predictions of their ancestry, which possibly explains larger observed differences between the pmd-filtered and the original projection.

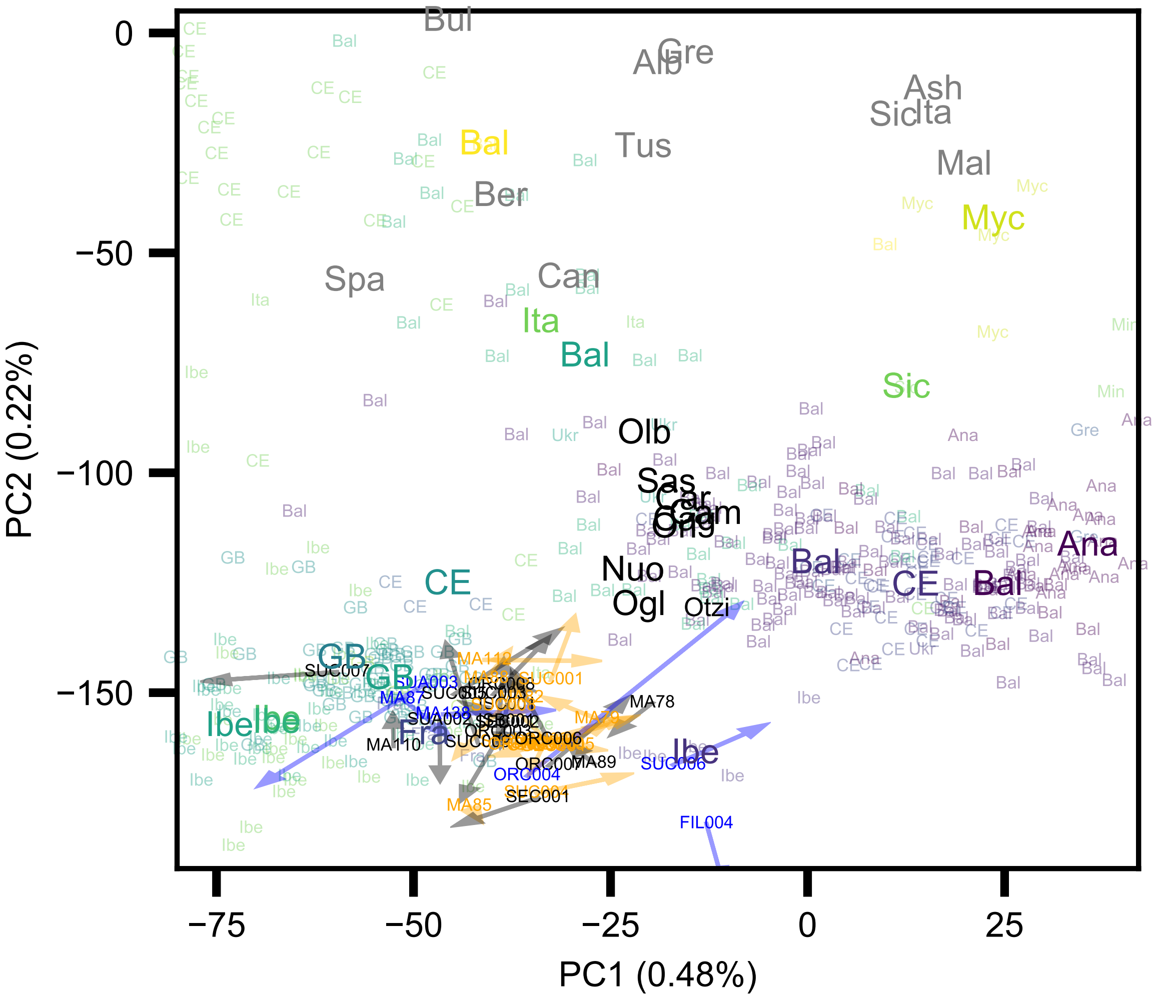


Supplementary Figure 1: **Impact of PMD filtering on PCA projections of ancient individuals.** The figure shows a visualization of PC1 and PC2 computed on modern individuals and projecting ancient individuals from our study, alongside ancients from previously published literature. Each arrow represents an ancient Sardinian individual, where the head of the arrow is the "pmd-filtered" projection and the tail is the "non-pmd-filtered" projection. We color each arrow given the following criteria: black for male individuals that had low X-based contamination estimates (<=0.05); orange for male individuals with high X-based contamination estimates (>0.05) or females; and blue for any remaining individuals with less than 35 thousand covered SNPs after PMD filtering.

#### aRchaic

We estimate DNA damage profiles using `aRchaic` (Al-Asadi et al., 2018). In the `aRchaic` model, mismatches are counted across all of an individual's sequence reads with corresponding measured features, including mismatch type, mismatch position on the read, flanking reference nucleotides, strand orientation. Each mismatch is modeled as originating from a mixture of $K$ profiles defined by these features and adaptively learned during inference via an EM algorithm. Maximum-likelihood estimates of mixture proportions for each individual are then displayed on a stacked bar chart where each row is a different sample and each colored bar represents the proportion of the $i$th individual's mismatches coming from the $k$th mismatch profile 2. When we apply `aRchaic` to our data we observe the typical pattern representative of ancient DNA, an enrichment of cytosine to thymine mismatches occurring preferentially at the ends of the read (Ginolhac et al., 2011; Jónsson et al., 2013). This observations helps to authenticate our data as being truly ancient. Further, we observe that samples treated with a protocol, UDGhalf treatment, to partially remove some of this damage signature from each sample, cluster distinctly from both untreated ancient and modern individuals.

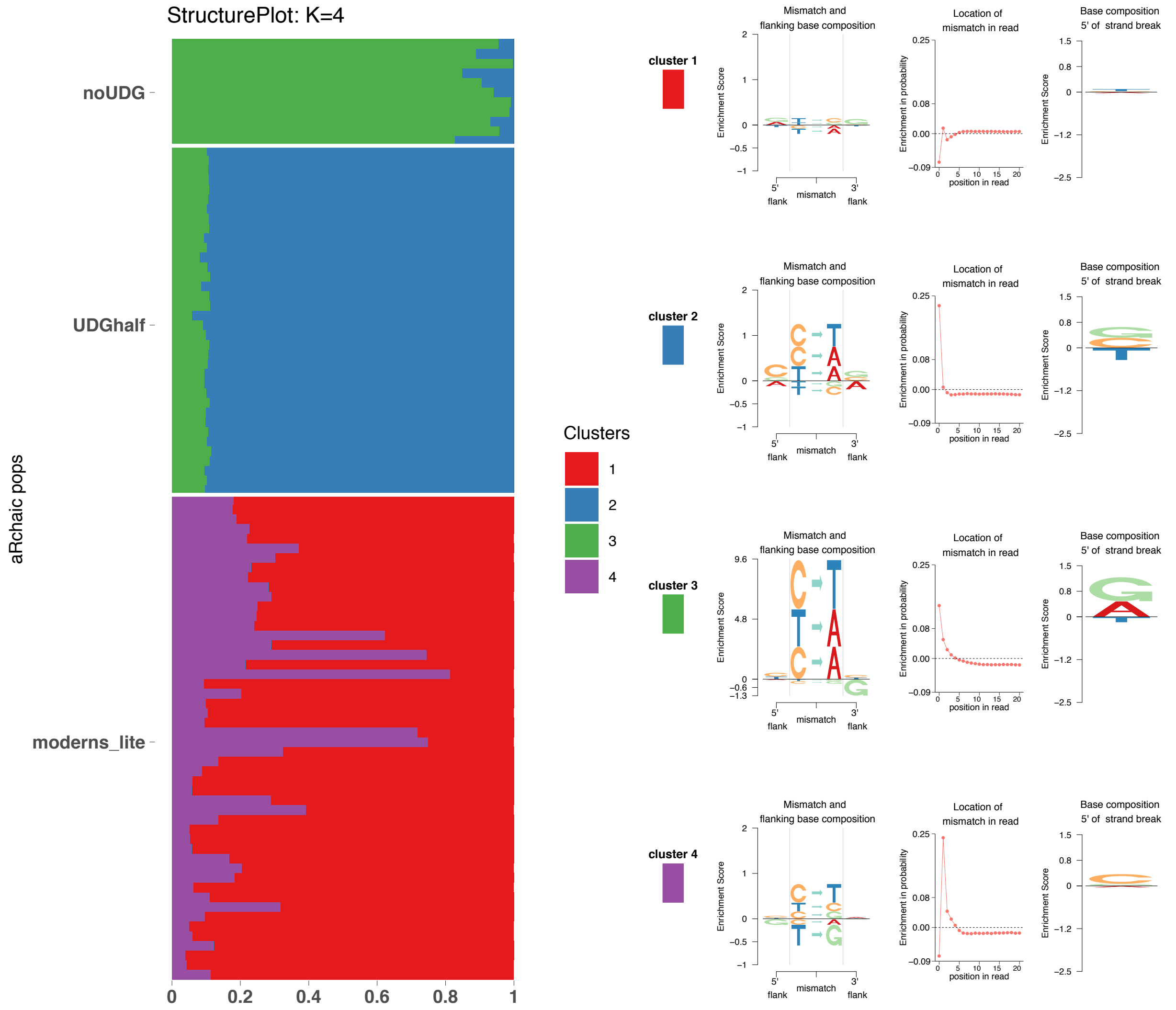


Supplementary Figure 2: **Results of the package `aRchaic` for clustering mismatch profiles in sequence read libaries.** On the left we plot a stacked bar-chart where each row represents a `bam` file and the colored portions of each bar represent the mixture proportion for a given cluster. As we can see each `bam` file's mixture proportions must be non-negative and sum to one. On the right we display representations of the inferred latent variables that define each cluster. The left most plot displays the enrichment or depletion of different mismatch types, the middle plot displays the enrichment probability of observing a mismatch at a particular position along the read, and finally the right hand plot displays an enrichment score for the mismatch type observed at a strand break on the 5' end of the fragment. All together these plots visualize both how the `bam` files are loaded on to each cluster as well as what defines each cluster.

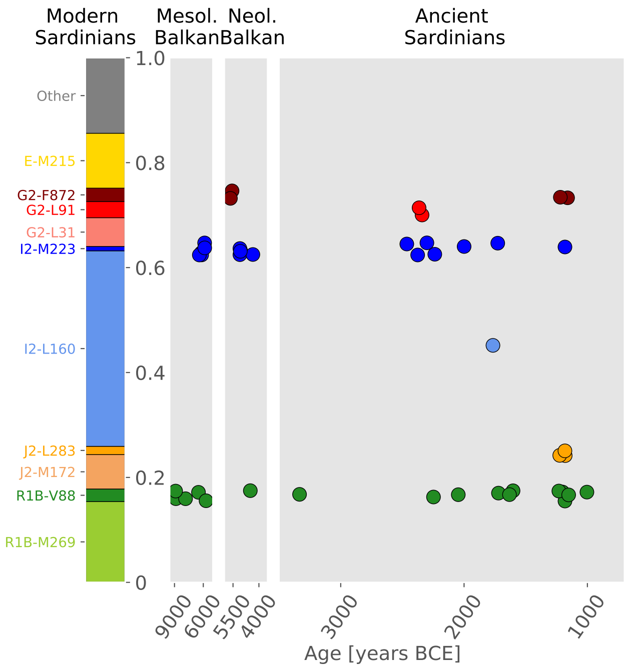


Supplementary Figure 3: **A summary of Y chromosome variation in ancient Sardinians.** On the left we plot a stacked bar-chart representing all major present-day haplogroups, as well as all haplogroups found in at least one ancient Sardinian.The relative frequencies are based on haplogroup abundances in 1,204 present-day Sardinian males reported in Francalacci et al. (2013). In the grey panels, each point corresponds to an ancient sample of a particular age (x-axis) and Y haplotype (y-axis, haplogroup denoted by alignment with modern haplogroups on the left, with jitter added to avoid overlap among individuals of the same age / haplotype configuration). The ancient Balkan data originate from the ancient reference panel (and we include ancient individuals from Hungary in the broader Balkan definition to increase sample number).

### 4 Analysis of ancient Y haplogroups

**Harald Ringbauer**

We were able to assign Y haplogroup designations for 25 ancient Sardinian males (Sup. Mat. 1A and B, Materials and Methods). Here we discuss some of the details and implications of our findings in fuller detail.

#### Comparison to present-day Sardinia Y haplogroup variation

Supp. Fig. 3 presents an overview of the results. Some caution is required in interpretation as we observed substantial clustering of haplogroups within sample site (Sup. Mat. 1B) suggesting some sites may not be random samples of the population. That said, overall there is a notable shift between present-day and ancient Sardinia. The "Sardinian" haplogroup I2-M26, carried by about 35% present-day Sardinian males and rare elsewhere (Francalacci et al., 2013), is found in only one ancient Sardinian sample from the early Middle Bronze Age. The R1b-M269 haplogroup, dominant in present-day continental Iberia after having arrived there with the Bronze Age Indo-European expansion (Olalde et al., 2018) and at 15% frequency in modern Sardinians (Francalacci et al., 2013), is absent throughout our transect from the Neolithic to Nuragic in Sardinia. Similarly, the haplogroup E-M215, a major present-day Sardinian haplogroup (10%) that is prevalent in present-day Northern Africa, was not identified among the ancient individuals we sample. We also detected signals of continuity. The Y haplogroups G2-L91 and R1b-V88, which were previously identified to have major Sardinia-specific sub-clades based on present-day Sardinian variation (Francalacci et al., 2013), were all identified in at least one ancient Sardinian male (Fig. 3).

| Branch | Equiv. Marker | # of Markers | Present-day Distribution |
|---|---|---|---|
| R1b | M343 | 13 | West Eurasia, North Africa |
| R1b1 | L754 | 15 | West Eurasia, North Africa |
| R1b1a1b | M269 | 76 | Western Europe |
| R1b1b | V88 | 37 | Sahel Zone, Sardinia |
| R1b1b2a | V2197 | 10 | Sahel Zone, Sardinia |
| R1b1b2a1 | V35 | 11 | Sardinia |
| R1b1b2a2a | V1944 | 10 | Sahel Zone |

Table 1: R1b subclades with markers on the 1240k target. For each subclade, we show the defining marker, the number of equivalent markers that intersect the 1240k capture panel, as well as its broad present-day geographic distribution.

#### R1b haplogroups in ancient Sardinians

The 1240k capture read data allowed us to call several R1b subclades in ancient individuals (see Supp. Tab. 1 for overview of available markers). While R1b-M269 was absent from our sample of ancient Sardinians, we detected R1b-V88 equivalent markers in 10 out of 25 ancient Sardinian males with robust Y haplogroup calls. Two additional males carried derived markers of the clade R, but we could not identify more refined subclades due to their very low coverage (Supp. Mat. 1B). The ancient geographic distribution of R1b-V88 haplogroups is particularly concentrated in the Seulo caves site and the South of the island (Supp. Mat. 1B).

At present, R1b-V88 is prevalent in central Africans, at low frequency in present-day Sardinians, and extremely rare in the rest of Europe (D'Atanasio et al., 2018). By inspecting our reference panel of western Eurasian ancient individuals, we identified R1b-V88 markers in 10 mainland European ancient samples (Fig. 4), all dating to before the Steppe expansion (> 3k years BCE). Two very basal R1b-V88 (with several markers still in the ancestral state) appear in Serbian HGs as old as 9,000 BCE (Fig. 5), which supports a Mesolithic origin of the R1b-V88 clade in or near this broad region. The haplotype appears to have become associated with the Mediterranean Neolithic expansion - as it is absent in early and middle Neolithic central Europe, but found in an individual buried at the Els Trocs site in the Pyrenees (modern Aragon, Spain)), dated 5,178-5,066 BCE (Haak et al., 2015) and in nine Sardinians. Interestingly, markers of the R1b-V88 subclade R1b-V2197, which is at present-day found in present Sardinians and most African R1b-V88 carriers, are derived only in the Els Trocs individual and two ancient Sardinian individuals (MA89, 3,370-3,110 BCE, MA110 1,220-1,050 BCE) (Fig. 5). MA110 additionally carries derived markers of the R1b-V2197 subclade R1b-V35, which is at present-day almost exclusively found in Sardinians (D'Atanasio et al., 2018).

This configuration suggests that the V88 branch first appeared in eastern Europe, mixed into Early European farmer individuals (after putatively sex-biased admixture (Mathieson et al., 2018)), and then spread with EEF to the western Mediterranean. Individuals carrying an apparently basal V88 haplotype in Mesolithic Balkans and across Neolithic Europe provide evidence against a previously suggested central-west African origin of V88 (González et al., 2013). A west Eurasian R1b-V88 origin is further supported by a recent phylogenetic analysis

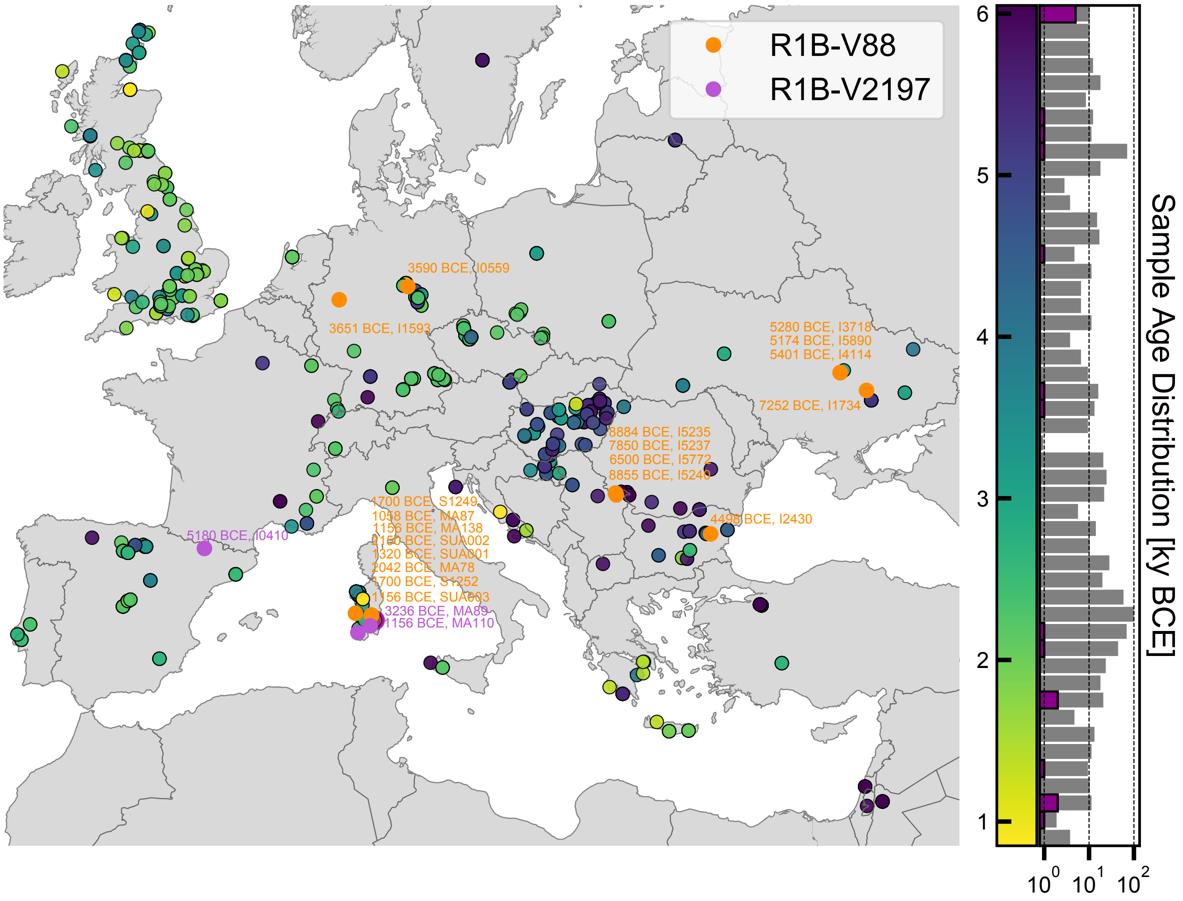


Supplementary Figure 4: **Geographic and temporal distribution of R1b-V88 Y-haplotypes in ancient European samples.** We plot the geographic position of all ancient samples inferred to carry R1b-V88 equivalent markers. Dates are given as years BCE (means of calibrated $2\sigma$ radio-carbon dates). Multiple V88 individuals with similar geographic positions are vertically stacked. We additionally color-code the status of the R1b-V88 subclade R1b-V2197, which is found in most present-day African R1b-V88 carriers.

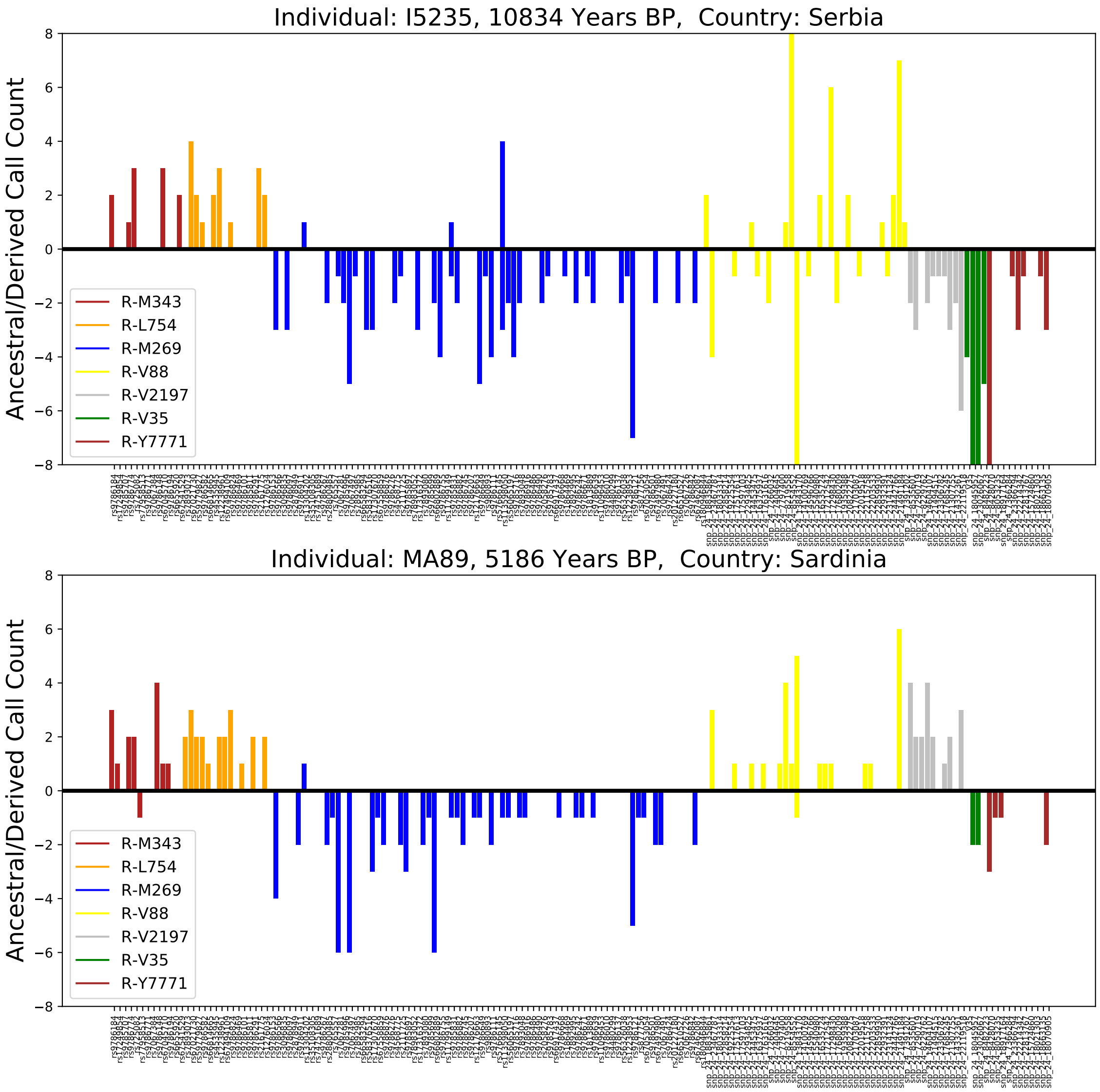


Supplementary Figure 5: **Read data summary for R haplogroup mutations of a Copper Age Sardinian (bottom panel) and a HG individual from Serbia (top panel).** The figure depicts for each marker along the x-axis, the number of calls for the ancestral (downwards) as well as derived alleles (upwards). Both individuals carry reads for the R1b-M269 markers in the ancestral state and for several derived V88 markers. The almost 11-thousand-year-old sample also has several of the equivalent R1b-V88 markers in the ancestral state, and this particular Sardinian has derived markers for the V88 subclade R1b-2197 (similar to the Neolithic Els Trocs sample from the Pyrenees).

that puts modern Sardinian carrier haplotypes basal to the African R1b-V88 haplotypes (D'Atanasio et al., 2018). The putative coalescence times between the Sardinian and African branches inferred there fall into the Neolithic Subpluvial ("green Sahara", about 7,000 to 3,000 years BCE). Previous observations of autosomal traces of Holocene admixture with Eurasians for several Chadic populations (Haber et al., 2016) provide further support for a speculative hypothesis that at least some amounts of EEF ancestry crossed the Sahara southwards. Genetic analysis of Neolithic human remains in the Sahara from the Neolithic Subpluvial would provide further insights into the timing and specific route of this haplogroup into Africa - and whether it was associated with a maritime wave of Cardial Neolithic along Western Mediterranean coasts (Zilhão, 2001), as well as subsequent movement across the Sahara (D'Atanasio et al., 2018; Fregel et al., 2018).

Overall, our analysis provides evidence that R1b-V88 traces back to eastern European Mesolithic hunter gatherers and spread with the Neolithic expansion in Iberia and Sardinia. These results reinforce that the geographic history of a Y-chromosome haplotype can be complex, and modern day spatial distributions need not reflect the initial spread. Broader sampling in Neolithic Mediterranean sites, including the North African coast, will hopefully provide new insights to further refine the origin and migrations of R1b-V88.

### 5 Pairwise similarity statistics

**Harald Ringbauer, Joseph H. Marcus**

#### Measures of pairwise genetic differentiation.

Supp. Fig. 6 and Supp. Fig. 7 depict the matrix of genetic similarities calculated using $f_3$-outgroup statistics and $F_{ST}$ as described in the main text (Materials and Methods). Numerical values are reported in Supp. Mat. 2A and B.

#### Pairwise Relatedness.

To identify close relatives within our dataset of ancient Sardinians, we directly assessed pairwise relatedness. We first filtered to markers that have calls for at least half of the 43 ancient Sardinians, and within those for markers with (pseudo-haploid) minor allele frequency (MAF) $> 0.2$. For this set of $n = 285,476$ markers, we calculated pairwise correlation of allele frequencies (relatedness) between individuals $i$ and $j$:

$$f(i,j) = \frac{(p_i - \bar{p}) \cdot (p_j - \bar{p})}{\bar{p} \cdot (1 - \bar{p})}, \tag{1}$$

averaged over all markers with available (pseudo-haploid) calls for both individuals. Allele frequency means ($\bar{p}$) were calculated from the full set of ancient Sardinians. We then subtracted the mean of this statistic to account for the slight bias introduced via calculating the mean $\bar{p}$ from the same sample.

Using this basic approach enabled us to identify a single pair of first-degree of relatives (expected $f = 0.25$). The remainder of ancient Sardinian samples likely do not contain any first or second degree relatives (Supp. Fig. 8).

This single pair consists of a female and male sample, `SUC002` and `SUC003`, both sampled from the Su Crucifissu Mannu site. The broadly uniform value of estimated $f$ around 0.25 throughout the genome (Supp. Fig. 8) suggests that these two samples are a parent-offspring pair, as full siblings would vary between $f = 0$, 0.25 and 0.5, depending on whether 0, 1 or 2 allele were co-inherited. Both samples had identical mtDNA haplogroup J1c3, providing some evidence that these pair of samples represent a mother and son.

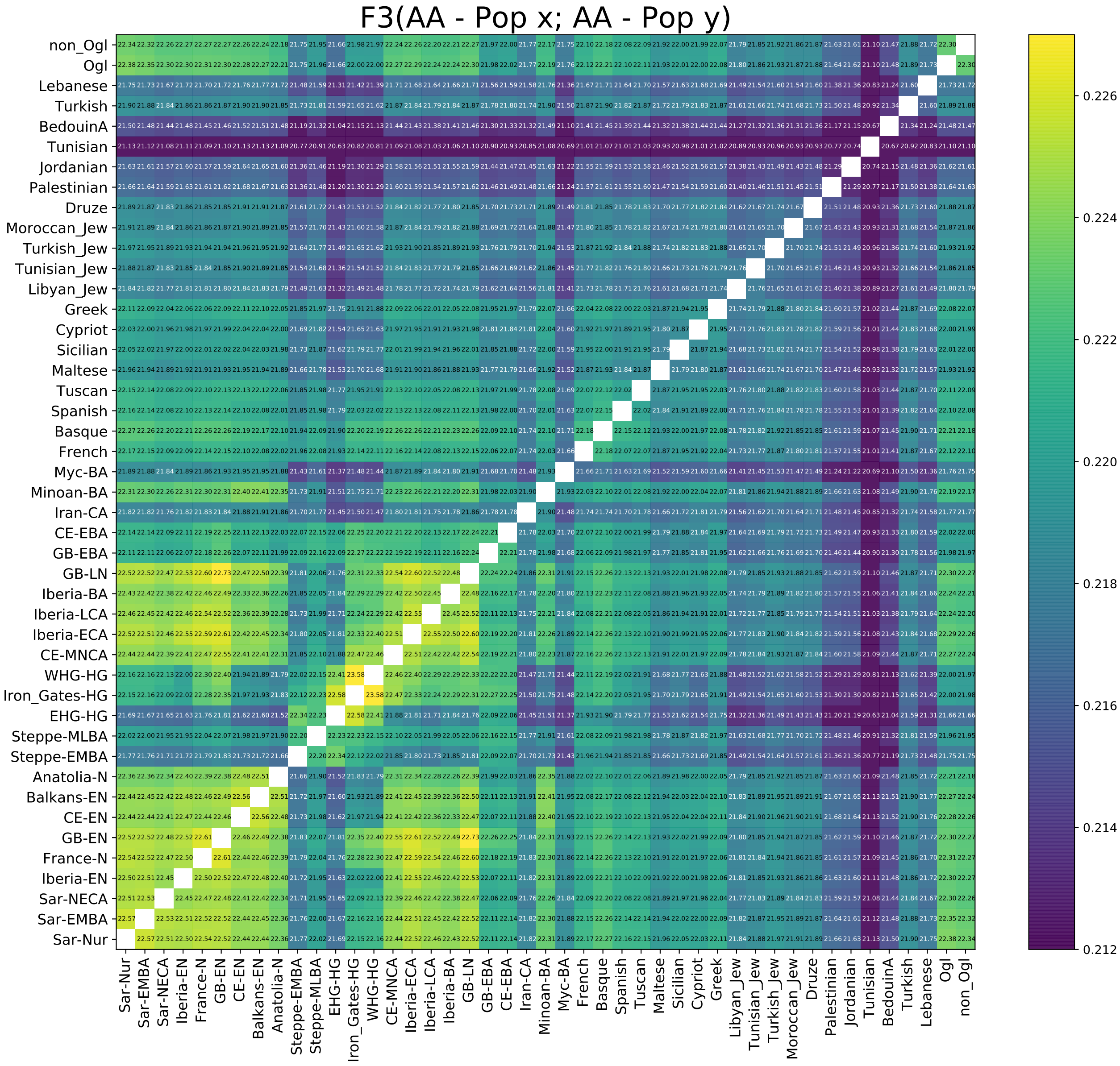


Supplementary Figure 6: **Matrix of f3-outgroup "shared genetic drift" metrics of pairwise similarity.** Populations are ordered broadly by period and geography (See Sup. Mat. 1G for legend to abbreviations).

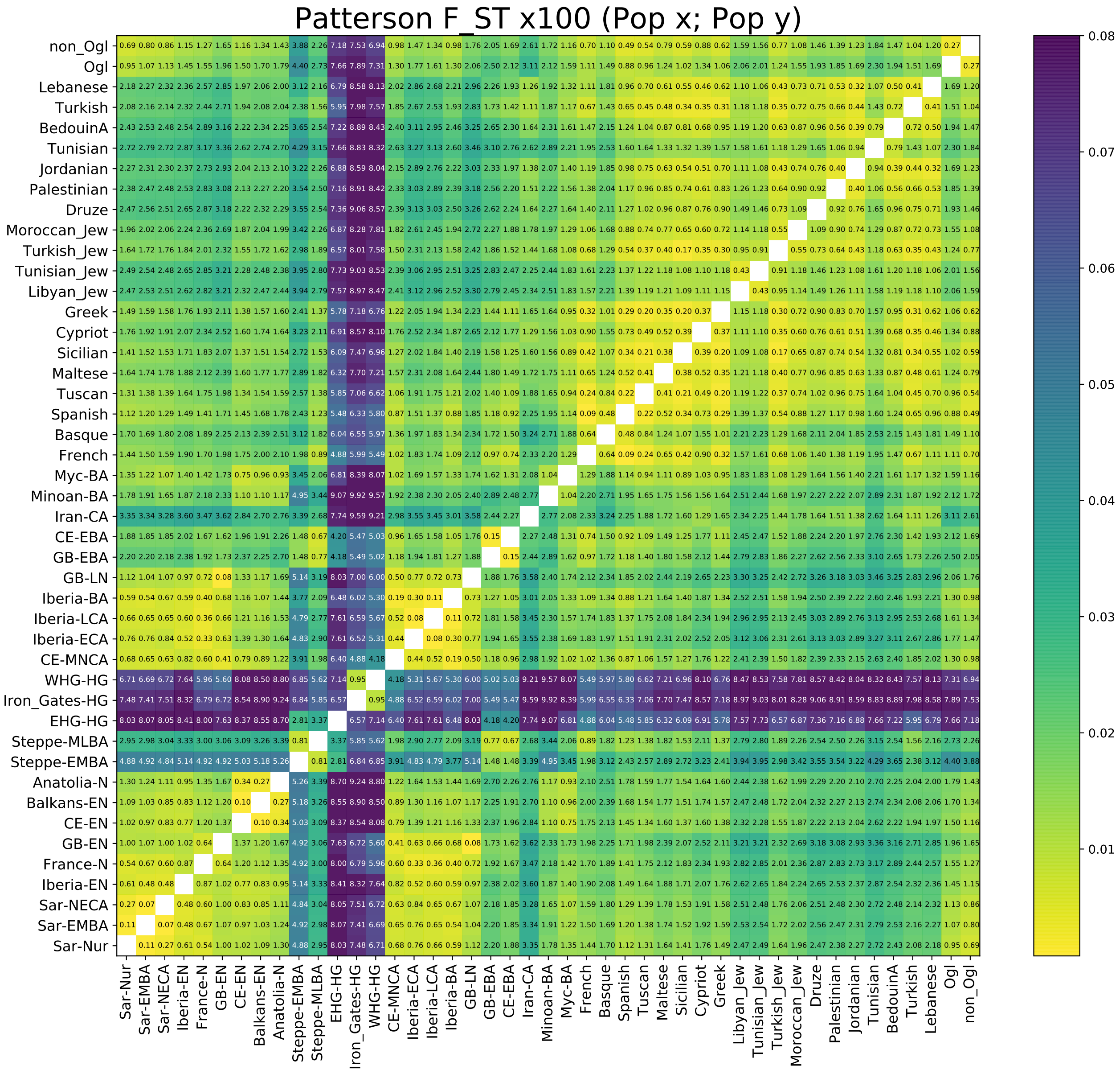


Supplementary Figure 7: **Matrix of $F_{ST}$ metrics of pairwise differentiation**. Populations are ordered broadly by period and geography (See Sup. Mat. 1G for legend to abbreviations).

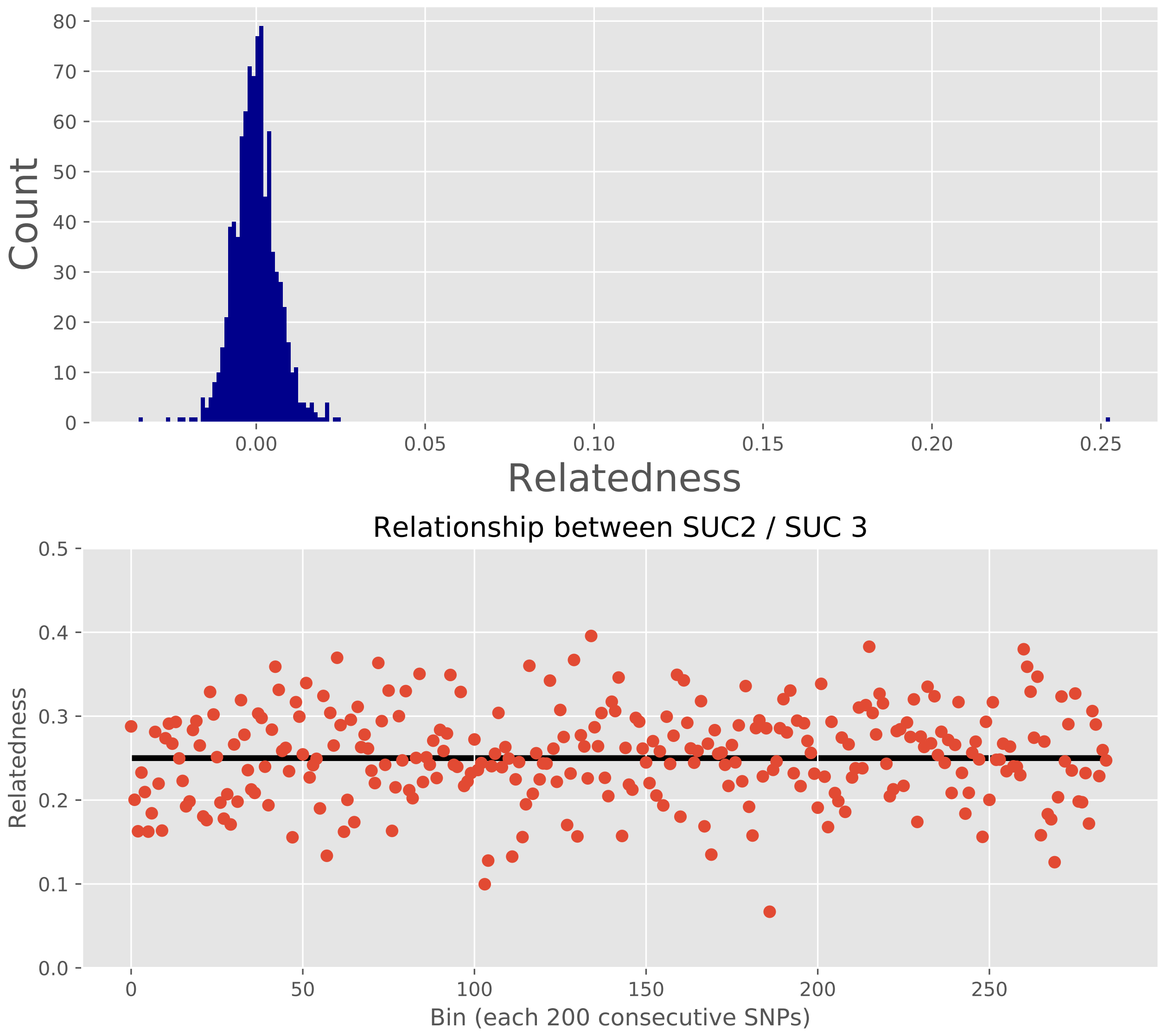


Supplementary Figure 8: **Pairwise Relatedness estimates in ancient Sardinians.** Upper panel: Histogram of all pairwise relatedness estimates for ancient Sardinians, plotted for all pairs with more than $10,000$ intersecting called SNPs. Only a single pair of samples has significantly elevated relatedness (SUC002 and SUC003, single count seen at just larger than 0.25). Lower panel: Estimated $f$ for the two putatively related samples (SUC002 and SUC003), calculated genome-wide using bins of 200 consecutive SNPs ordered along the reference genome.

### 6 Admixture Analysis with qpAdm.

**Harald Ringbauer**

#### Methods

For this analysis we use the software package Admixtools (`https://github.com/DReichLab/AdmixTools`, Version 5.1), which contains the qpAdm program. This tool utilizes the fact that $f$-statistics of an admixed population are a linear mixture of $f$-statistics using putative source population (together called "left" populations) (Haak et al., 2015; Harney et al., 2018).

The rank of a matrix of $f$-statistics with all pairs of a set of outgroup populations ("right" populations) increases when adding a putative source population to the set of left populations, providing a means to assess whether the null model of admixture can be rejected. It is important to stress that a $p$-value from this approach is influenced by sample number and number of SNP markers, and when interpreting results, one must be cautious of the equivalent of Type II errors in the face of limited data.

The results, in particular the $p$-value that describes the model fit, also depend on the set of populations used as an outgroup. To assess the robustness of our results to the outgroup choice, we utilized two groups of populations as out-groups: First a set of 13 outgroup populations from the Human Origins dataset that capture modern human variation ("M13") (Haak et al., 2015). Second, a group of 15 ancient samples ("A15") whose value for disentangling divergent strains of ancestry present in Europe has been previously described in (Lazaridis et al., 2017). If populations from the group "A15" (see table just below) were included in the left populations (WHG and/or Anatolia-N), we removed them from the right set, and used the remaining 14 and/or 13 populations as outgroup (labelled "A14"/"A13").

| Label | Outgroups |
|---|---|
| **M13** | Ami, Biaka, Bougainville, Chukchi, Eskimo, Han, Ju-hoan-North, Karitiana, Mbuti, Papuan, She, Ulchi, Yoruba |
| **A15** | Mota, UstIshim, Kostenki14, GoyetQ116-1, Vestonice16, MA1, ElMiron, Villabruna, WHG, EHG, CHG, Iran-N, Natufian, Levant-N, Anatolia-N |

We found that both sets of outgroups yield qualitatively similar results for our analysis. For conciseness and clarity, in the following we concentrate on the results based on the A15 outgroup, which supposedly reflects finer aspects of ancient West Eurasian genetic variation.

#### Sardinia in relation to mainland Hunter-Gatherer, Early European Farmer, and Steppe ancestries.

We ran a two-way admixture model between Neolithic Anatolians (Anatolia-N) and Hunter Gatherers (WHG) and a three-way model with a Bronze Age Steppe group (Steppe-EMBA) as a proxy for Steppe ancestry. We tested various Neolithic European mainland populations, as well as samples of Sardinia grouped into three time periods (Tab. 2).

| | Source Populations | | | | | Admixture Fractions | | | Standard Error | | |
|---|---|---|---|---|---|---|---|---|---|---|---|
| **Target** | **A** | **B** | **C** | **Outgroup** | **p-Value** | **A** | **B** | **C** | **A** | **B** | **C** |
| Sar-NECA | WHG | Anatolia-N | - | A13 | **0.677** | 0.173 | 0.827 | - | 0.014 | 0.014 | - |
| Sar-EMBA | WHG | Anatolia-N | - | A13 | **0.062** | 0.163 | 0.837 | - | 0.007 | 0.007 | - |
| Sar-Nur | WHG | Anatolia-N | - | A13 | **0.182** | 0.166 | 0.834 | - | 0.009 | 0.009 | - |
| Iberia-EN | WHG | Anatolia-N | - | A13 | **0.414** | 0.080 | 0.920 | - | 0.011 | 0.011 | - |
| France-N | WHG | Anatolia-N | - | A13 | **0.135** | 0.213 | 0.787 | - | 0.015 | 0.015 | - |
| Iberia-ECA | WHG | Anatolia-N | - | A13 | $8.5 \cdot 10^{-6}$ | 0.275 | 0.725 | - | 0.010 | 0.010 | - |
| Iberia-LCA | WHG | Anatolia-N | - | A13 | **0.06** | 0.251 | 0.749 | - | 0.009 | 0.009 | - |
| CE-EN | WHG | Anatolia-N | - | A13 | **0.8** | 0.046 | 0.954 | - | 0.006 | 0.006 | - |
| CE-MNCA | WHG | Anatolia-N | - | A13 | **0.286** | 0.312 | 0.688 | - | 0.009 | 0.009 | - |
| CE-EBA | WHG | Anatolia-N | - | A13 | $< 10^{-30}$ | 0.297 | 0.703 | - | 0.011 | 0.011 | - |
| CE-LBA | WHG | Anatolia-N | - | A13 | $< 10^{-30}$ | 0.296 | 0.704 | - | 0.011 | 0.011 | - |
| Balkans-EN | WHG | Anatolia-N | - | A13 | **0.652** | 0.016 | 0.984 | - | 0.008 | 0.008 | - |
| Balkans-MNCA | WHG | Anatolia-N | - | A13 | **0.673** | 0.068 | 0.932 | - | 0.005 | 0.005 | - |
| Balkans-BA | WHG | Anatolia-N | - | A13 | $< 10^{-30}$ | 0.174 | 0.826 | - | 0.006 | 0.006 | - |
| Sar-NECA | WHG | Anatolia-N | Steppe | A12 | **0.588** | 0.172 | 0.824 | 0.003 | 0.016 | 0.023 | 0.026 |
| Sar-EMBA | WHG | Anatolia-N | Steppe | A12 | **0.041** | 0.163 | 0.837 | 0.000 | 0.009 | 0.012 | 0.014 |
| Sar-Nur | WHG | Anatolia-N | Steppe | A12 | **0.123** | 0.167 | 0.833 | 0.000 | 0.010 | 0.014 | 0.016 |
| Iberia-EN | WHG | Anatolia-N | Steppe | A12 | **0.276** | 0.081 | 0.919 | 0.000 | 0.013 | 0.017 | 0.019 |
| France-N | WHG | Anatolia-N | Steppe | A12 | **0.088** | 0.213 | 0.787 | 0.000 | 0.018 | 0.023 | 0.026 |
| Iberia-ECA | WHG | Anatolia-N | Steppe | A12 | $5.0 \cdot 10^{-6}$ | 0.275 | 0.725 | 0.000 | 0.011 | 0.016 | 0.018 |
| Iberia-LCA | WHG | Anatolia-N | Steppe | A12 | **0.036** | 0.251 | 0.749 | 0.000 | 0.011 | 0.015 | 0.018 |
| Iberia-BA | WHG | Anatolia-N | Steppe | A12 | $6.0 \cdot 10^{-3}$ | 0.239 | 0.689 | 0.072 | 0.010 | 0.014 | 0.016 |
| CE-EN | WHG | Anatolia-N | Steppe | A12 | **0.663** | 0.046 | 0.954 | -0.000 | 0.007 | 0.010 | 0.012 |
| CE-MNCA | WHG | Anatolia-N | Steppe | A12 | **0.247** | 0.308 | 0.679 | 0.013 | 0.011 | 0.015 | 0.018 |
| CE-EBA | WHG | Anatolia-N | Steppe | A12 | $2.3 \cdot 10^{-4}$ | 0.126 | 0.414 | 0.460 | 0.006 | 0.008 | 0.010 |
| CE-LBA | WHG | Anatolia-N | Steppe | A12 | **0.079** | 0.128 | 0.403 | 0.469 | 0.008 | 0.011 | 0.013 |
| Balkans-EN | WHG | Anatolia-N | Steppe | A12 | **0.524** | 0.016 | 0.984 | 0.000 | 0.009 | 0.013 | 0.015 |
| Balkans-MNCA | WHG | Anatolia-N | Steppe | A12 | **0.534** | 0.068 | 0.932 | 0.000 | 0.006 | 0.008 | 0.010 |
| Balkans-BA | WHG | Anatolia-N | Steppe | A12 | **0.13** | 0.117 | 0.703 | 0.180 | 0.006 | 0.009 | 0.011 |

Table 2: **Admixture proportions of ancient European populations inferred by qpAdm.** The upper block of models are two-way models between western hunter gatherer (WHG) and Anatolia Neolithic samples (Anatolia-N) as sources. The lower block of models are three-way models, adding Steppe-EMBA samples as source. (Note: Steppe-EMBA is abbreviated as Steppe in the table). See Sup. Mat. 1G for legend to abbreviations.

#### Continuity on Sardinia across time periods.

We tested a model of continuity within the ancient Sardinians with qpAdm, using A15 as an outgroup. We use the grouping described in the main text: Neolithic and Early Copper Age samples (4500-3000 BCE, $n = 3$), Early Bronze Age (2500-1500 BCE, $n = 19$) and Nuragic (1400-1000 BCE, $n = 17$). We similarly tested a model of continuity from Nuragic into modern samples of the provinces Ogliastra ($n = 419$) and Cagliari ($n = 289$).

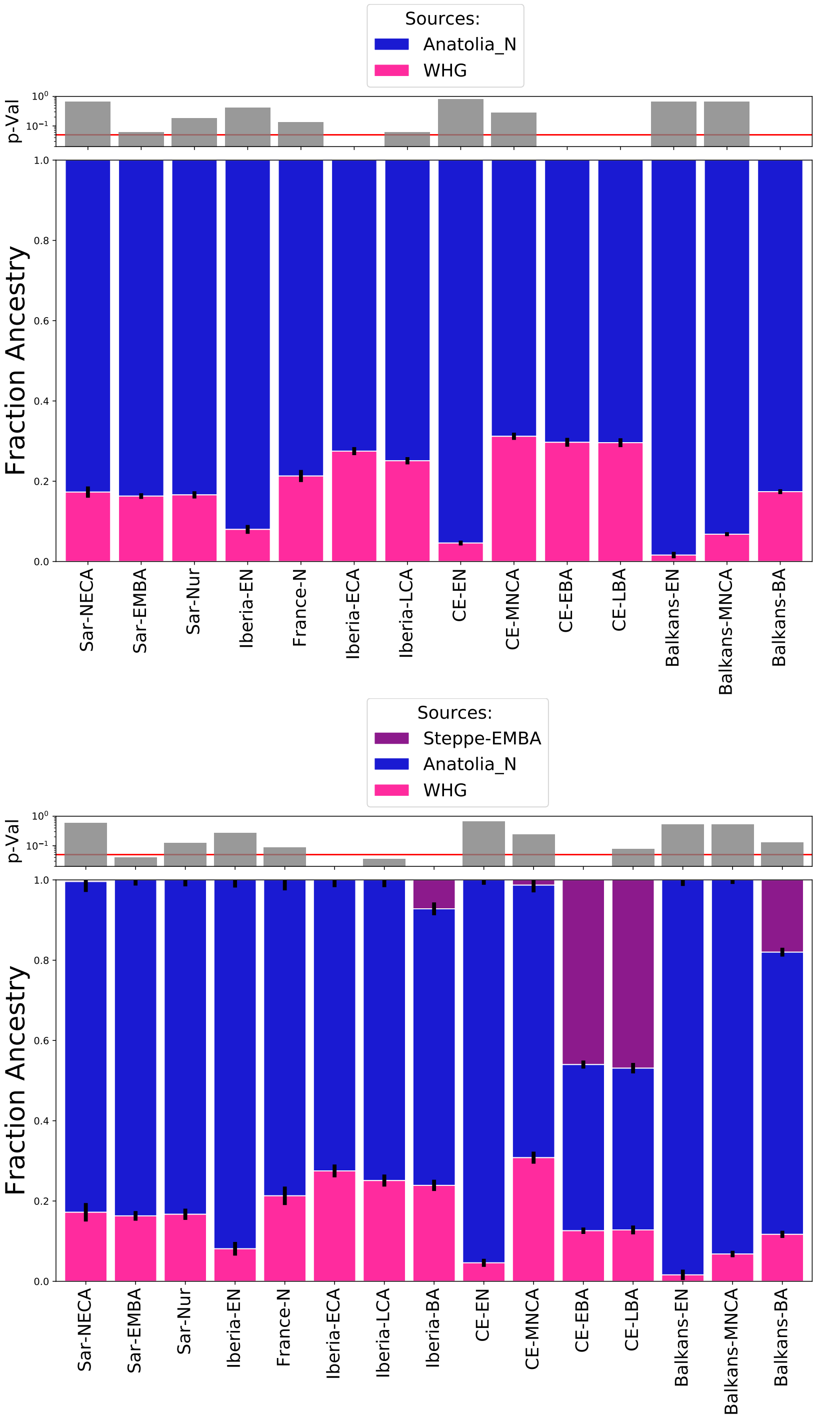


Supplementary Figure 9: **Visualization of two-way (top) and three-way admixture (bottom) of ancient European populations.** Error bars depict standard errors. Upper panel visualizes $p$-values whether this model of admixture can be rejected. The red line depicts a standard significance cut off ($p = 0.05$).

| **Target** | **Source** | **p-value** | **Outgroup** |
|---|---|---|---|
| Sar-NECA | Sar-EMBA | **0.40** | A15 |
| Sar-EMBA | Sar-Nur | **0.13** | A15 |
| Sar-NECA | Sar-Nur | **0.54** | A15 |
| Sar-NECA | Cagliari | $2.8 \cdot 10^{-6}$ | A15 |
| Sar-EMBA | Cagliari | $5.2 \cdot 10^{-43}$ | A15 |
| Sar-Nur | Cagliari | $4.4 \cdot 10^{-43}$ | A15 |
| Sar-Nur | Ogliastra | $3.4 \cdot 10^{-26}$ | A15 |
| Sar-NECA | France-N | **0.074** | A15 |
| Sar-NECA | Iberia-EN | $2.0 \cdot 10^{-8}$ | A15 |
| Sar-NECA | GB-LN | $7.8 \cdot 10^{-10}$ | A15 |
| Sar-NECA | Iberia-LCA | $3.4 \cdot 10^{-12}$ | A15 |
| Sar-NECA | GB-EN | $2.0 \cdot 10^{-13}$ | A15 |
| Sar-NECA | Balkans-MNCA | $8.4 \cdot 10^{-18}$ | A15 |
| Sar-NECA | Balkans-BA | $4.7 \cdot 10^{-18}$ | A15 |
| Sar-NECA | Iberia-ECA | $8.6 \cdot 10^{-19}$ | A15 |
| Sar-NECA | CE-EN | $4.1 \cdot 10^{-34}$ | A15 |
| Sar-NECA | CE-MNCA | $3.2 \cdot 10^{-39}$ | A15 |
| Sar-NECA | Balkans-EN | $1.7 \cdot 10^{-40}$ | A15 |

Table 3: **Support for models of continuity between two populations inferred by qpAdm.** Upper block: Models for continuity between ancient Sardinians. Middle block: Models for continuity between ancient and present-day Sardinia. Lower block: Models of continuity between other ancient populations and ancient Sardinia

#### Admixture of present-day Sardinians.

Next, we fitted present-day Sardinian provinces as an admixture between ancient Sardinians and potential source populations.

##### Two-way admixture models.

Initially, we tested a model of the Cagliari samples as a mix of ancient Sardinians and various source populations to narrow down a plausible model for Sardinian admixture. We chose Cagliari, as PCA analysis revealed that it is a relatively homogeneous population in the center of present-day Sardinian variation (Fig. 2, main text) and because we had a high number of samples from this province ($n = 289$). To maximize the power to reject admixture models, we used the full A15 dataset together as an outgroup.

| | **Source Populations** | | | | | **Admixture Fractions** | | | **Standard Error** | | |
|---|---|---|---|---|---|---|---|---|---|---|---|
| **Target** | **A** | **B** | **C** | **"Right" outgroup** | **p-value** | **A** | **B** | **C** | **A** | **B** | **C** |
| Cag | Sar-Nur | Sicilian | - | A15 | **0.031** | 0.565 | 0.435 | - | 0.021 | 0.021 | - |
| Cag | Sar-Nur | Maltese | - | A15 | **0.013** | 0.590 | 0.410 | - | 0.022 | 0.022 | - |
| Cag | Sar-Nur | Turkish | - | A15 | $8.7 \cdot 10^{-3}$ | 0.715 | 0.285 | - | 0.014 | 0.014 | - |
| Cag | Sar-Nur | Greek | - | A15 | $7.2 \cdot 10^{-4}$ | 0.571 | 0.429 | - | 0.020 | 0.020 | - |
| Cag | Sar-Nur | Tuscan | - | A15 | $2.8 \cdot 10^{-6}$ | 0.540 | 0.460 | - | 0.026 | 0.026 | - |
| Cag | Sar-Nur | Lebanese | - | A15 | $1.2 \cdot 10^{-6}$ | 0.724 | 0.276 | - | 0.016 | 0.016 | - |
| Cag | Sar-Nur | Libyan-Jew | - | A15 | $3.0 \cdot 10^{-7}$ | 0.661 | 0.339 | - | 0.020 | 0.020 | - |
| Cag | Sar-Nur | Cypriot | - | A15 | $1.9 \cdot 10^{-7}$ | 0.662 | 0.338 | - | 0.020 | 0.020 | - |
| Cag | Sar-Nur | Lombardy | - | A15 | $1.1 \cdot 10^{-12}$ | 0.532 | 0.468 | - | 0.031 | 0.031 | - |
| Cag | Sar-Nur | Tunisian | - | A15 | $1.2 \cdot 10^{-23}$ | 0.836 | 0.164 | - | 0.015 | 0.015 | - |
| Cag | Sar-Nur | Spanish | - | A15 | $8.0 \cdot 10^{-26}$ | 0.680 | 0.320 | - | 0.027 | 0.027 | - |
| Cag | Sar-Nur | French | - | A15 | $1.0 \cdot 10^{-27}$ | 0.761 | 0.239 | - | 0.023 | 0.023 | - |
| Cag | Sar-Nur | Basque | - | A15 | $< 10^{-30}$ | 0.747 | 0.253 | - | 0.035 | 0.035 | - |
| Cag | Sar-Nur | Sicily-Bk | - | A15 | **0.43** | 0.535 | 0.465 | - | 0.281 | 0.281 | - |
| Cag | Sar-Nur | Anatolia-BA | - | A15 | $1.8 \cdot 10^{-12}$ | 0.615 | 0.385 | - | 0.058 | 0.058 | - |
| Cag | Sar-Nur | Myc-BA | - | A15 | $6.9 \cdot 10^{-13}$ | 0.303 | 0.697 | - | 0.230 | 0.230 | - |
| Cag | Sar-Nur | Levant-BA | - | A15 | $8.8 \cdot 10^{-21}$ | 0.769 | 0.231 | - | 0.026 | 0.026 | - |
| Cag | Sar-Nur | Steppe-MLBA | - | A15 | $1.6 \cdot 10^{-22}$ | 0.840 | 0.160 | - | 0.015 | 0.015 | - |
| Cag | Sar-Nur | Poland-BA | - | A15 | $2.1 \cdot 10^{-27}$ | 0.807 | 0.193 | - | 0.024 | 0.024 | - |
| Cag | Sar-Nur | Netherlands-BA | - | A15 | $6.2 \cdot 10^{-29}$ | 0.833 | 0.167 | - | 0.020 | 0.020 | - |
| Cag | Sar-Nur | CE-LBA | - | A15 | $5.5 \cdot 10^{-30}$ | 0.808 | 0.192 | - | 0.022 | 0.022 | - |
| Cag | Sar-Nur | GB-LBA | - | A15 | $< 10^{-30}$ | 0.830 | 0.170 | - | 0.021 | 0.021 | - |
| Cag | Sar-Nur | Minoan-BA | - | A15 | $< 10^{-30}$ | 1.000 | 0.000 | - | 0.255 | 0.255 | - |
| Cag | Sar-Nur | Italy-North-Bk | - | A15 | $< 10^{-30}$ | 1.000 | 0.000 | - | 0.406 | 0.406 | - |
| Cag | Sar-Nur | Iberia-BA | - | A15 | $< 10^{-30}$ | 1.000 | 0.000 | - | 0.313 | 0.313 | - |
| Cag | Sar-Nur | Balkans-BA | - | A15 | $< 10^{-30}$ | 1.000 | 0.000 | - | 0.147 | 0.147 | - |

Table 4: Admixture proportions inferred by qpAdm for target Cagliari, with one source set to ancient Nuragic ($n = 15$). Top: Present-day source populations. Bottom: Ancient source populations

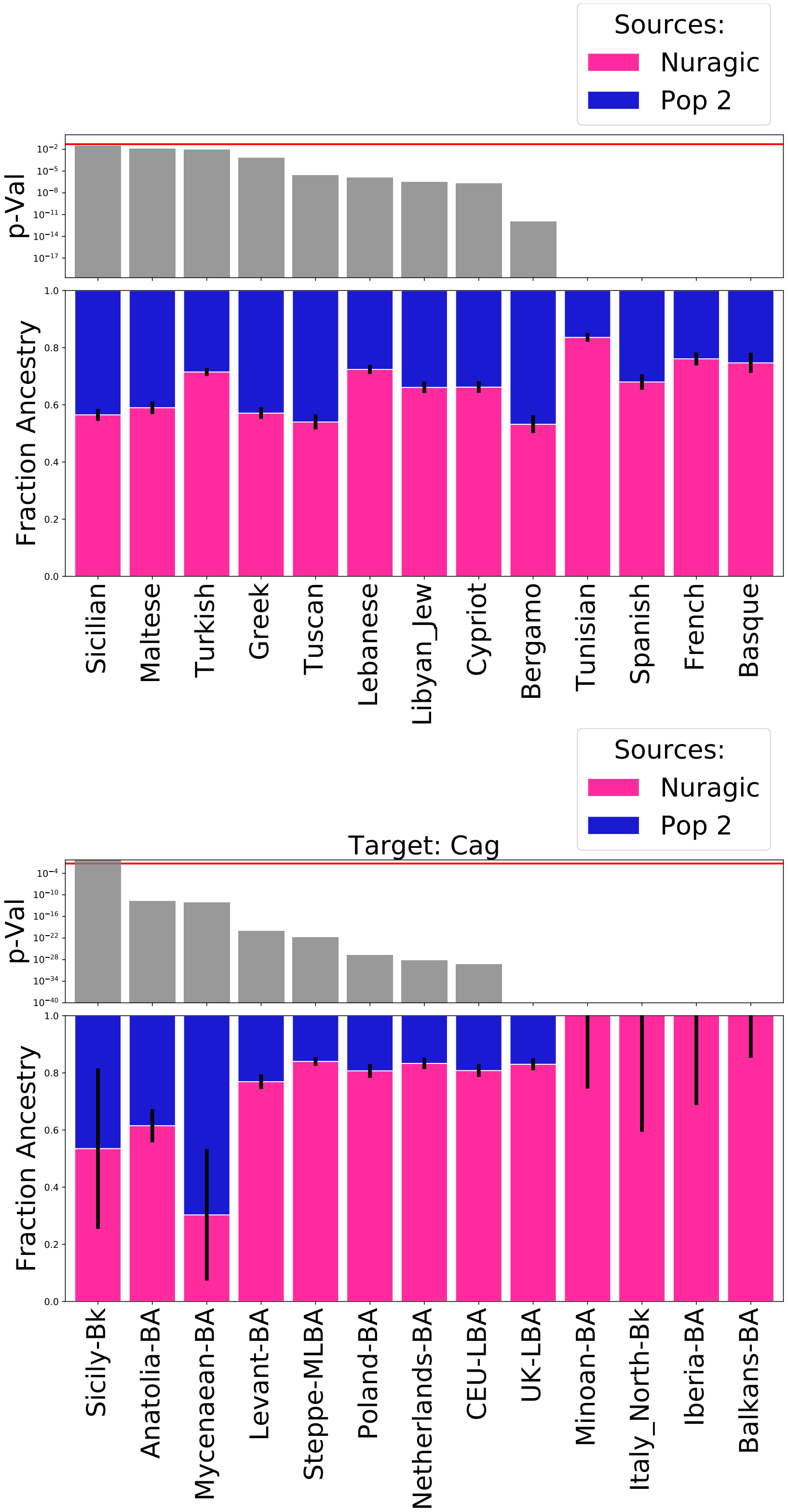


Supplementary Figure 10: **Visualization of various two-way admixture models of $n = 289$ modern Cagliari samples.** We hold one source fixed to Ancient Sardinian Nuragic samples, and tested several putative source populations (x-axis). Error bars depict standard errors. Top: Present-day source populations. Bottom: Ancient source populations.

**Three-way admixture models.**

The two-way admixture model suggested Sicily as the best fitting source for admixture into Sardinia after the Nuragic period ($p = 0.031$ for modern Sicilians, and $p = 0.43$ for Beaker period Sicilian samples "Sicily-Bk", Tab. 4).

Given the history of the region, one possibility is Sicily serves in the two-way model as a proxy for more complex set of ancestries that entered Sardinia after the Nuragic period. To investigate this possibility, we tested potential three-way models that include two present-day populations in addition to the Nuragic Sardinian population as sources. We identified several three-way models that fit as well or even better than Sicily as a single additional source. The best fitting models (of a subset of possible trios we tested) are generally a combination of one population from a set of "northern Mediterranean" population (including Lombardy [ Bergamo], Greek, Tuscan, French) and another population from a set of broadly "eastern Meditarranean" populations (including Lebanese, Turkish and north African Jewish populations). Sicilian and Maltese samples also produce viable three-way admixture models but the estimates of ancestry for the N. Mediterranean component shrink to small values, bringing the fitted parameters in effect towards the two-way mixture models. We interpret this as reflecting that Maltese and Sicilian are a mixture of N. Mediterranean and E. Mediterranean ancestries such that they can serve as single-source proxies in two-way admixture models for a more complex admixture process.

| | **Source Populations** | | | | | **Admixture Fractions** | | | **Standard Error** | | |
|---|---|---|---|---|---|---|---|---|---|---|---|
| **Target** | **A** | **B** | **C** | **Outgroup** | **p-Value** | **A** | **B** | **C** | **A** | **B** | **C** |
| Cag | Sar-Nur | Lombardy | Turkish-Jew | A15 | **0.186** | 0.529 | 0.212 | 0.259 | 0.023 | 0.043 | 0.035 |
| Cag | Sar-Nur | Lombardy | Libyan-Jew | A15 | **0.086** | 0.532 | 0.272 | 0.196 | 0.024 | 0.039 | 0.028 |
| Cag | Sar-Nur | Lombardy | Maltese | A15 | **0.07** | 0.562 | 0.086 | 0.351 | 0.027 | 0.074 | 0.060 |
| Cag | Sar-Nur | Lombardy | Tunisian-Jew | A15 | **0.064** | 0.515 | 0.285 | 0.200 | 0.023 | 0.037 | 0.029 |
| Cag | Sar-Nur | Lombardy | Sicilian | A15 | **0.06** | 0.546 | 0.065 | 0.389 | 0.024 | 0.068 | 0.060 |
| Cag | Sar-Nur | Lombardy | Moroccan-Jew | A15 | **0.05** | 0.536 | 0.247 | 0.217 | 0.024 | 0.044 | 0.033 |
| Cag | Sar-Nur | Lombardy | Lebanese | A15 | **0.049** | 0.571 | 0.269 | 0.160 | 0.026 | 0.040 | 0.023 |
| Cag | Sar-Nur | Lombardy | Druze | A15 | **0.04** | 0.562 | 0.260 | 0.178 | 0.024 | 0.040 | 0.025 |
| Cag | Sar-Nur | Lombardy | Cypriot | A15 | **0.024** | 0.538 | 0.270 | 0.192 | 0.023 | 0.037 | 0.027 |
| Cag | Sar-Nur | Lombardy | Jordanian | A15 | **0.018** | 0.563 | 0.291 | 0.146 | 0.025 | 0.037 | 0.022 |
| Cag | Sar-Nur | Lombardy | Palestinian | A15 | **0.014** | 0.563 | 0.301 | 0.135 | 0.026 | 0.037 | 0.020 |
| Cag | Sar-Nur | Lombardy | Turkish | A15 | $8.6 \cdot 10^{-3}$ | 0.668 | 0.089 | 0.243 | 0.037 | 0.074 | 0.042 |
| Cag | Sar-Nur | Lombardy | BedouinA | A15 | $4.6 \cdot 10^{-3}$ | 0.561 | 0.323 | 0.116 | 0.026 | 0.036 | 0.018 |
| Cag | Sar-Nur | Lombardy | Tunisian | A15 | $1.0 \cdot 10^{-4}$ | 0.547 | 0.375 | 0.079 | 0.028 | 0.034 | 0.014 |
| Cag | Sar-Nur | Turkish-Jew | Lombardy | A15 | **0.186** | 0.529 | 0.259 | 0.212 | 0.023 | 0.035 | 0.043 |
| Cag | Sar-Nur | Turkish-Jew | Greek | A15 | **0.153** | 0.560 | 0.165 | 0.274 | 0.020 | 0.054 | 0.057 |
| Cag | Sar-Nur | Turkish-Jew | Tuscan | A15 | **0.115** | 0.544 | 0.215 | 0.241 | 0.022 | 0.048 | 0.056 |
| Cag | Sar-Nur | Turkish-Jew | French | A15 | **0.094** | 0.573 | 0.321 | 0.105 | 0.021 | 0.028 | 0.024 |
| Cag | Sar-Nur | Turkish-Jew | Basque | A15 | **0.066** | 0.548 | 0.337 | 0.115 | 0.023 | 0.026 | 0.026 |
| Cag | Sar-Nur | Turkish-Jew | Spanish | A15 | **0.052** | 0.555 | 0.309 | 0.137 | 0.022 | 0.030 | 0.032 |
| Cag | Sar-Nur | Turkish-Jew | Turkish | A15 | $5.9 \cdot 10^{-3}$ | 0.688 | 0.068 | 0.243 | 0.049 | 0.158 | 0.112 |

Table 5: Admixture proportions inferred by qpAdm for three-way admixture models. We modeled present-day Sardinian Cagliari with one source fixed to ancient Nuragic Sardinian individuals. Top: northern Mediterranean (proxy: Lombardy ['Bergamo' in HOA data]) as fixed Source 2). Bottom: eastern Mediterranean (proxy: Turkish Jew) as fixed Source 2. Only models with feasible admixture fractions are shown (see also Fig. 11)

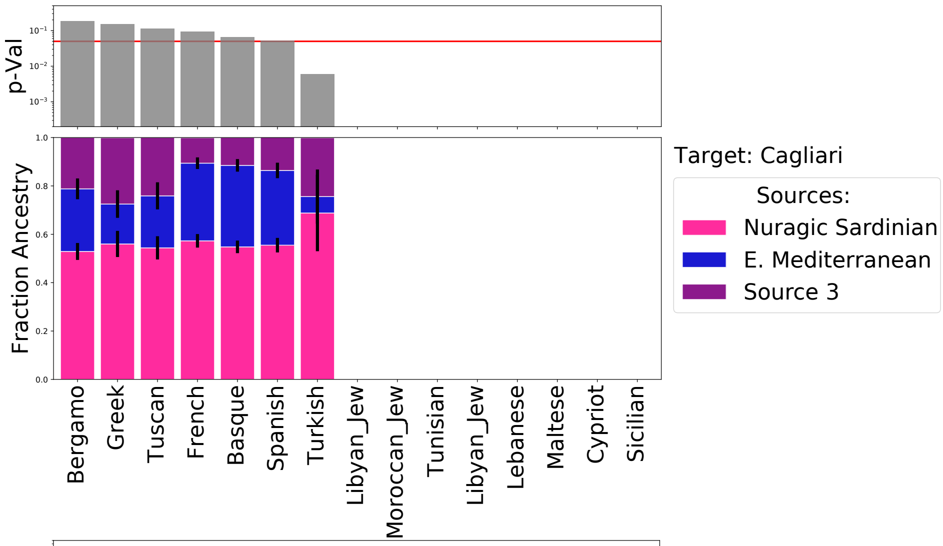


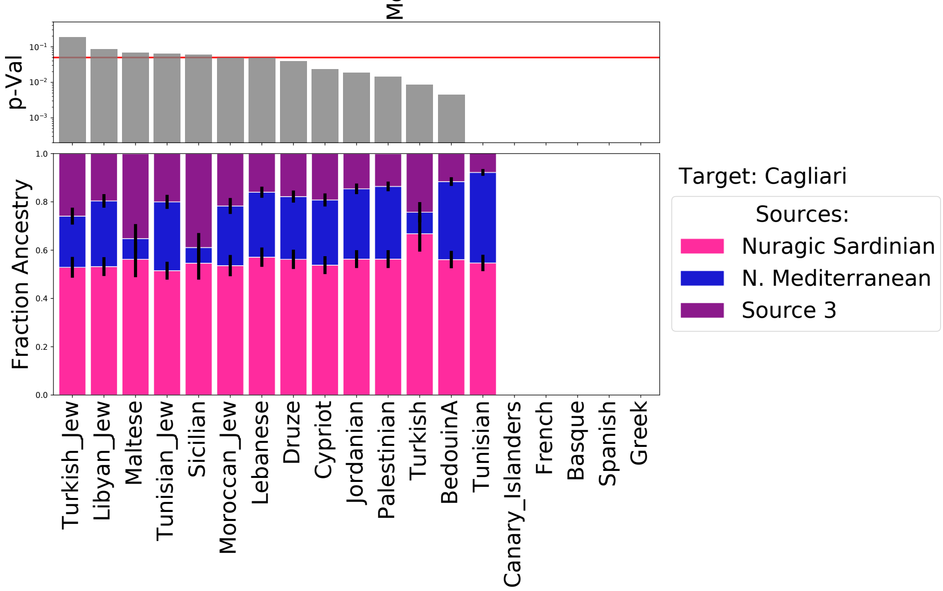


Supplementary Figure 11: : **Visualization of various three-way admixture models of modern Cagliari individuals**. Top: Northern Mediterranean (proxy: Lombardy ['Bergamo' in HOA data]) as fixed source additional to Sardinian Nuragic. Bottom: Eastern Mediterranean (proxy: Turkish Jew) as fixed source additional to Sardinian Nuragic. Error bars depict standard errors. Blank results represent models that did not produce viable results (inferred admixture proportions outside $[0, 1]$).

Using the three-way admixture models, we investigated substructure within Sardinia by fitting all 9 provinces of Sardinia as a three-way admixture between "northern Mediterranean" and "eastern Meditarranean" sources (Supp. Tab. 6 and Supp. Fig. 12). Again, we used A15 as set of right outgroup populations. The results demonstrate geographic substructure within Sardinia with respect to the admixture components. The highest "northern Mediterranean" component is found in the north eastern part of the island (Olbia and to a lesser degree Sassari), while the "eastern Mediterranean" component is highest in the south-western provinces (Carbonia and Medio Campidano). The more isolated provinces, Nuoro and Ogliastra, are inferred to have among the lowest proportions of post-Nuragic admixture.

| | Source Populations | | | | | Admixture Fractions | | | Standard Error | | |
|---|---|---|---|---|---|---|---|---|---|---|---|
| **Target** | **A** | **B** | **C** | **Outgroup** | **p-Value** | **A** | **B** | **C** | **A** | **B** | **C** |
| Cag | Sar-Nur | Tuscan | Lebanese | A15 | **0.058** | 0.576 | 0.298 | 0.126 | 0.026 | 0.046 | 0.029 |
| Car | Sar-Nur | Tuscan | Lebanese | A15 | **0.332** | 0.527 | 0.317 | 0.157 | 0.026 | 0.048 | 0.030 |
| Cam | Sar-Nur | Tuscan | Lebanese | A15 | **0.088** | 0.586 | 0.245 | 0.169 | 0.028 | 0.047 | 0.029 |
| Ori | Sar-Nur | Tuscan | Lebanese | A15 | $8.2 \cdot 10^{-3}$ | 0.614 | 0.261 | 0.125 | 0.029 | 0.053 | 0.032 |
| Ogl | Sar-Nur | Tuscan | Lebanese | A15 | **0.081** | 0.642 | 0.270 | 0.087 | 0.030 | 0.053 | 0.033 |
| Nuo | Sar-Nur | Tuscan | Lebanese | A15 | **0.242** | 0.638 | 0.273 | 0.088 | 0.028 | 0.048 | 0.031 |
| Sas | Sar-Nur | Tuscan | Lebanese | A15 | **0.287** | 0.538 | 0.365 | 0.097 | 0.026 | 0.045 | 0.029 |
| Olb | Sar-Nur | Tuscan | Lebanese | A15 | **0.293** | 0.460 | 0.482 | 0.058 | 0.033 | 0.055 | 0.033 |

Table 6: Admixture proportions inferred by qpAdm for three-way admixture model of present-day Sardinian populations. We fit a three-way model of ancient Sardinian Nuragic ancestry, Eastern Mediterranean ancestry (proxy: Lebanese) and Northern Mediterranean ancestry (proxy: Tuscan)

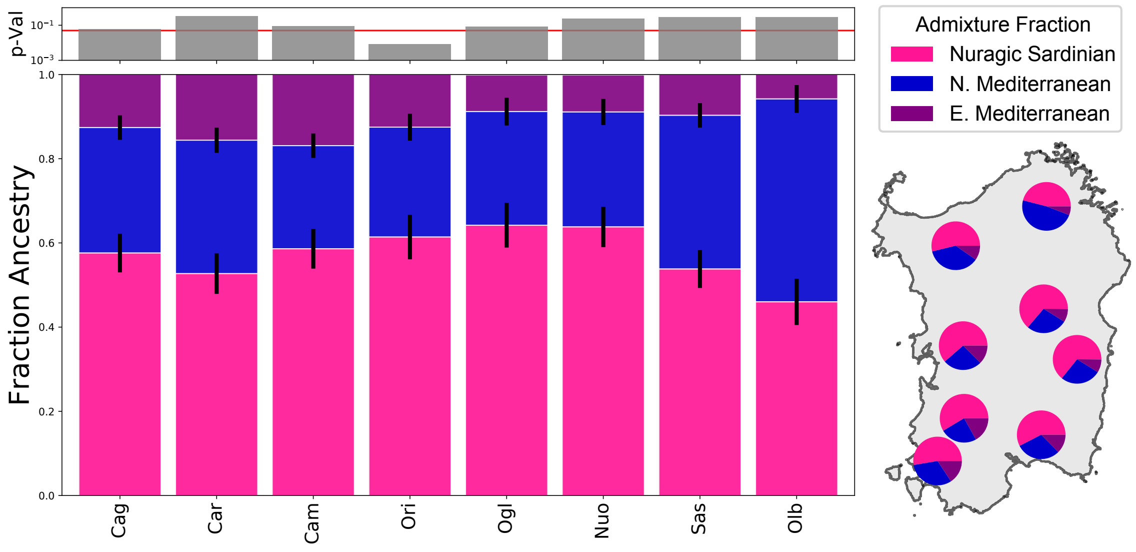


Supplementary Figure 12: : **Visualization of admixture proportions of a three-way admixture model of modern Sardinian populations.** We fit a three-way model of ancient Sardinian Nuragic ancestry, Eastern Mediterranean ancestry (proxy: Lebanese) and Northern Mediterranean ancestry (proxy: Tuscan). Error bars depict standard errors.

To assess the robustness of the three-way model, we tested a four-way admixture model with a proxy for ancient N. African ancestry (pre-colonial Guanche, Canary Islands, $n$ = 5) from Rodríguez-Varela et al. (2017) as an additional source to our best fit three-way model with sources Nuragic Sardinia, eastern Mediterranean (proxy: Lebanese) and northern Mediterranean (proxy: Tuscan). This four-way model does not yield an improved model fit and the Guanche ancestry component is inferred to be very low (Supp. Fig. 13). We additionally tested $n = 5$ Neolithic Moroccans of Fregel et al. (2018) as a fourth source (instead of Guanche), which yielded qualitatively very similar admixture fractions (results not shown).

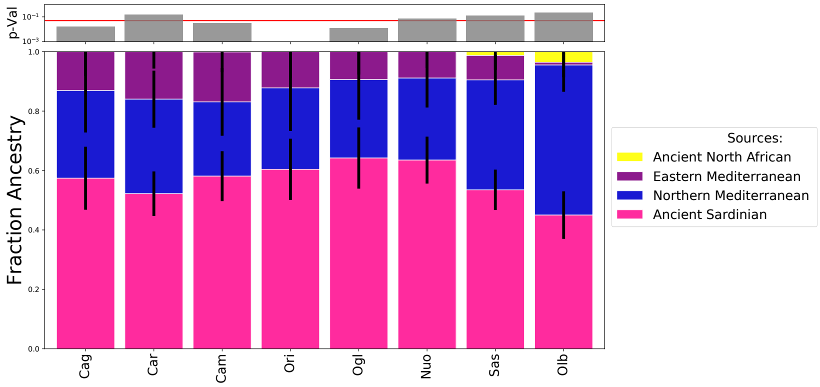


Supplementary Figure 13: : **Robustness of three-way admixture model to inclusion of a fourth ancient north African source.** We fit a four-way model of mixture between ancient Sardinian Nuragic ancestry, Eastern Mediterranean ancestry (proxy: Lebanese), Northern Mediterranean ancestry (proxy: Tuscan), and Ancient North African ancestry (proxy: ancient Guanche, Canary islands). Error bars depict standard errors.

### 7 Population Structure Models

**Joseph H. Marcus and Tyler A. Joseph**

Here we describe extended results applying variants of the Pritchard, Stephens, and Donnelly model (Pritchard et al., 2000) to the dataset of ancient individuals and modern individuals from west Eurasia.

#### 7.1 ADMIXTURE

We applied `ADMIXTURE` to a joint dataset of ancient and modern samples, as described in the Materials and Methods (Alexander et al., 2009). In (Fig. 14) we display a gallery plot, i.e. the typical stacked bar plot for $K = 2$ through 11. For each $K$, we run 5 replicates of `ADMIXTURE` and plot the the runs reaching the highest log likelihood using the interactive visualization tool `pong` (Behr et al., 2016).

#### 7.2 DyStruct

We compared the results from `ADMIXTURE` to a time-aware population structure model: `DyStruct` (Joseph and Pe'er, 2018). `DyStruct` implements a novel variational inference algorithm based on the Pritchard, Stephens, Donnelly model (Pritchard et al., 2000) that incorporates fluctuations in allele frequencies due to differences in sample times. Specifically, `DyStruct` defines a normal approximation to genetic drift that serves as a prior for allele frequency estimates for different time points. At each time point the model is equivalent to the PSD model, but allele frequency estimates between time points are regularized by the prior to ensure allele frequencies estimated from samples nearby in time are closer than allele frequencies from samples further apart. This corrects for genetic drift in populations between samples, potentially leading to different conclusions than `ADMIXTURE`.

We applied `DyStruct` to an un-normalized genotype matrix of ancient and modern samples. Sample times were converted to generation times assuming a 25 year generation time, and provided as input to `DyStruct`. (Supp. Fig. 15) displays the results for $K = 4$. Empirically, `DyStruct` appears to place emphasis on explaining modern populations as mixtures of ancient populations by assigning singular clusters to ancient samples, and describing modern samples as mixtures of these ancient clusters. Hence, ancient samples in `DyStruct` appear as more "extreme" versions of their cluster assignments in `ADMIXTURE`. Consequently, estimates of the genetic contribution from ancient samples into modern populations are different between both models. For instance, modern Sardinian individuals in `DyStruct` appear to inherit a larger fraction of early European farmer ancestry, Steppe/EHG ancestry instead of WHG ancestry, and a smaller portion of shared ancestry from Neolithic Iran and Neolithic Levant.

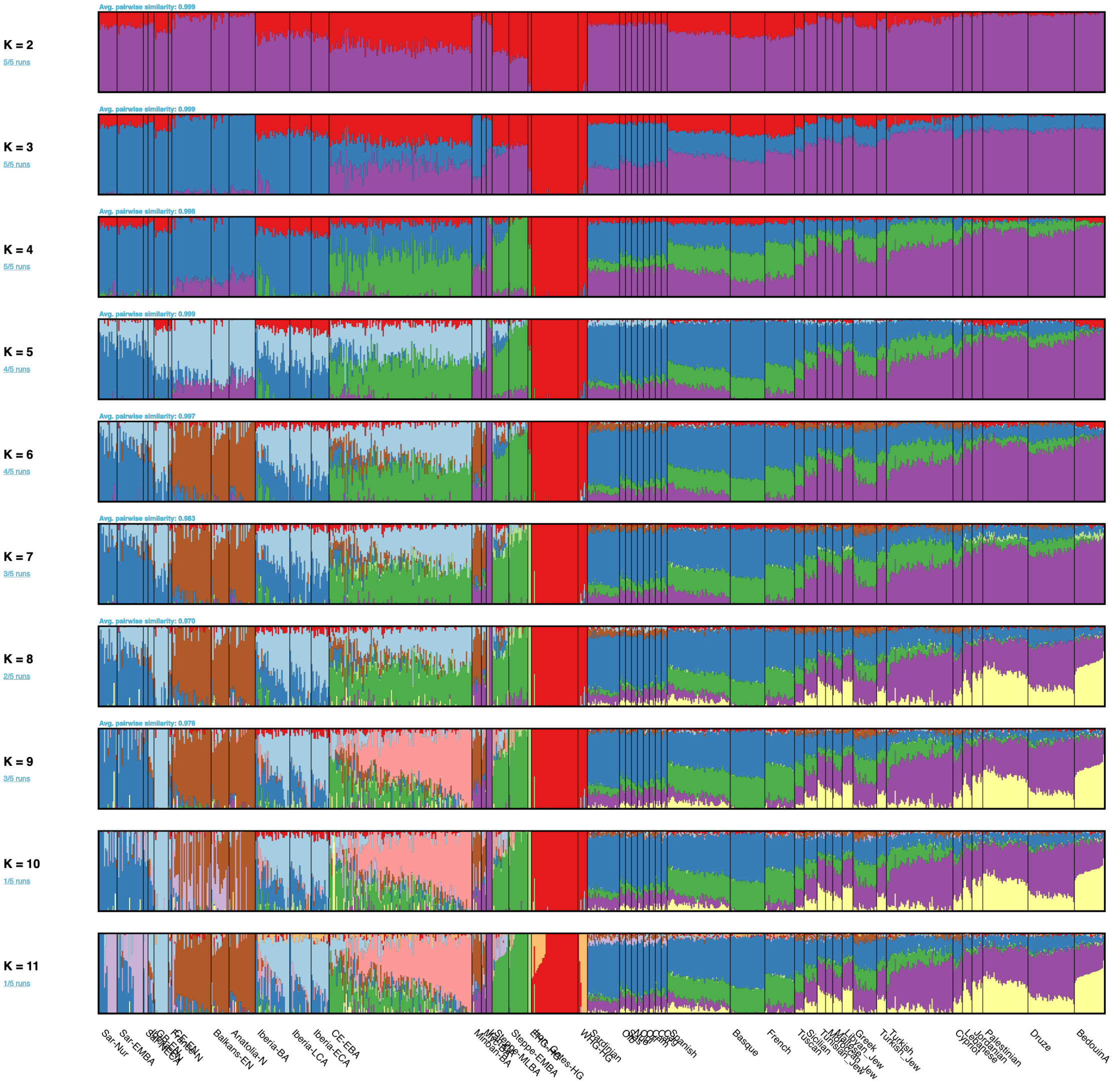


Supplementary Figure 14: **"Gallery" plot of ADMIXTURE solutions for $K = 2, \ldots, 11$ generated with the interactive visualization tool pong.** We depict a subset of both ancient (left side) and present-day populations (right side). We display one solution from a series of 5 replicates for each $K$.

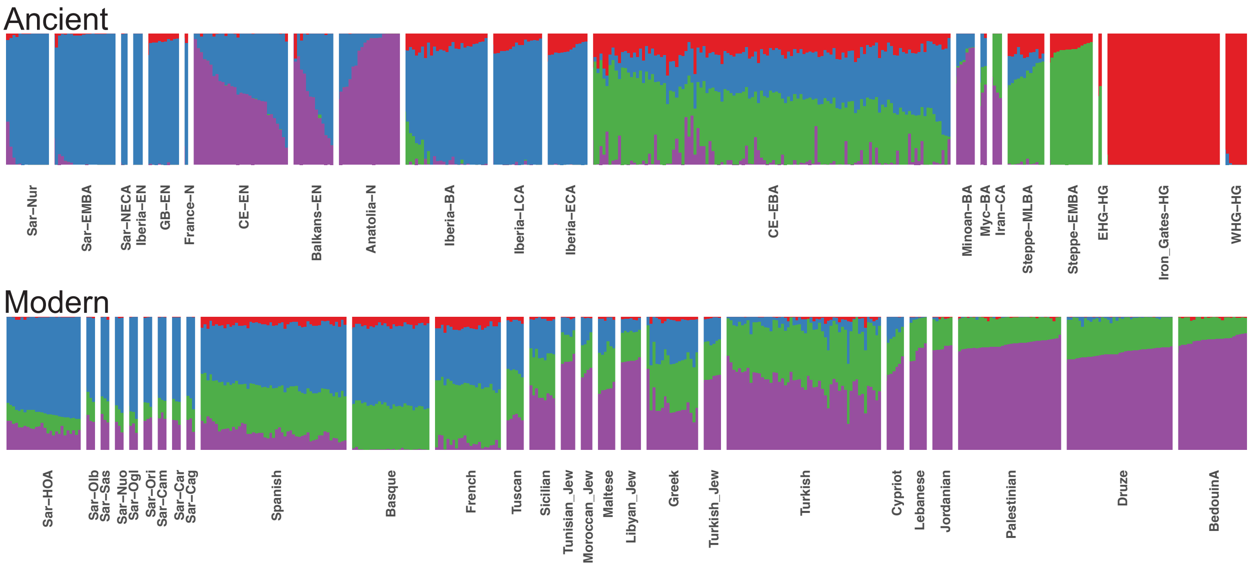


Supplementary Figure 15: **Visualization of admixture coefficients estimated by DyStruct on a subset of the ancient and modern samples analyzed for $K = 4$.** DyStruct identifies three major population clusters in ancient samples consistent with other published literature: early European farmers (blue), Steppe / Eastern Hunter-gatherers (green), and Western Hunter-gatherers (purple). The forth cluster identified (red) corresponds Middle Eastern individuals from Turkey, Armenia, and Iran. The remaining ancient individuals (not pictured) appear as mixtures of these clusters. Consistent with PCA, ancient Sardinian individuals have a strong affinity to a cluster that is putatively defined by early farming populations.

### 8 Shrinkage correction in PC score prediction

**Joseph Marcus**

In Fig. 2 of the main text, we perform principal components analysis (PCA) on contemporary west Eurasian individuals and project each ancient individual onto modern PCs, one at a time, by solving a simple least squares problem. It is known that the estimated principal scores are biased and exhibit a regression towards the mean effect (shrinkage towards 0 if the data is mean centered) for high dimensional data i.e. when the number of features (SNPs) is much greater than the number of samples (individuals) (Lee et al., 2010; Wang et al., 2015; Liu et al., 2018). To correct for this shrinkage effect when predicting PC scores for out of sample individuals, we implemented a shrinkage correction factor through a jackknife re-sampling approach originally proposed by Lee et al. (2010) (for computational experiments see `https://github.com/jhmarcus/pcshrink/blob/master/notebook/patterson-example.ipynb`).

The procedure was performed through the following steps: (1) We compute a rank-$K$ truncated SVD on the full dataset to obtain a first set of uncorrected PC scores. (2) We remove each individual from the dataset and compute a rank-K truncated SVD on the remaining individuals (3) We project the held-out individual on to the PCA computed from the dataset of step (2). Using the eigenvectors computed for each individual, we then constructed a jackknife estimator of the bias. We then applied this correction factor to the ancient individuals' PC scores to create our final visualization. For comparison we also applied two correction procedures, "shrinkmode" and "autoshrink", implemented in `smartpca` (Patterson et al., 2006). In (Supp. Fig. 16) we see no major qualitative differences between the corrected ancient PC scores for the three correction approaches.

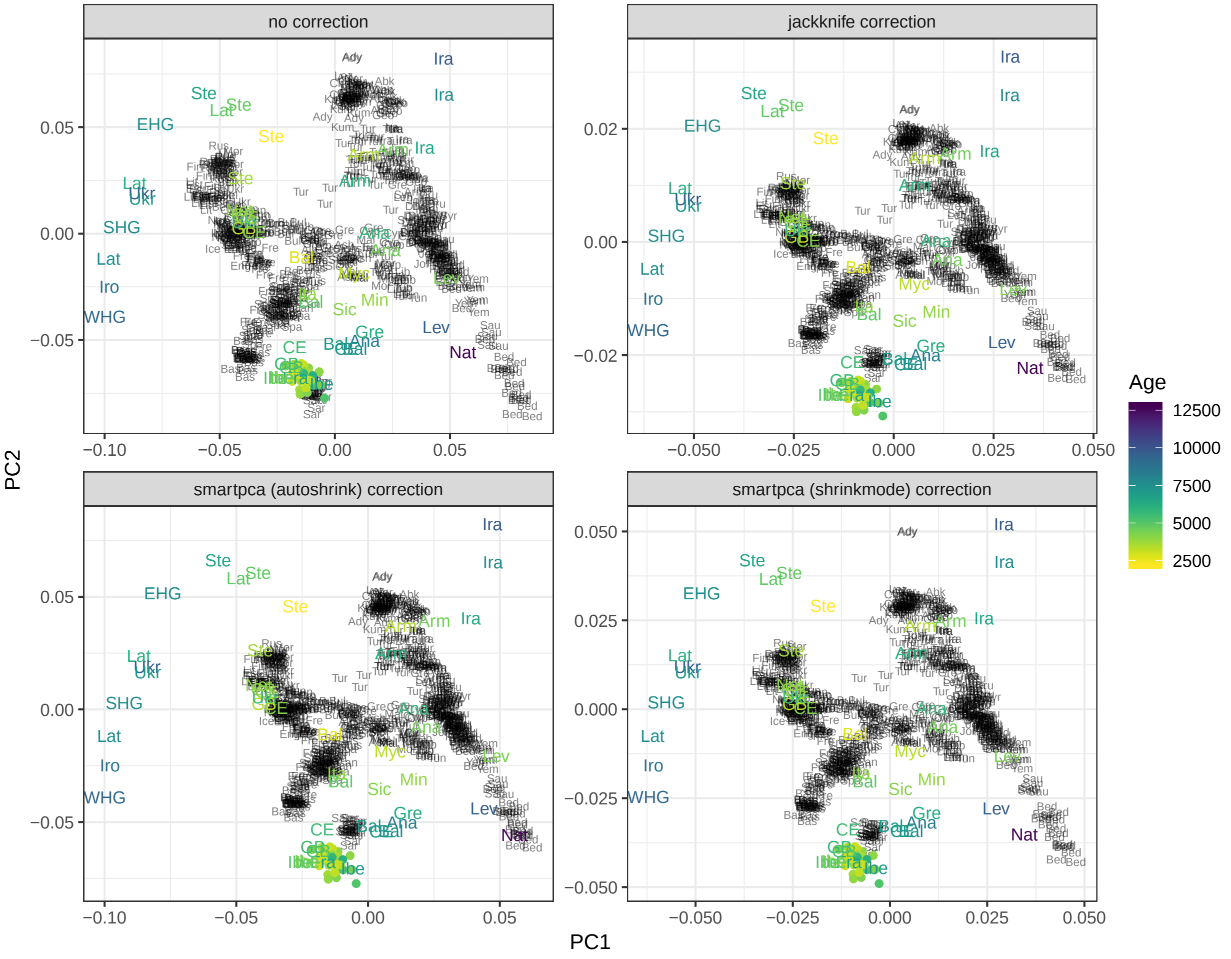


Supplementary Figure 16: **Visualization of the effect of different shrinkage correction approaches on the top two PCs for projected ancient individuals.** All panels show the results an initial PCA on modern Western Eurasian individuals, each of whom are represented as a black three-letter short hand for their assigned population label. We project each ancient individual onto these modern PCs and then represent the median projected PC value of each ancient group as a three-letter short hand colored by the group's median age. Each panel shows a different different correction approach (in the top left showing no correction). We do not observe substantial differences between the three correction approaches especially in the region around the ancient Sardinian individuals from this study.

### References

Hussein Al-Asadi, Kushal Dey, John Novembre, and Matthew Stephens. Inference and visualization of dna damage patterns using a grade of membership model. *bioRxiv*, page 327684, 2018.

David H Alexander, John Novembre, and Kenneth Lange. Fast model-based estimation of ancestry in unrelated individuals. *Genome Research*, 19(9):1655–1664, 2009.

Aaron A Behr, Katherine Z Liu, Gracie Liu-Fang, Priyanka Nakka, and Sohini Ramachandran. pong: Fast analysis and visualization of latent clusters in population genetic data. *Bioinformatics*, 32(18):2817–2823, 2016.

Eugenia D'Atanasio, Beniamino Trombetta, Maria Bonito, Andrea Finocchio, Genny Di Vito, Mara Seghizzi, Rita Romano, Gianluca Russo, Giacomo Maria Paganotti, Elizabeth Watson, et al. The peopling of the last Green Sahara revealed by high-coverage resequencing of trans-saharan patrilineages. *Genome Biology*, 19(1):20, 2018.

Paolo Francalacci, Laura Morelli, Andrea Angius, Riccardo Berutti, Frederic Reinier, Rossano Atzeni, Rosella Pilu, Fabio Busonero, Andrea Maschio, Ilenia Zara, et al. Low-pass DNA sequencing of 1200 Sardinians reconstructs European Y-chromosome phylogeny. *Science*, 341(6145):565–569, 2013.

Rosa Fregel, Fernando L Méndez, Youssef Bokbot, Dimas Martin-Socas, Maria D Camalich-Massieu, Jonathan Santana, Jacob Morales, María C Ávila-Arcos, Peter A Underhill, Beth Shapiro, et al. Ancient genomes from North Africa evidence prehistoric migrations to the Maghreb from both the Levant and Europe. *Proceedings of the National Academy of Sciences*, 115(26):6774–6779, 2018.

Aurelien Ginolhac, Morten Rasmussen, M Thomas P Gilbert, Eske Willerslev, and Ludovic Orlando. mapdamage: testing for damage patterns in ancient DNA sequences. *Bioinformatics*, 27(15):2153–2155, 2011.

Miguel González, Verónica Gomes, Ana Maria López-Parra, António Amorim, Angel Carracedo, Paula Sánchez-Diz, Eduardo Arroyo-Pardo, and Leonor Gusmao. The genetic landscape of Equatorial Guinea and the origin and migration routes of the Y chromosome haplogroup R-V88. *European Journal of Human Genetics*, 21(3):324, 2013.

Wolfgang Haak, Iosif Lazaridis, Nick Patterson, Nadin Rohland, Swapan Mallick, Bastien Llamas, Guido Brandt, Susanne Nordenfelt, Eadaoin Harney, Kristin Stewardson, et al. Massive migration from the steppe was a source for Indo-European languages in Europe. *Nature*, 522(7555):207–211, 2015.

Marc Haber, Massimo Mezzavilla, Anders Bergström, Javier Prado-Martinez, Pille Hallast, Riyadh Saif-Ali, Molham Al-Habori, George Dedoussis, Eleftheria Zeggini, Jason Blue-Smith, et al. Chad genetic diversity reveals an African history marked by multiple Holocene Eurasian migrations. *The American Journal of Human Genetics*, 99(6):1316–1324, 2016.

Éadaoin Harney, Hila May, Dina Shalem, Nadin Rohland, Swapan Mallick, Iosif Lazaridis, Rachel Sarig, Kristin Stewardson, Susanne Nordenfelt, Nick Patterson, et al. Ancient DNA from Chalcolithic Israel reveals the role of population mixture in cultural transformation. *Nature Communications*, 9(1):3336, 2018.

Hákon Jónsson, Aurélien Ginolhac, Mikkel Schubert, Philip LF Johnson, and Ludovic Orlando. mapdamage2. 0: fast approximate bayesian estimates of ancient DNA damage parameters. *Bioinformatics*, 29(13):1682–1684, 2013.

Tyler A Joseph and Itsik Pe'er. Inference of population structure from ancient DNA. In *International Conference on Research in Computational Molecular Biology*, pages 90–104. Springer, 2018.

Iosif Lazaridis, Alissa Mittnik, Nick Patterson, Swapan Mallick, Nadin Rohland, Saskia Pfrengle, Anja Furtwängler, Alexander Peltzer, Cosimo Posth, Andonis Vasilakis, et al. Genetic origins of the Minoans and Mycenaeans. *Nature*, 548(7666):214–218, 2017.

Seunggeun Lee, Fei Zou, and Fred A Wright. Convergence and prediction of principal component scores in high-dimensional settings. *Annals of Statistics*, 38(6):3605, 2010.

Lydia T Liu, Edgar Dobriban, Amit Singer, et al. *e* pca: High dimensional exponential family pca. *The Annals of Applied Statistics*, 12(4):2121–2150, 2018.

Iain Mathieson, Songül Alpaslan-Roodenberg, Cosimo Posth, Anna Szécsényi-Nagy, Nadin Rohland, Swapan Mallick, Iñigo Olalde, Nasreen Broomandkhoshbacht, Francesca Candilio, Olivia Cheronet, et al. The genomic history of southeastern Europe. *Nature*, 555 (7695):197, 2018.

Iñigo Olalde, Selina Brace, Morten E Allentoft, Ian Armit, Kristian Kristiansen, Thomas Booth, Nadin Rohland, Swapan Mallick, Anna Szécsényi-Nagy, Alissa Mittnik, et al. The Beaker phenomenon and the genomic transformation of Northwest Europe. *Nature*, 555 (7695):190, 2018.

Nick Patterson, Alkes L Price, and David Reich. Population structure and eigenanalysis. *PLoS Genetics*, 2(12):e190, 2006.

Jonathan K Pritchard, Matthew Stephens, and Peter Donnelly. Inference of population structure using multilocus genotype data. *Genetics*, 155(2):945–959, 2000.

Ricardo Rodríguez-Varela, Torsten Günther, Maja Krzewińska, Jan Storå, Thomas H Gillingwater, Malcolm MacCallum, Juan Luis Arsuaga, Keith Dobney, Cristina Valdiosera, Mattias Jakobsson, et al. Genomic analyses of Pre-European Conquest human remains from the Canary Islands reveal close affinity to modern North Africans. *Current Biology*, 27(21):3396–3402, 2017.

Pontus Skoglund, Bernd H Northoff, Michael V Shunkov, Anatoli P Derevianko, Svante Pääbo, Johannes Krause, and Mattias Jakobsson. Separating endogenous ancient dna from modern day contamination in a siberian neandertal. *Proceedings of the National Academy of Sciences*, 111(6):2229–2234, 2014.

Chaolong Wang, Xiaowei Zhan, Liming Liang, Gonçalo R Abecasis, and Xihong Lin. Improved ancestry estimation for both genotyping and sequencing data using projection procrustes analysis and genotype imputation. *The American Journal of Human Genetics*, 96(6):926–937, 2015.

João Zilhão. Radiocarbon evidence for maritime pioneer colonization at the origins of farming in west Mediterranean Europe. *Proceedings of the national Academy of Sciences*, 98(24): 14180–14185, 2001.
